# Evolution and adaptation of *Pseudomonas aeruginosa* in the paranasal sinuses of people with cystic fibrosis

**DOI:** 10.1101/2020.10.29.359844

**Authors:** Catherine R. Armbruster, Christopher W. Marshall, Jeffrey A. Melvin, Anna C. Zemke, Arkadiy I. Garber, John Moore, Kelvin Li, Paula F. Zamora, Ian L. Fritz, Christopher Manko, Madison Weaver, Jordan Gaston, Alison Morris, Barbara Methé, Stella E. Lee, Vaughn S. Cooper, Jennifer M. Bomberger

**Affiliations:** Department of Microbiology and Molecular Genetics, University of Pittsburgh School of Medicine, Pittsburgh, PA, USA; Department of Biological Sciences, Marquette University, Milwaukee, WI, USA; Department of Medicine/PACCM, University of Pittsburgh School of Medicine, Pittsburgh, PA, USA; School of Life Sciences, Arizona State University, Tempe, AZ, USA; Department of Otolaryngology, University of Pittsburgh Medical Center, Pittsburgh, PA, USA; Center for Medicine and the Microbiome, University of Pittsburgh and University of Pittsburgh Medical Center, Pittsburgh, PA, USA; Pittsburgh Center for Evolutionary Biology & Medicine, University of Pittsburgh School of Medicine, Pittsburgh, PA, USA

## Abstract

People with the genetic disorder cystic fibrosis (CF) harbor lifelong respiratory infections, with morbidity and mortality frequently linked to chronic lung infections dominated by the opportunistically pathogenic bacterium *Pseudomonas aeruginosa*. During chronic CF lung infections, a single clone of *P. aeruginosa* can persist for decades and dominate end-stage CF lung disease due to its propensity to adaptively evolve to the respiratory environment, a process termed “pathoadaptation”. Chronic rhinosinusitis (CRS), chronic inflammation and infection of the sinonasal space, is highly prevalent in CF and the sinuses may serve as the first site in the respiratory tract to become colonized by bacteria that then proceed to seed lung infections. We identified three evolutionary genetic routes by which *P. aeruginosa* evolves in the sinuses of people with CF, including through the evolution of mutator lineages and proliferative insertion sequences and culminating in early genomic signatures of host-restriction. Our findings raise the question of whether a significant portion of the pathoadaptive phenotypes previously thought to have evolved in response to selective pressures in the CF lungs may have first arisen in the sinuses and underscore the link between sinonasal and lung disease in CF.

**Graphical abstract and highlights:**

- *Pseudomonas aeruginosa* undergoes adaptive evolution in the sinuses of people with CF
- Over time, pathoadapted strains display early signatures of genome degradation consistent with recent host restriction
- Mutations previously thought to occur in CF lungs may have first evolved in sinuses

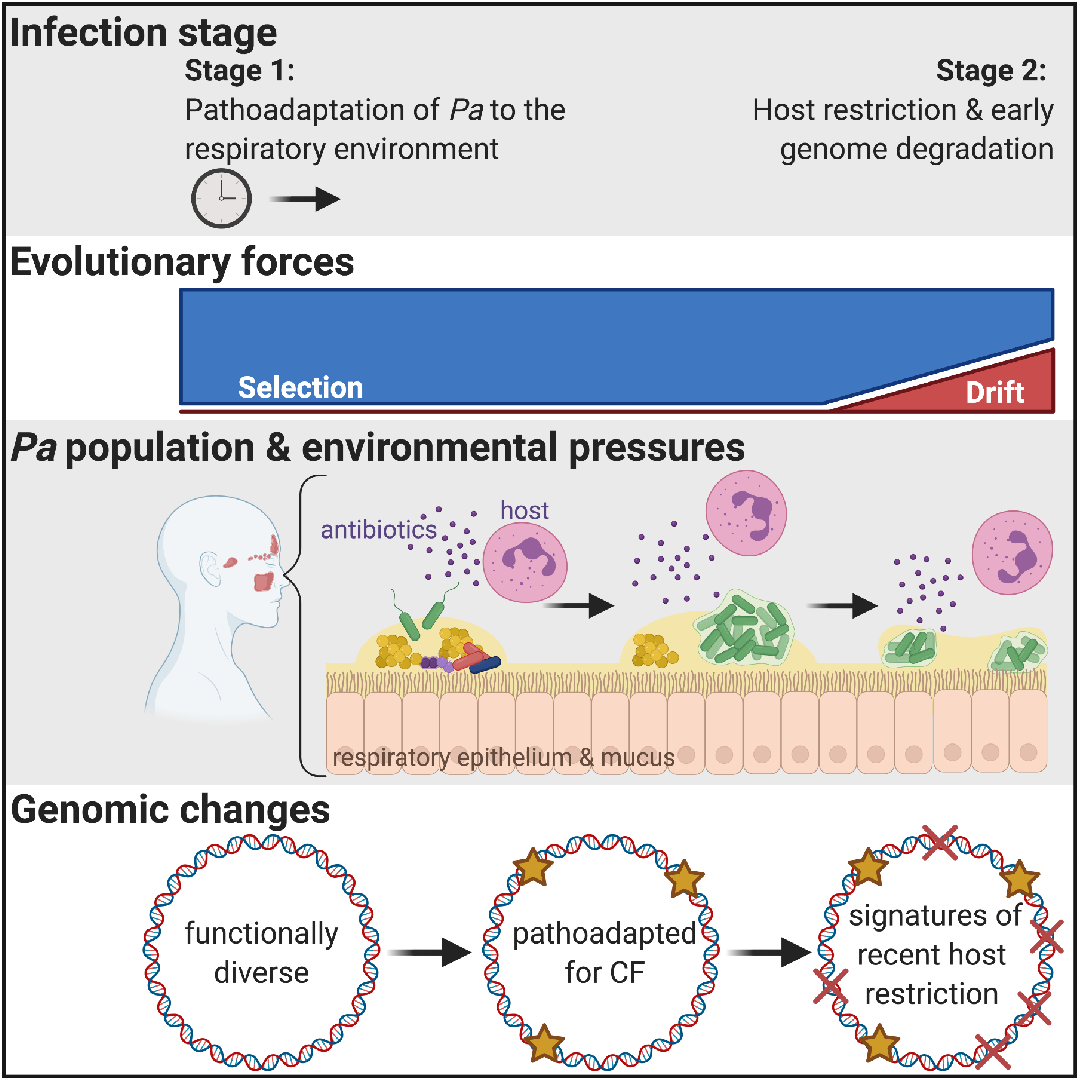

## Introduction

Chronic bacterial infections and the resulting robust but often ineffective host inflammatory response drive the progression of cystic fibrosis (CF) airway disease, causing significant morbidity in people with CF and in some cases, culminating in the need for a lung transplant and/or resulting in mortality due to respiratory failure (*1*). While CF respiratory infections are often polymicrobial, especially early in life (*2*), *Pseudomonas aeruginosa* most commonly dominates end-stage lung disease (*3*). The typical progression of *P. aeruginosa* lung infections in CF includes an intermittent colonization period (typically during childhood), during which sputum is infrequently culture positive for *P. aeruginosa* and antibiotics can reduce the infection to levels that are undetectable clinically. Over time, this progresses to the chronic infection stage, during which sputum is consistently culture positive for *P. aeruginosa* that is no longer responsive to antibiotic therapy (typically during the late teenage years or early adulthood) (*4, 5*). During the intermittent infection stage, people with CF may harbor multiple genetically distinct *P. aeruginosa* strains, but by the time the infection has reached the chronic stage, just one clonal yet phenotypically diverse *P. aeruginosa* population persists (*6*).

Longitudinal studies of *P. aeruginosa*-colonized CF sputum have identified a predictable process in which *P. aeruginosa* adapts to persist in the CF respiratory environment (termed “pathoadaptation”). During the course of pathoadaptation, selection acts on mutations that inactivate genes important for acute infections (e.g. flagellar motility, pyoverdine siderophore production, and Las quorum sensing) and that increase expression of genes promoting chronic infection (e.g. genes that promote biofilm matrix production) (*7, 8*). This accumulation of pathoadaptive mutations is thought to enable a collection of *P. aeruginosa* lineages derived from a single clone to persist for decades in the CF lung environment. Furthermore, pathoadaptive phenotypes have been epidemiologically linked with poor health outcomes: antibiotic resistance and failure to eradicate infection (*9–11*), later stages of disease (*12*), or diminished lung function (*13*) in people with CF. Our goal in studying the selective pressures and evolutionary mechanisms underpinning pathoadaptation in CF is to inform the development of therapeutic strategies that intervene in the pathoadaptation process, thereby interfering with the ability of *P. aeruginosa* to establish a chronic lung infection.

Several studies support a model in which bacteria inhaled from the environment first colonize the sinonasal cavity (referred to as the upper respiratory tract (URT)), with the URT subsequently acting as the main reservoir of bacteria that seeds the lung in CF (*14–24*). Chronic rhinosinusitis (CRS), chronic inflammation and bacterial infection of the paranasal sinuses, is a prominent manifestation of CF in the URT and the prevalence of CRS approaches 100% in people with CF (*25, 26*). The microbiota of CF CRS resembles that of the lung in terms of taxa present and diversity (*27–29*); children harbor comparatively diverse microbes, whereas adults tend to be dominated by one or very few organisms (*2*), with *P. aeruginosa* being a dominant pathogen (*27, 30*). Culture positivity for *P. aeruginosa* in the URT has been observed prior to lung colonization (*18*) and people with CF aspirate sinonasal secretions into their lungs (*31*). Furthermore, concordance in *P. aeruginosa* genotypes has been demonstrated in the sinuses and sputum, and post-transplant lungs become re-infected with the same genotype that was present prior to transplant (*32–34*). Previous studies employed *in vitro* phenotyping and targeted sequencing of genes frequently mutated in CF sputum isolates to show that *P. aeruginosa* isolated from the paranasal sinuses of people with CF acquires phenotypes and mutations previously observed among CF sputum isolates (*19, 32*). Recently, Bartell *et al*. examined pediatric CF sinus *P. aeruginosa* genomes as part of a larger study of evolution in the respiratory tract (*35*), further suggesting similarities between evolutionary paths toward persistence in the upper and lower respiratory tracts. Finally, our laboratory recently characterized an adult CF CRS cohort and found that sinus exacerbation increases the odds of having a pulmonary exacerbation before the next study visit, suggesting a link between sinus and lung disease progression (*36*). Despite this mounting evidence, whole genome-level evidence linking genetic changes within *P. aeruginosa* isolated from the sinuses to known CF *P. aeruginosa* pathoadaptive phenotypes is currently lacking, leaving unanswered the question of how *P. aeruginosa* may pathoadapt to conditions in the URT and whether these enable colonization of the lower airway.

Here, we tested the hypothesis that *P. aeruginosa* adaptively evolves in the sinuses of adults with CF and CRS. We performed the first genome-wide study of sinus *P. aeruginosa* populations in adults with CF and CRS, producing 67 genomes (including 6 polished assemblies using a hybrid assembly approach with short and long-read sequences), and also characterized the sinus microbiome in which *P. aeruginosa* populations evolved. These analyses revealed that the sinuses harbor genetically distinct, clonal populations of *P. aeruginosa* that display the full diversity of pathoadaptive phenotypes previously described among isolates from CF sputum from the lower airway. The evolutionary dynamics of these populations involved three distinct modes, including the evolution of mutators and proliferative mobile genetic elements. Finally, we show that more evolved *P. aeruginosa* lineages display early signatures of genome degradation consistent with recent host-restriction during long term colonization of the sinuses in CF. Our work advances the growing body of literature suggesting that *P. aeruginosa* may first evolve and adapt to the respiratory environment in the sinuses, prior to colonizing the lungs.

## Results

### The sinus microbiome of adults with CF CRS resembles the CF sputum microbiota

The goal of this study was to evaluate the role of the sinuses as a site where *P. aeruginosa* can persist and adaptively evolve before reaching the lower respiratory tract. We performed a longitudinal study of 33 CF adults with symptomatic CRS and prior functional endoscopic sinus surgery to treat their CF CRS (FESS; Table 1) (*36*). During quarterly clinic visits, we obtained endoscopically guided specimens for 16S amplicon sequencing and bacterial culture, and sinus secretions were collected for iron and inflammatory cytokine analyses. Longitudinal sinus microbiome samples were collected from 18 of the 33 participants in order to gain insight into the selective pressures on *P. aeruginosa* in the sinuses. Similar to reports of adult CF sputum (*37–41*) and previous reports of the microbiota of adult CF CRS (*27, 28, 42*), sinus microbiota alpha diversity indices were low (Figure 1A). The median Shannon diversity across all samples from all participants with longitudinal samples was 0.3, with an interquartile range of 0.6. The most common bacterial taxon was Staphylococcus spp., followed by Pseudomonadaceae (based on our culture results, this taxon represents *P. aeruginosa*) (Supplemental figure 1). Because reduced sputum microbiome diversity is correlated with worse lung function in CF (*39, 43, 44*), we next asked whether a similar relationship between microbial community diversity and disease occurs in the sinuses during CF CRS. We measured sinus disease severity through assessment of symptoms with the validated Sino-nasal Outcome Test, SNOT-22(*45*), and with scoring of endoscopic exam findings (modified Lund-Kennedy; mLK)(*46*). We also examined the iron concentration in sinus secretions (a proxy for tissue damage or dysregulated nutritional immunity), and FEV1 (lung function indicator). We found that increased microbial community evenness was associated with decreased mLK score and that increased Simpson’s diversity was predictive of decreased IL-32 levels in sinus washes (p-values < 0.025; multiple linear regression). In summary, a more diverse sinus microbiota, not dominated by one or very few taxa, tended to be associated with less severe sinus disease.

**Figure 1.**
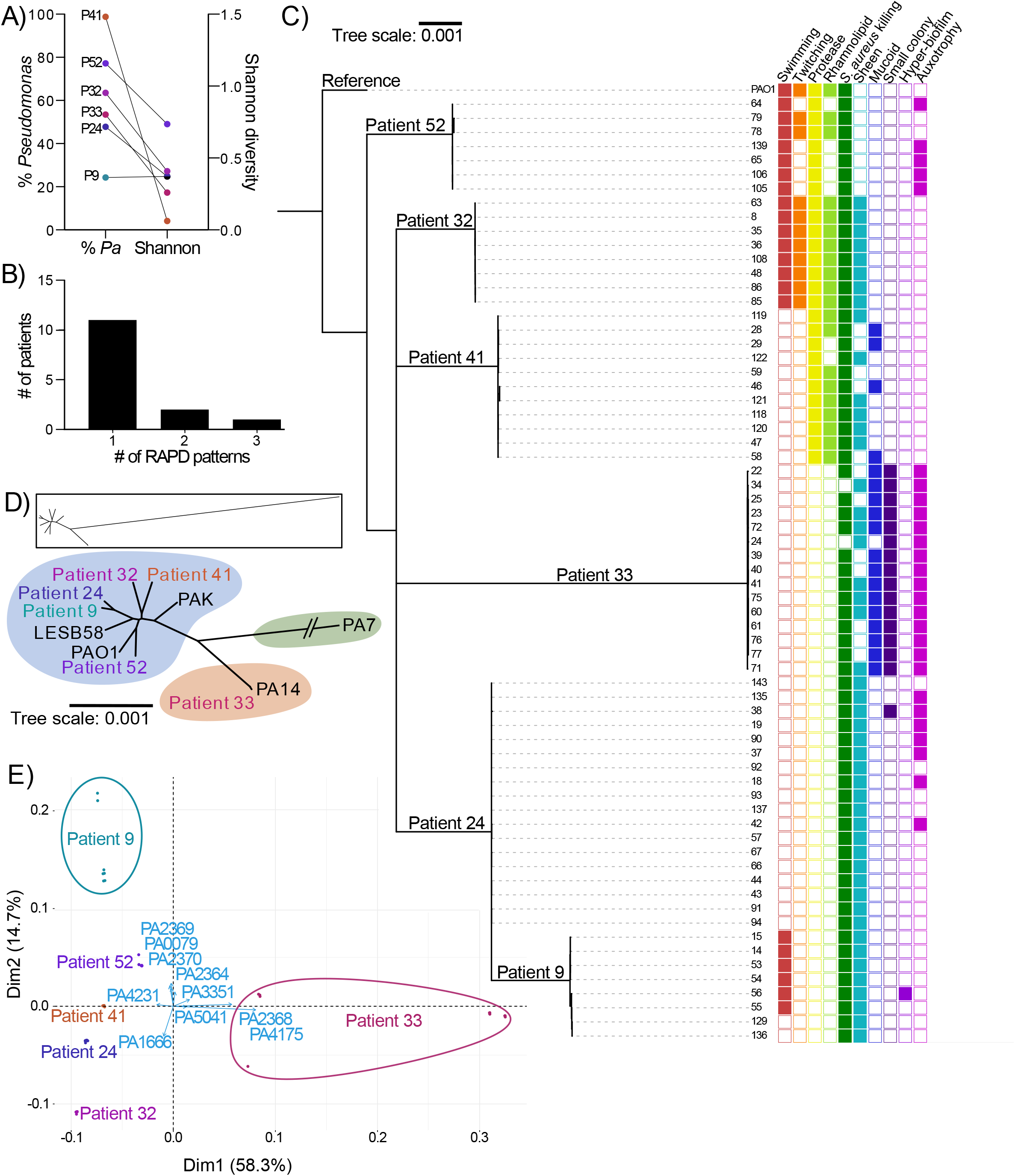
Most study participants harbor a single, clonal lineage of *P. aeruginosa* in their paranasal sinuses, similar to reports of *P. aeruginosa* in CF adults’ lungs. **A)** Average relative abundance of Pseudomonadaceae or *Pseudomonas* spp. (*Pa*) and corresponding average Shannon diversity from 16S amplicon sequencing of sinus swabs, from the 6 study participants who were selected for *P. aeruginosa* whole genome sequencing. The numbers beginning with P on the plot refer to the patient ID numbers in C of this figure. **B)** All isolates collected over the course of the 2 year study were genotyped by Random Amplification of Polymorphic DNA (RAPD) pattern. **C)** Whole genome sequenced sinus isolates display diverse pathoadapted phenotypes, with Patient 33’s lineage appearing the most and Patient 32’s the least pathoadapted. Between study participant genetic diversity of whole genome sequenced *P. aeruginosa* isolates is greater than within an individual person, as expected if each study participant harbored their own clonal lineage. RAxML tree built from core genome SNP distances using all isolates from the six lineages. Reference strain PAO1 is included on the tree. A filled in circle indicates that a particular isolate displays the phenotype (e.g. a filled in circle for “Swimming” means that isolate can swim). Auxotrophy refers to amino acid auxotrophy for all isolates except those in Patient 52’s lineage, in which case it refers to nucleotide auxotrophy. **D)** Sinus *P. aeruginosa* lineages belonged to 2 of the 3 major *P. aeruginosa* clades. The phylogenetic tree depicts relatedness of these 7 strains based on SNPs in the core genome (roary) and was built with RAxML using the GTRGAMMA model with 100 bootstraps. The shaded branches on the tree belong to the three major *P. aeruginosa* clades, as defined by the International *Pseudomonas aeruginosa* Consortium (*47*). Lab strain labels are labeled in black, whereas clinical isolates from this study (the first/reference isolate from the 6 lineages) are colored according to figure 1. With the exception of Patient 33’s lineage which was more closely related to the lab strain PA14, all *P. aeruginosa* lineages were most closely related to the lab strain PAO1 and the Liverpool epidemic strain from CF sputum, LESB58. **E)** Sinus lineages encode a core set of known virulence genes that differ from each other at the single nucleotide level. PCA biplot of dN/dS values (estimated with PAML) for known virulence-associated genes from each whole genome sequenced clinical isolate compared to the distantly-related *P. aeruginosa* reference strain, *P. aeruginosa* PA7. The first two dimensions of the PCA explain 73% of the variance in the core virulence factor dN/dS dataset. Each point on the plot represents a single isolate, with points overlapping if the isolates had identical or nearly identical dN/dS for virulence factors compared to PA7. Large circles were drawn around Patients 9 and 33’s isolates because these isolates showed a high amount of within-lineage variability. The vectors in blue overlaid on the PCA plot represent the top 10 virulence genes that drove differences between isolates. Virulence genes were identified based on annotations of the PAO1 genome available at: http://pseudomonas.com/virulenceFactorEvidence/list. To avoid missing values (which are incompatible with a PCA plot), genes were omitted from this analysis if homologs were not present in all isolates and in PA7. The final plot represents dN/dS values for 236 genes out of the 369 genes annotated as virulence-associated genes in PAO1 from pseudomonas.com.

**Table 1.** Clinical parameters of the 3 cohorts of study participants examined in this study. The “Longitudinal microbiome” cohort includes 18 individuals from the larger 33 person study, for whom we completed longitudinal 16S amplicon sequencing on paranasal sinus swabs. The “Longitudinal *P. aeruginosa* isolates” cohort consists of individuals for whom we had *P. aeruginosa* isolates from at least 2 study visits, with 13 of 14 people also being members of the longitudinal microbiome cohort. The “Whole genome sequenced *P. aeruginosa* cohort” is a subset of 6 people for whom we whole genome sequenced all *P. aeruginosa* isolates and is a subset of the longitudinal *P. aeruginosa* cohort. Continuous variables are reported as median (25-75% percentiles). Age, BMI and ppFEV1 are reported for first visit. Drug use is reported for any time during study. CF-Related diabetes (CFRD) is reported for at any time within the study or +/- 12 months of enrollment, whichever is greater. Clinical parameters of the full patient cohort (n=33) were published in Zemke, IFAR 2019.

|  | Longitudinal microbiome | Longitudinal *P. aeruginosa* isolates | Whole genome sequenced *P. aeruginosa* |
| --- | --- | --- | --- |
| Cohort population (%) | 18/33 (54.5) | 14/33(42.4) | 6/33(18.2) |
| Number of study visits | 6 (4-9) | 6 (4-10) | 9 (8-11) |
| Length of follow-up (months) | 24.2 (20.0-29.4) | 26.0 (21.0-30.3) | 30.0 (19.7-30.5) |
| Male (%) | 5/18 (28) | 4/14 (29) | 1/6 (17) |
| Median Age | 26.8 (24.2-30.7) | 26.8 (24.2-27.7) | 27.3 (26.5-27.6) |
| Median Enrollment BMI (kg/m<sup>2</sup>) | 21.8 (19.5-26.7) | 20.1 (19.5-23.03) | 22.3 (18.45-24.14) |
| CFTR mutation (%) | 10 (56) F508del homozygous<br>7 (39) F508del/other<br>1 (5.5) other/other | 7 (50) F508del homozygous<br>6 (43) F508del/other<br>1 (7.1) other/other | 4 (66.7) F508del homozygous<br>2 (33.3) F508del/other<br>0 other/other |
| CF-related diabetes (%) | 8/18 (44) | 7/14 (50) | 5/6 (83) |
| Baseline percent predicted FEV<sub>1</sub> | 67 (51-83) | 55 (37-67.5) | 63 (32-98) |
| Topical sinus antibiotic use? (%) | 16/18 (89) | 13/14 (93) | 6/6 (100) |
| Inhaled antibiotic use (%) | 18/18 (100) | 14/14 (100) | 6/6 (100) |
| CFTR corrector/modulator use (%) | 9/18 (50) | 5/14 (36) | 2/6 (33) |
| Documented *P. aeruginosa* in sputum during the 12 months prior to enrollment (%) | 11/18 (1 missing, 6 no) | 10/14 (1 missing, 3 no) | 5/6 (1 missing) |

### Adults tend to harbor one or very few clonal lineages of P. aeruginosa during CF CRS

Because the sinuses are proposed to serve as the first site of colonization by *P. aeruginosa* that is inhaled from the environment, we hypothesized that the sinuses of study participants may harbor multiple genetically distinct strains of *P. aeruginosa* that display acute virulence traits. Alternatively, because our study participants were adults and were likely colonized by *P. aeruginosa* for multiple years prior to our study, we reasoned that participants could instead already harbor a single *P. aeruginosa* lineage that displays pathoadaptive traits such as loss of acute virulence factor production and increased biofilm formation. To test these two models, we collected *P. aeruginosa* isolates from the sinuses of 14 study participants (Table 1) on at least 2 separate study visits (n = 101 isolates). After confirming that isolates were *P. aeruginosa* by Sanger sequencing of the 16S gene, we used random amplified polymorphic DNA typing (RAPD) to determine the strain-level genetic relatedness of the isolates. Similar to reports of *P. aeruginosa* isolates from adult CF sputum, most participants were colonized by a single clonal lineage of *P. aeruginosa* in their sinuses over the course of this study (Figure 1B), suggesting that the sinuses can harbor a single strain of *P. aeruginosa* over a time period long enough to support adaptive evolution to the respiratory environment. This finding also refutes our initial hypothesis that the sinuses would be colonized by a succession of environmental isolates.

### Genomic diversity of P. aeruginosa lineages in the sinuses of 6 people with CF

We sought to identify differences between these lineages that are attributable to their evolutionary history prior to the start of our study. We sequenced the genomes of all isolates collected from the six study participants highlighted in Figure 1A (n = 67 genomes; Table 3). Based on RAPD typing, each of these 6 study participants harbored a genetically distinct *P. aeruginosa* lineage in their sinuses. We assembled each isolate genome and built a phylogenetic tree based on single nucleotide polymorphisms (SNPs) within the core genome of all 67 isolates. Isolates from each participant grouped together in a clade (Figure 1C), confirming the RAPD results that *P. aeruginosa* lineages were clonal within these 6 participants.

Based on available sequences of clinical and environmental isolates, the population structure of *P. aeruginosa* is comprised of three major clades, with the lab strains PAO1, LESB58, and PAK belonging to the most common clade, PA14 belonging to the second most common clade, and PA7 belonging to the least commonly sampled, third clade (*47*). Five of the six sequenced lineages belonged to clade one but share no recent history, whereas the lineage from Patient 33 belonged to clade two (Figure 1D). These phylogenetic differences highlight the possibility that variability in *P. aeruginosa* pathogenesis in the CF airway may be explained by the evolutionary history of the infecting *P. aeruginosa* clone. We examined the potential for pre-existing variability in virulence factor production by comparing the ratio of non-synonymous to synonymous variants (dN/dS) for 236 putative virulence genes in all 67 isolates, using the distantly related PA7 strain as the reference for comparison. A principal component analysis (PCA) of this genetic variation in shared virulence traits demonstrated that isolates clustered within study participants’ lineages, which is consistent with independent and unique evolutionary trajectories altering these factors (Figure 1E). The top 10 genes that differentiated isolates were involved in the three *P. aeruginosa* type VI secretion systems that mediate interactions with the host and other bacteria (PA0079/*tssK1*, PA1666/*lip2*, PA2364/*lip3*, PA2368/*hsiF3*, PA2369/*hsiG3*, PA2370/*hsiH3*), flagellar (PA3351/*flgM*) or type IV pili-mediated (PA5041/*pilP*) motility, or nutrient acquisition (the protease PA4175/*piv* and a gene required for pyochelin siderophore biosynthesis, PA4231/*pchA*). Finally, we compared the core and accessory genomes of the six lineages and found diversity in the size of their unique accessory genomes (Figure 2A). Excluding genes for which no function could be predicted, the functional categories (COG) that were most common in the lineages’ unique accessory genomes were genes for mobile genetic elements (especially in Patient 24’s lineage) and for transcription factors (Figure 2B). Each lineage encoded at least two predicted prophages, with Pf1 and phiCTX prophages being the most prevalent (Supplemental table 2). Overall, this sequence-level variability in predicted virulence genes between lineages from CF adults’ sinuses demonstrates that the strength and direction of selection acting on these alleles likely differed prior to the start of our study, either due to differences in the sinus environment between study participants or differences in the environment in which the infecting clone resided prior to entering the sinuses.

**Figure 2.**
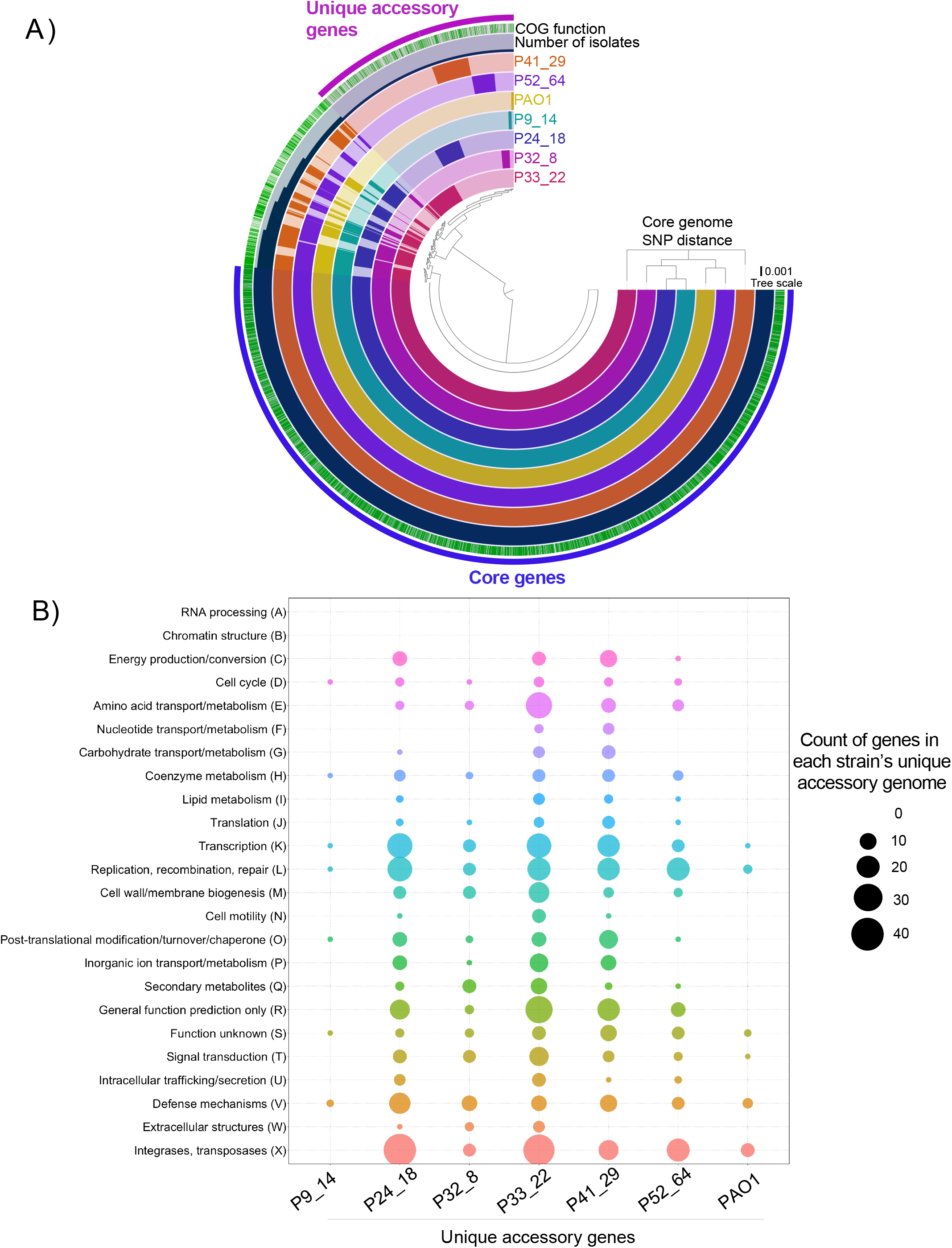
Accessory genomes are enriched with mobile genetic elements, as well as genes involved in surviving and responding to environmental changes. **A)** Gene cluster dendogram of hybrid assemblies from six CF CRS study participants made with anvi-pan-genome in anvi’o version 5.5, with --mcl-inflation 8 and --use-ncbi-blast. The center, circular dendogram depicts gene clusters (ORFs) shared or unique to each of the 6 reference CRS-CF *P. aeruginosa* genomes, plus PAO1. Colored rings from the outside to inside are: Regions of core genes (blue; genes shared in all 7 strains) and Unique accessory genes (pink; genes only present in one isolate), COG function annotation status (green indicates genes that are assigned a COG function), and Number of isolates (dark blue; the number of isolates in which each gene cluster was present). The innermost 7 rings each represent a different hybrid assembled CRS-CF reference isolate or PAO1. The phylogenetic tree on the right depicts relatedness of these 7 strains based on SNPs in the core genome (roary) and was built with RAxML using the GTRGAMMA model with 100 bootstraps and imported into anvi’o using the anvi-import-misc-data function. PAO1 and P9_14 had the fewest unique genes, whereas Patients 24 and 33 had the highest number of unique accessory genes. **B)** Mobile genetic elements make up many of the unique accessory genes that differ between *P. aeruginosa* lineages, especially for P24_18. Plot depicting the count of genes in each COG category, for the unique accessory genome from each strain. In particular, Patient 24 and 33’s unique accessory genomes were enriched for mobile genetic elements, with these two strains having the highest amount of genes in COG categories for Replication, recombination and repair (L) and integrases/transposases (X). Counts of genes that could not be assigned into a COG category are not included on the plot, but are as follows for each genome: P9_14: 19, P24_18: 122, P32_8: 40, P33_22: 233, P41_29: 106, P52_64: 99, PAO1: 7.

### Sinus P. aeruginosa isolates display classic CF pathoadaptive phenotypes that are associated with persistence in the lower respiratory tract

A key aspect of the ability of *P. aeruginosa* to establish chronic lung infections in CF is thought to be its propensity to adapt to the respiratory environment by mutations that diminish or inactivate genes deleterious for persistent colonization. These mutations include those in genes required for flagellar motility, high-affinity siderophore pyoverdine biosynthesis, toxins and proteases, and negative regulators of expression of genes promoting biofilm formation (*6*). However, it is unknown to what extent any of these mutations are selected for in the sinuses, potentially before *P. aeruginosa* colonizes the lower respiratory tract (LRT). To test this model, we asked whether the sinus isolates would display traits more consistent with acute virulence (e.g. motile, producing acute virulence factors) or rather, traits associated with pathoadaptation to the respiratory tract, based on common phenotypes of isolates from sputum of CF adults. Most sinus isolates matched the latter model of pathoadaptation, exhibiting phenotypes such as lysis and sheen (putative LasR variants), increased biofilm extracellular matrix production, and reduced motility (Table 2). Furthermore, most *P. aeruginosa* isolates had lost the ability to produce rhamnolipid and secreted proteases, yet they retained the ability to kill *S. aureus* in an agar plate-based competition assay. Mucoid or rugose colony morphology was less common, with three of 14 participants harboring mucoid isolates and two harboring rugose isolates. Finally, to assess acute toxicity and immune induction, we examined the cytotoxicity of isolates by measuring transepithelial electrical resistance (TEER) and IL-8 secretion following biofilm biogenesis, respectively, in a polarized airway cell model (CFBE41o–) (*48*). Isolates from Patient 32 displayed similar cytotoxicity and induced similar levels of IL-8 secretion as the laboratory strain PAO1, which was isolated from an acute infection (burn wound (*49*)) (Figure 3). Isolates from Patients 9, 24, 41, and 52 displayed intermediate levels of cytotoxicity by 12 hours in the TEER assay, whereas isolates from Patient 33 did not induce an appreciable drop in TEER by this same time point. Based on their interactions with airway epithelial cells and consistent with the *in vitro* phenotyping of pathoadaptive traits described above, we hypothesize that the different *P. aeruginosa* lineages represent a spectrum of pathoadaptation, with Patient 32’s being the least and Patient 33’s lineage the most pathoadapted.

**Figure 3.**
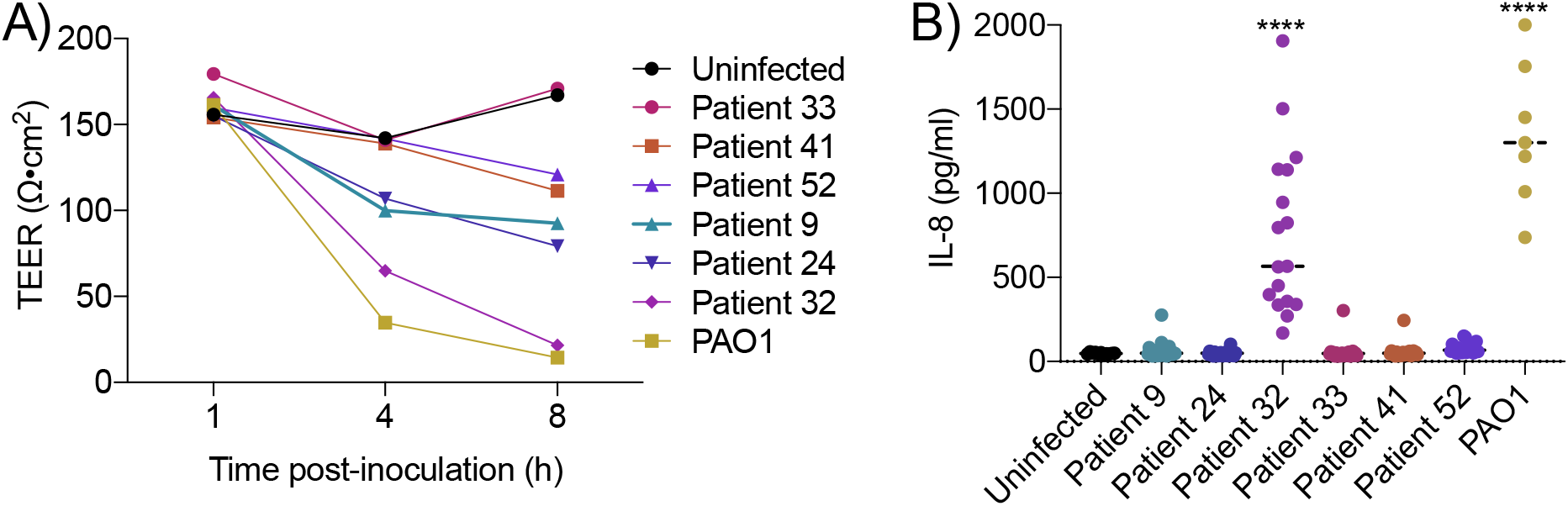
Reduced cytotoxicity of sinus *P. aeruginosa* tracks with level of pathoadaptation. **A)** *P. aeruginosa* lineages display heterogeneity in cytotoxicity, but tend to be less cytotoxic and induce a lesser IL-8 response than PAO1. Isolates from the most phenotypically pathoadapted lineage (Patient 33) are the least cytotoxic. Average trans-epithelial electrical resistance (TEER) of representative early and late isolates from each of the six whole genome sequenced *P. aeruginosa* lineages grown as biofilms on polarized CFBE41o– cells. The laboratory strain PAO1 (an acute skin infection isolate) is represented in black triangles on all plots. With the exception of Patient 32’s *P. aeruginosa* lineage, all CF CRS isolates tested were less virulent than PAO1 (higher TEER values over time). **B)** *P. aeruginosa* lineages display heterogeneity in IL-8 induction that tracks with TEER assay cytotoxicity. An ELISA was performed at the four hour time point during the TEER assay, to measure IL-8 apically secreted by CFBE41o-cells into MEM clear. **** p < 0.0001 by one-way ANOVA with Tukey’s post hoc test for multiple comparisons.

**Table 2.**
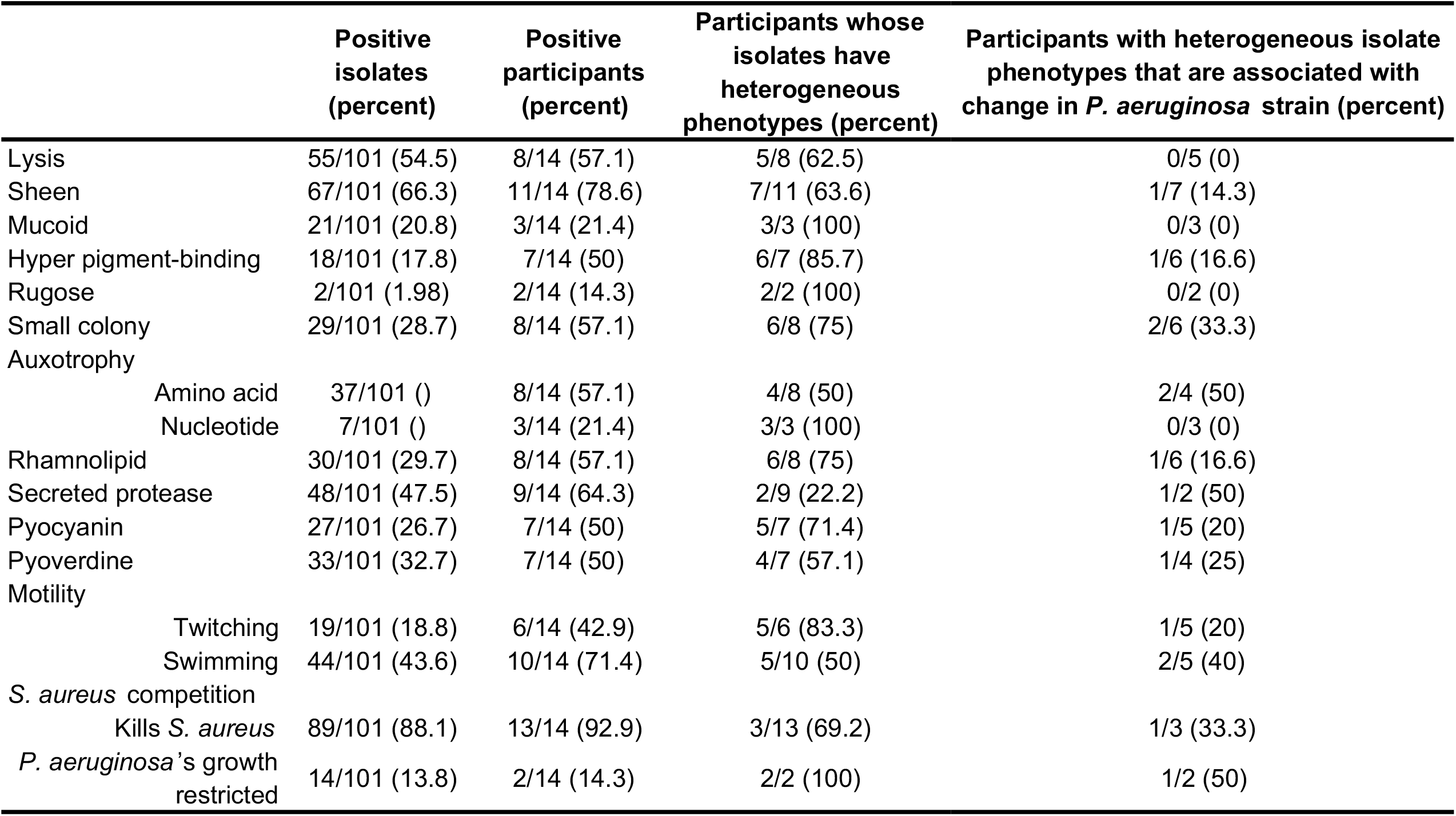
*P. aeruginosa* from the paranasal sinuses of adults with CF display *in vitro* phenotypes commonly observed among evolved lung isolates. Most phenotypes are heterogeneous within study participants. Positive isolates is the total number (percent) of isolates in the study that were positive for that phenotype, across all participants. Positive participants is the number (percent) of study participants who had at least one isolate positive for that phenotype. Heterogeneous phenotypes refers to the number (percent) of participants who had at least one isolate positive and one negative for that phenotype. Changes in *P. aeruginosa* strain refers to the number of participants with a heterogeneous phenotype for whom the change in phenotype was associated with a change in RAPD pattern, with the percent in parentheses. *S. aureus* competition phenotypes are based on an *in vitro* assay in which *P. aeruginosa* was spotted onto a lawn of *S. aureus* lab strain SA113 suspended in 0.3% agar on an LB agar plate. *S. aureus* killing was determined by clearing off the lawn. *P. aeruginosa* growth restriction occurred when a visible *P. aeruginosa* colony or colony borders could not be seen, but *S. aureus* grew as a lawn.

|  | Positive isolates (percent) | Positive participants (percent) | Participants whose isolates have heterogeneous phenotypes (percent) | Participants with heterogeneous isolate phenotypes that are associated with change in *P. aeruginosa* strain (percent) |
| --- | --- | --- | --- | --- |
| Lysis | 55/101 (54.5) | 8/14 (57.1) | 5/8 (62.5) | 0/5 (0) |
| Sheen | 67/101 (66.3) | 11/14 (78.6) | 7/11 (63.6) | 1/7 (14.3) |
| Mucoid | 21/101 (20.8) | 3/14 (21.4) | 3/3 (100) | 0/3 (0) |
| Hyper pigment-binding | 18/101 (17.8) | 7/14 (50) | 6/7 (85.7) | 1/6 (16.6) |
| Rugose | 2/101 (1.98) | 2/14 (14.3) | 2/2 (100) | 0/2 (0) |
| Small colony | 29/101 (28.7) | 8/14 (57.1) | 6/8 (75) | 2/6 (33.3) |
| Auxotrophy |  |  |  |  |
| Amino acid | 37/101 () | 8/14 (57.1) | 4/8 (50) | 2/4 (50) |
| Nucleotide | 7/101 () | 3/14 (21.4) | 3/3 (100) | 0/3 (0) |
| Rhamnolipid | 30/101 (29.7) | 8/14 (57.1) | 6/8 (75) | 1/6 (16.6) |
| Secreted protease | 48/101 (47.5) | 9/14 (64.3) | 2/9 (22.2) | 1/2 (50) |
| Pyocyanin | 27/101 (26.7) | 7/14 (50) | 5/7 (71.4) | 1/5 (20) |
| Pyoverdine | 33/101 (32.7) | 7/14 (50) | 4/7 (57.1) | 1/4 (25) |
| Motility |  |  |  |  |
| Twitching | 19/101 (18.8) | 6/14 (42.9) | 5/6 (83.3) | 1/5 (20) |
| Swimming | 44/101 (43.6) | 10/14 (71.4) | 5/10 (50) | 2/5 (40) |
| *S. aureus* competition |  |  |  |  |
| Kills *S. aureus* | 89/101 (88.1) | 13/14 (92.9) | 3/13 (69.2) | 1/3 (33.3) |
| *P. aeruginosa*'s growth restricted | 14/101 (13.8) | 2/14 (14.3) | 2/2 (100) | 1/2 (50) |

### P. aeruginosa adaptively evolves in the sinus environment during CF CRS

Next, we examined how these clonal lineages of *P. aeruginosa* evolved in the sinuses during the course of our two-year study. We built phylogenetic trees based on core genome SNPs for each of the 6 participants and observed that isolate collection date largely tracked with genetic distance and relatedness, especially in Patients 9 and 33’s lineages, suggesting that sampling of the sinuses during sequential study visits tended to capture newly evolved isolates (Supplemental figure 2). The average number of mutations per year, including non-synonymous, synonymous, indels, and intergenic mutations, ranged from 3 to 95, the upper limit representing the presence of a mutator lineage that has a mutation in the mismatch repair system (Supplemental table 1). In two participants (Patients 9 and 33), the number of shared mutations among subsequent isolates increased over time (Supplemental figure 3), which is consistent with sequential population sweeps driven by selection that lead to fixation. Furthermore, isolates in all six people acquired more non-synonymous mutations than synonymous mutations (Supplemental figure 4). These observations suggest that *P. aeruginosa* is adaptively evolving over time in response to selective pressures in the sinus environment. However, due to the short evolutionary timescale of our study, some non-synonymous mutations that are neutral or weakly deleterious may also have accumulated (*50*). Therefore, the non-synonymous mutations observed in our study are likely a combination of mutations that confer a selective benefit and those that are weakly deleterious, some of which may eventually be removed from the population.

### The sinus environment selects for at least three distinct evolutionary genetic routes to pathoadaptation

We identified three distinct evolutionary-genetic routes that *P. aeruginosa* can take to adapt during prolonged colonization of the paranasal sinuses in CF, including (1) selection on *de novo* nucleotide substitutions (*P. aeruginosa* lineages from Patients 41 and 52), (2) hypermutability caused by defects in DNA repair (Patient 9 and 33’s lineages, and one isolate from Patient 24), and (3) proliferative IS element transposition, a previously underappreciated cause of genetic diversification (Patient 24’s lineage). In contrast, one patient lineage retained acute virulence traits and does not appear to follow a canonical route toward pathoadaptation in CF (Patient 32).

First, the “classic” mode of host adaptation involves selection on mostly nonsynonymous mutations and small insertions or deletions (indels) that recurrently inactivate acute virulence factors and augment traits associated with chronic infection (e.g. biofilm matrix overproduction and antibiotic resistance). These adaptations involve mutations in specific virulence effectors themselves or in global regulatory proteins, which may increase the magnitude of some traits by disrupting negative regulators (*9, 11*). *P. aeruginosa* isolates from the sinuses of Patients 41 and 52 fell into this category of “classic” pathoadaptation, acquiring an average of 7 to 23 total mutations per isolate per year (Supplemental table 1). Based on cytotoxicity (Figure 1CD) and virulence factor production (Table 2 and Figure 1E), these two lineages represent intermediate levels of pathoadaptation.

The second mode involves the evolution of mutator lineages with increased genome-wide mutation rates. One *P. aeruginosa* lineage had already evolved the mutator phenotype prior to the start of our study (in Patient 33) and mutators evolved during our study in two additional study participants’ lineages (from Patients 9 and 24). The two lineages where we recovered the most mutator isolates (Patients 9 and 33) were the same two lineages that displayed a linear increase in fixed mutations over time (Supplemental figure 3), suggesting the mutator phenotype accelerates fixation of mutations in CF *P. aeruginosa* populations. Mutators are most frequently associated with mutational inactivation of *mutS* or *mutL*, leading to defects in mismatch repair (*51–56*). All isolates collected from Patient 33, the most phenotypically pathoadapted lineage, contained a single-base deletion in *mutS* (1319del, 446*). The average rate of mutations accumulation among isolates from Patient 33 was higher than all other lineages, except for the mutator isolates in Patient 9’s lineage (Supplemental table 1).

*P. aeruginosa* isolates were collected from Patient 9 during four study visits, with mutators being isolated on the latter three of the four study days (Figure 4A). The emergence of mutators in this person was also followed by an increased relative abundance of the Pseudomonadaceae taxon relative to *Staphylococcus* spp. (Figure 4B). All four mutator isolates had non-synonymous mutations in *dnaQ* and *dinB*, and three of the four mutators had an additional non-synonymous mutation in *recR*. The *dnaQ* gene encodes the epsilon subunit of DNA polymerase III, the inactivation of which has been associated with a strong mutator phenotype in *E. coli* (*57*). DinB (DNA polymerase IV) is an error-prone polymerase that leads to a mutator phenotype when overexpressed (*58*). Whereas the three non-mutator isolates in Patient 9’s lineage (15, 53, and 54) differed from the reference isolate (P9_14) by 1 to 3 non-synonymous mutations, the mutator isolates each had from 226 to 244 non-synonymous mutations and up to 378 total mutations relative to the first isolate from their respective lineages (Supplemental table 1). Mutator isolates lost acute virulence traits such as swimming motility (Figure 4C) and pyocyanin production (Figure 4D), and had increased biofilm formation (Figure 4E). Isolates from Patient 9 were unique in our study in that all of the latest isolates were mutators, suggesting they outcompeted the early, non-mutators. However, we identified one additional putative mutator isolate with a disrupted DNA repair gene and a relative increase in mutation number from Patient 24.

**Figure 4.**
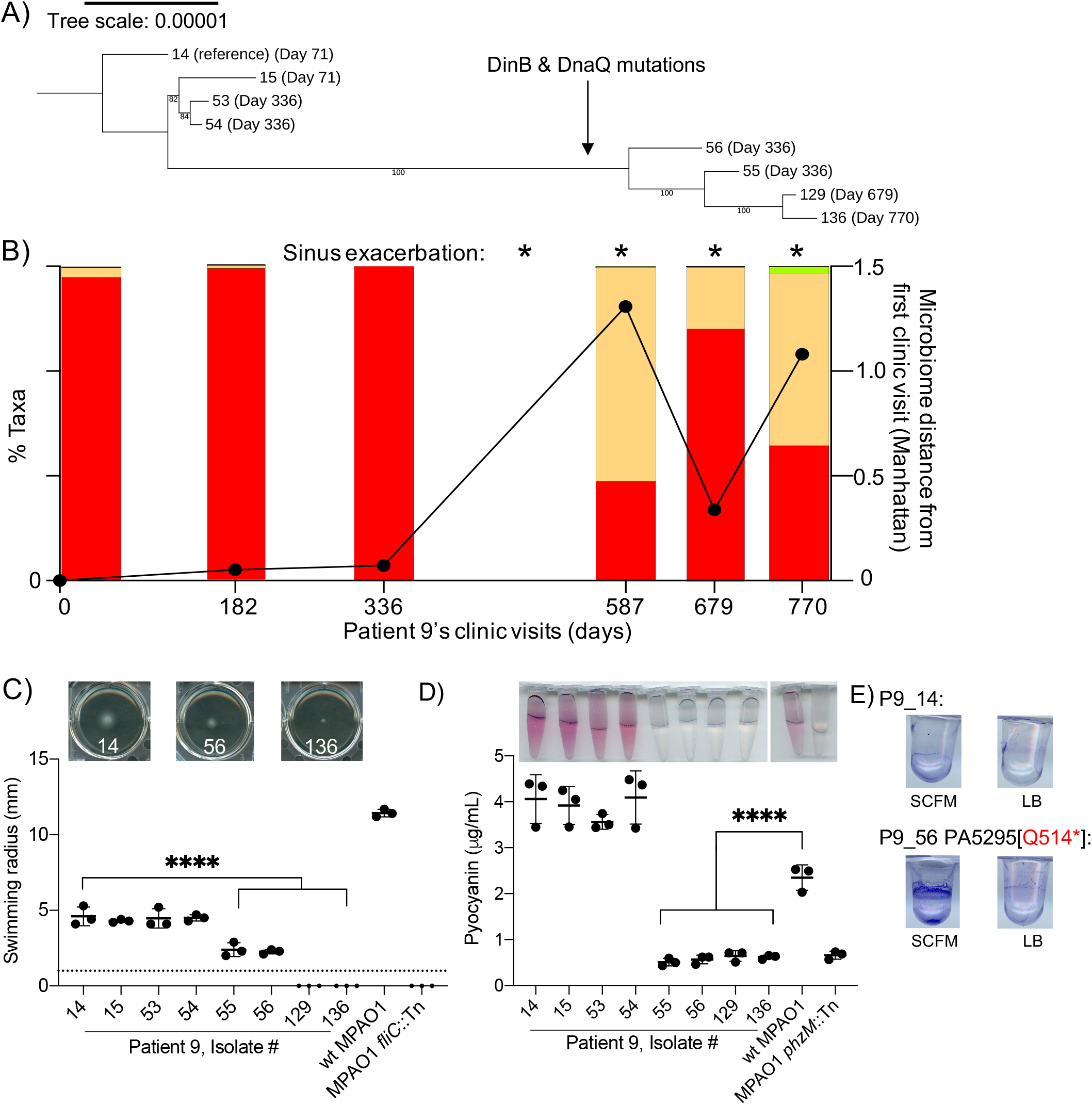
Mutators in Patient 9’s *P. aeruginosa* lineage are associated with shifts in the sinus microbiome, exacerbation, and rapid transition to phenotypes associated with chronic lung infection in CF. **A)** Phylogenetic tree depicting the timing of mutations associated with the mutator phenotype**. B)** Taxa bar plot depicting the timing of emergence of mutator *P. aeruginosa* isolates. Asterisk indicates a visit during which Patient 9 experienced a sinus exacerbation. **C)** Loss of swimming motility in the mutator lineage from Patient 9. Swimming motility was assessed in LB with 0.3% agar after 24 hours at 37°C. The dashed line marks the limit of detection for swimming activity (1mm). Data are from 3 biological replicate experiments. Asterisk = p < 0.0001 relative to the first isolate (one-way ANOVA with post hoc Dunnett’s test for multiple comparisons). Relative to the reference isolate #14, the mutator isolates (55, 56, 129, and 136) all have reduced swimming motility, with complete loss of swimming motility in isolates from the last two time points (129 and 136). Representative swimming assay images above the plot show the gradual loss of swimming motility in Patient 9’s mutator isolates. **D)** Loss of pyocyanin production over time in mutator isolates from Patient 9’s *P. aeruginosa* lineage. Pyocyanin production was extracted and measured after growth in King’s A media, shaking overnight at 37°C. Data are from 3 biological replicate experiments. Asterisk = p < 0.0001 relative to wt MPAO1 (one-way ANOVA with post hoc Dunnett’s test for multiple comparisons). Isolates # 55, 56, 129, and 136 are mutators. MPAO1 *phzM*::Tn is a mutant with a transposon inactivating a gene required for pyocyanin production. Samples in tubes are representative pyocyanin extractions in HCl for each clinical isolate or control. Pyocyanin is pink in 0.1N HCl. **E)** An early stop codon in the hybrid diguanylate cyclase/phosphodiesterase PA5295 in the mutator isolate P9_56 leads to a hyper-biofilm phenotype that is most pronounced in synthetic CF sputum media compared to in LB.

The third mode of pathoadaptation involved the expansion in copy number of insertion sequences (IS elements), which inactivated acute virulence genes in Patient 24’s lineage. Pathogens can adapt to new hosts by mutations produced by transposition of IS elements, which disrupt one or more genes and catalyze genomic rearrangements (*59*). Evolution of antibiotic resistance (*60–63*) and, less frequently, inactivation of acute virulence genes by IS have been reported among *P. aeruginosa* isolates from CF sputum and other human infections (*60*). Unlike in other lineages, most mutations that became fixed in Patient 24’s lineage were insertion or deletion mutations (Supplemental figure 3B). Patient 24’s lineage had the highest number of unique mobile genetic element genes compared to all other sequenced lineages (Figure 3B). Whereas the reference isolates from five other lineages had between 13 and 31 copies of transposase genes, the first isolate from Patient 24 had 92 copies of transposases (Supplemental table 1). Remarkably, these transposases were encoded by IS elements that disrupted many genes that are commonly inactivated by SNPs or indel mutations among *P. aeruginosa* isolated from chronic CF lung infections (Supplemental table 3), especially the IS4 family IS element ISPa45. For example, we observed transposase-mediated single gene disruptions in the first, assembled genome from Patient 24 (P24_18) that persist in all isolates in this lineage. These disrupted genes include the quorum sensing regulator RhlR (PA3477; Figure 5A) and the porin OprD (PA2700) (Supplemental table 3). Consistent with RhlR and OprD inactivation respectively, none of the isolates in Patient 24’s lineage produce rhamnolipid (Figure 1E) and all isolates are resistant to the carbapenem meropenem (Supplemental table 4). Furthermore, the polished assembly of isolate P24_18 largely shares synteny with PAO1, allowing us to identify sites of large chromosomal deletions. Such deletions are often associated with the presence of a transposase at the site of the deletion. For example, all isolates from Patient 24 exhibited lysis and sheen phenotypes when grown on LB agar (Figure 1E), which has been described among CF *P. aeruginosa* sputum isolates as indicative of LasR inactivation and overproduction of the quinolone HHQ (the molecule responsible for the lysis and sheen colony phenotype) (*64*). The likely cause of this phenotype is a 118 gene deletion of the region including LasR/PA1430 relative to the PAO1 genome mediated by the IS4 family transposase ISPa45 (Figure 5B). Finally, over the course of the study, we observed new transposase-mediated deletions in isolates from Patient 24 relative to the first isolate (Supplemental table 3). For example, isolate 38 has a multi-gene deletion spanning the DNA repair protein MutS. Whereas all other isolates have fewer than 15 SNPs or indels relative to P24_18, isolate #38 has 140 SNPs or indels (Figure 5C) likely due to a mutator phenotype caused by the transposase-mediated loss of MutS. IS elements have been previously shown to lead to a mutator phenotype and genomic diversification during long-term experimental evolution of *E. coli* (*65, 66*) and there is one report of a *P. aeruginosa* isolate from CF sputum harboring an IS element (IS6100) that disrupted *mutS* through a large inversion, leading to a mutator phenotype (*67*). Our results suggest that in addition to hypermutation, IS elements may play an important, but as yet understudied, role in accelerating the evolution of pathoadaptive phenotypes in CF.

**Figure 5.**
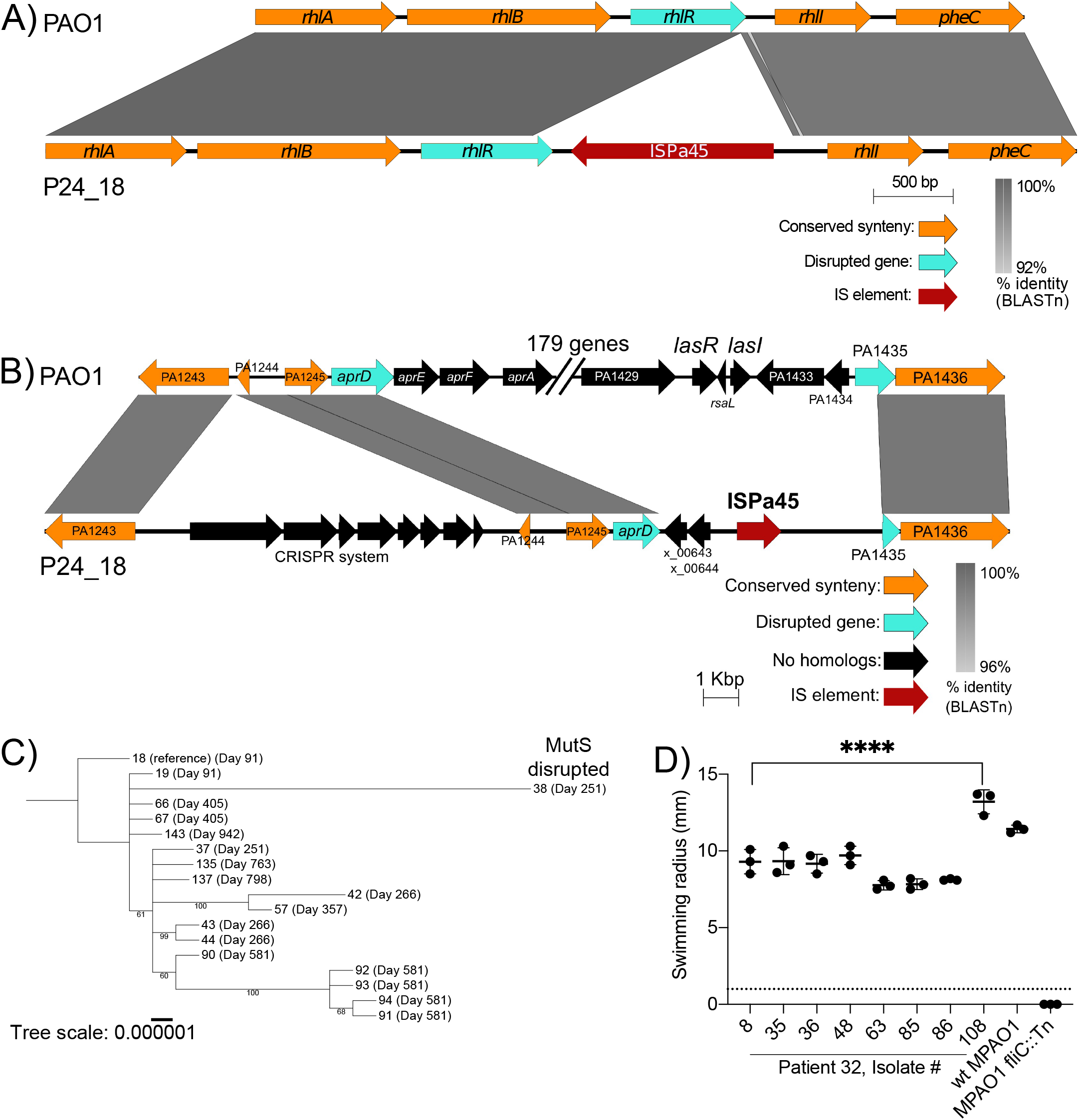
IS elements mediate inactivation of genes involved in acute virulence (including the Rhl and Las quorum sensing systems) and DNA repair. **A)** The transposase ISPa45, an IS element, disrupts the coding region of the transcriptional regulator RhlR in *P. aeruginosa* isolates in Patient 24’s lineage (Reference isolate: P24_18). Open reading frames were aligned by BLASTn using EasyFig version 2.2.2. The top genome is the lab strain PAO1 and the bottom genome is the hybrid-assembled first isolate (reference) from Patient 24’s *P. aeruginosa* lineage. Genes colored orange retained their synteny between PAO1 and P24_18, RhlR (teal) was disrupted by the insertion of an IS element (red) in P24_18. Grey shading indicates the percent identity between PAO1 and P24_18, with a gap in grey shading spanning the inserted sequence. RhlR was disrupted at the 3’ end and a small fragment with homology to the disrupted 3’ end of RhlR is located upstream of ISPa45 in P24_18. **B)** P24_18 has a 188 gene deletion relative to PAO1, including genes required for Las quorum sensing (*lasR*, *lasI*). The two PAO1 homologs flanking this large deletion in P24_18 (*aprD* and PA1435) are both truncated relative to PAO1. In P24_18, the 188 missing genes are replaced by the ISPa45 transposase and two additional predicted ORFs without homologs in PAO1 (x_00643 and x_00644). Additionally, P24_18 has a CRISPR system inserted 3 ORFs upstream of the large deletion. **C)** An IS element (ISPa45) disrupted the DNA repair gene, *mutS*, in isolate # 38 from Patient 24’s lineage, leading to a mutator phenotype as depicted by the long branch length of this isolate on a phylogenetic tree. Phylogenetic tree is based on SNPs in the core genome of Patient 24’s isolates, as determined by roary. The tree was built with RAxML using the GTRGAMMA model with 100 bootstraps and was rooted to the isolate used as the reference for breseq (isolate #18). Dates of isolation are listed next to each isolate. Branches with bootstrap values <50 are collapsed, and bootstrap values > 50 are listed for all branches. **D)** In Patient 32’s lineage, the lineage which displayed the fewest mutations during the study and the fewest pathoadaptive phenotypes, isolate # 108 acquired a single base pair deletion resulting in a frameshift in the flagellar regulator FleS. This mutation resulted in an increase in swimming motility in isolate #108 compared to all other isolates from Patient 32. **** p < 0.0001 by one-way ANOVA with post hoc Dunnett’s test for multiple comparisons.

Finally, one *P. aeruginosa* lineage (Patient 32) retained acute virulence, displays very few pathoadaptive traits, and did not appear to follow any of the three modes of evolution described in the other lineages. Isolates from this lineage uniquely retained levels of cytotoxicity and pro-inflammatory cytokine production similar that of the laboratory strain PAO1 (Figure 1CD). Phenotypically, isolates in this lineage also retained a number of acute virulence traits, including swimming and twitching motility and production of secreted virulence factors such as protease and rhamnolipid (Figure 1E). Additionally, isolates in this lineage were the most susceptible to antibiotics when compared to isolates from the other five lineages, as well as when compared to PAO1 for all antibiotics except the aminoglycosides gentamicin and tobramycin (Supplemental table 4). This lineage acquired the fewest mutations overall (Supplemental table 1), including the fewest mutations in known virulence factor genes (Supplemental table 5). The last isolate from this lineage (P32_108) acquired a single nucleotide deletion resulting in a frameshift in *fleS*, a gene required for flagellar biosynthesis. However, rather than having abrogated production of the flagellum and leading to reduced swimming motility (a pathoadaptive trait), this isolate displays significantly increased swimming relative to earlier isolates from Patient 32’s lineage (Figure 5D). Overall, we found that while *P. aeruginosa* lineages from study participants evolve by different genetic processes in the sinuses, they converge upon more pathoadapted phenotypes, with the exception of Patient 32’s lineage.

### P. aeruginosa lineages exhibit parallel evolution in the sinuses that converges on loss of acute virulence and transition to CF chronic infection phenotypes

*P. aeruginosa* undergoes parallel evolution during chronic CF lung infections and these pathoadaptive mutations promote persistence in the respiratory tract (*6*). Similar to the large airways, the sinuses are lined with a ciliated, mucus-producing, pseudostratified respiratory epithelium that may provide a similar nutritional and physiochemical environment as the LRT (*68*–*71*). Furthermore, many participants in our study received topical treatment with antibiotics in their sinuses. Competition from other microbes represents another potential source of selective pressures, and the microbiota of the sinuses in our cohort also resembles that of the LRT (*27, 28*). Together, the similarities in the respiratory epithelium, antimicrobial exposure, and the microbiota in the URT with what has been described in the LRT suggest that *P. aeruginosa* may be subjected to similar selective pressures in both environments. Therefore, we hypothesized that *P. aeruginosa* adaptively evolves in the paranasal sinuses and that the mutations acquired by sinus isolates would resemble those previously described in isolates from sputum. Furthermore, we hypothesized that pathoadaptive mutations would become fixed in the population because they confer a fitness benefit, for example, due to increased immune evasion and antimicrobial resistance. To test this hypothesis, we assessed whether parallel evolution occurred at the site-, gene-, or pathway-level across any of the *P. aeruginosa* lineages of the six study participants, as has been reported to occur among sputum isolates in CF (*7*). Site-level mutational parallelism occurs when the same site on the same gene is mutated in two or more lineages, gene-level parallelism occurs when the same gene is mutated (but at different sites on the gene), and pathway-level mutational parallelism occurs when different genes within the same pathway are mutated (e.g. genes involved in siderophore biosynthesis, transport, or regulation). Examining whether and how mutational parallelism occurs will help us to understand whether there are common selective pressures in the sinuses, perhaps due to pressure from competing microbes, the host, and antimicrobial treatment, that could prepare *P. aeruginosa* for persistence in the LRT.

First, we examined genes that underwent gene or site-level mutational parallelism (Supplemental table 6) or genes that had 2 or more unique mutations within the same *P. aeruginosa* lineage (Supplemental table 7). Site level mutational parallelism occurred in 5 genes and was attributable to the expansion/contraction of mutations in homopolymers or short tandem repeats in all but one instance (Supplemental table 6). Eleven genes both exhibited parallel evolution across study participants and experienced two or more unique mutations within a single person, suggesting that independent lineages arising within an infection or among different people are subject to common selective pressures in the sinuses. We could confirm that this gene-level parallel evolution was likely due to positive selection and not random chance for some genes. Any gene mutated 4 or more times within or between study participants’ lineages (PA1532/*dnaX*, PA2424/*pvdL*, PA4020/*mpl*) was unlikely to be due to random chance (p = 0.0026; permutation test). Of the 38 genes that were mutated in 2 lineages, 13 were unlikely to be due to random chance, including PA42666/*fusA1* that is known to be mutated in response to selection from aminoglycosides (*72*) (Supplemental table 6; p < 0.03 by permutation test). Notably, while the three lineages containing mutator isolates (Patients 9, 24, and 33) displayed mutations in genes that have previously been described among CF sputum isolates (e.g. *mexXY*, *oprD*, and *wbpL*) and therefore are likely candidates for pathoadaptive mutations, we could not rule out that these mutations were observed due to random chance, given the high number of mutations occurring in these lineages (p > 0.05; permutation test).

We then examined pathway-level mutational parallelism by binning all non-synonymous mutations within coding regions of genes into functional categories (Clusters of Orthologous Groups; COGs). All sequenced lineages acquired mutations in genes predicted to be involved in amino acid transport and metabolism, motility, transcription, and signal transduction (Supplemental figure 5). To better understand whether pathway-level parallel evolution occurred in response to common selective pressures in the CF sinuses, we closely examined three aspects of *P. aeruginosa* physiology that may be under selection due to pressure from competing microbes, the host, and antimicrobial treatment: (1) iron acquisition (Supplemental table 8), (2) the transition from acute virulence (Supplemental table 5) to the biofilm mode of growth associated with less virulent, chronic infections (Supplemental table 9), and (3) antibiotic resistance (Supplemental table 10). There were many instances in which mutations in genes in these pathways could be linked to their expected phenotypes, based on the rich body of literature examining similar mutations in CF lung isolates (Supplemental text). Overall, these findings underscore the similarities in selective pressures present in the URT and LRT that drive the evolution of pathoadaptive mutations in *P. aeruginosa* populations in both respiratory tract sites in CF.

### P. aeruginosa displays early signatures of genome degradation in the sinuses

Genomes of host-restricted bacterial pathogens or symbionts are degraded relative to closely related free-living and non-host-restricted bacteria because of their small population sizes and the increased effect of drift relative to selection (*73*). Two changes that occur rapidly upon host restriction owing to drift are: (1) an increase in gene fragmentation and pseudogene formation and (2) an increase in the number of IS elements present in the genome (*74*). Furthermore, nutritional auxotrophies evolve as genomes degrade and the host-restricted bacterium relies on supplementation by the nutrient-rich host environment, which relaxes purifying selection on prototrophy (*75*). The lineage from Patient 33 displays the most chronic, host-adapted phenotypes of the six whole genome-sequenced lineages in our study, and the lineage of Patient 32 appears the least host-adapted, based on cytotoxicity (Figure 1C,D) and *in vitro* phenotyping (Figure 1E). We predicted that Patient 33’s lineage may display the most signatures of genome degradation, whereas Patient 32’s would have the least. In testing this prediction, we also included two lab strains to serve as references for low and high levels of host adaptation. We expected that the laboratory strain PA14, which was originally isolated from a burn wound (*76*), would display very little evidence of genome degradation and evolution toward host restriction because PA14 likely did not evolve for an extended period of time in the host environment during the infection from which it was isolated. We also examined the LESB58 genome from the Liverpool epidemic strain of *P. aeruginosa* that has been transmitted between people with CF worldwide and that was originally isolated from CF sputum (*77*), which we predicted would display signatures of genome degradation.

We focused on enumerating pseudogenes, which are inactivated remnants of previously intact, functional genes. Pseudogenes commonly form when bacteria undergo an ecological shift that relaxes selection on genes that are no longer required for survival in its new environment (*78*). Inactivated genes are removed relatively quickly from the genomes of free-living bacteria, but are abundant in recently host-restricted bacteria (*79*). We used Pseudofinder to predict pseudogenes (*80*). Consistent with our hypothesis, we found that the first isolate from Patient 33 had the lowest coding density and highest number of predicted pseudogenes, which included fragmented genes and run-on open reading frames (Figure 6A, Supplemental table 1). In contrast, the reference isolate from Patient 32 had the fewest number of pseudogene predictions of all sinus lineages. Both lab strains (PA14 and LESB58) had fewer fragmented and run-on ORFs than any of the six CF CRS lineages. The relatively high pseudogene load among isolates from Patient 33 is consistent with the fact that that lineage displays the most host-adapted and chronic phenotypes, presumably because isolates in this lineage have begun to inactivate and degrade genes no longer required after long-term colonization of the CF respiratory tract. In contrast, the laboratory strain PA14, as well as the epidemic CF strain LESB58, had the fewest number of predicted pseudogenes, presumably because these isolates evolved to retain their ability to live outside the host environment and/or to transmit between people with CF. Isolates in Patient 33’s lineage required amino acid supplementation when grown in a chemically-defined, minimal media, suggesting amino acid auxotrophy had evolved prior to the start of our study (Figure 6B). An additional nutritional auxotrophy, in the form of inactivating mutations in genes required for *de novo* purine biosynthesis, evolved in isolates from Patient 52 (*purH*/PA4854; Figure 6C). Compared to the other CRS *P. aeruginosa* lineages, the reference isolate from Patient 32 had the fewest predicted pseudogenes (Supplemental table 1) and no evidence of defective amino acid or nucleotide biosynthesis, consistent with the fewer mutations in this lineage, further suggesting a more recent colonization in this person.

**Figure 6.**
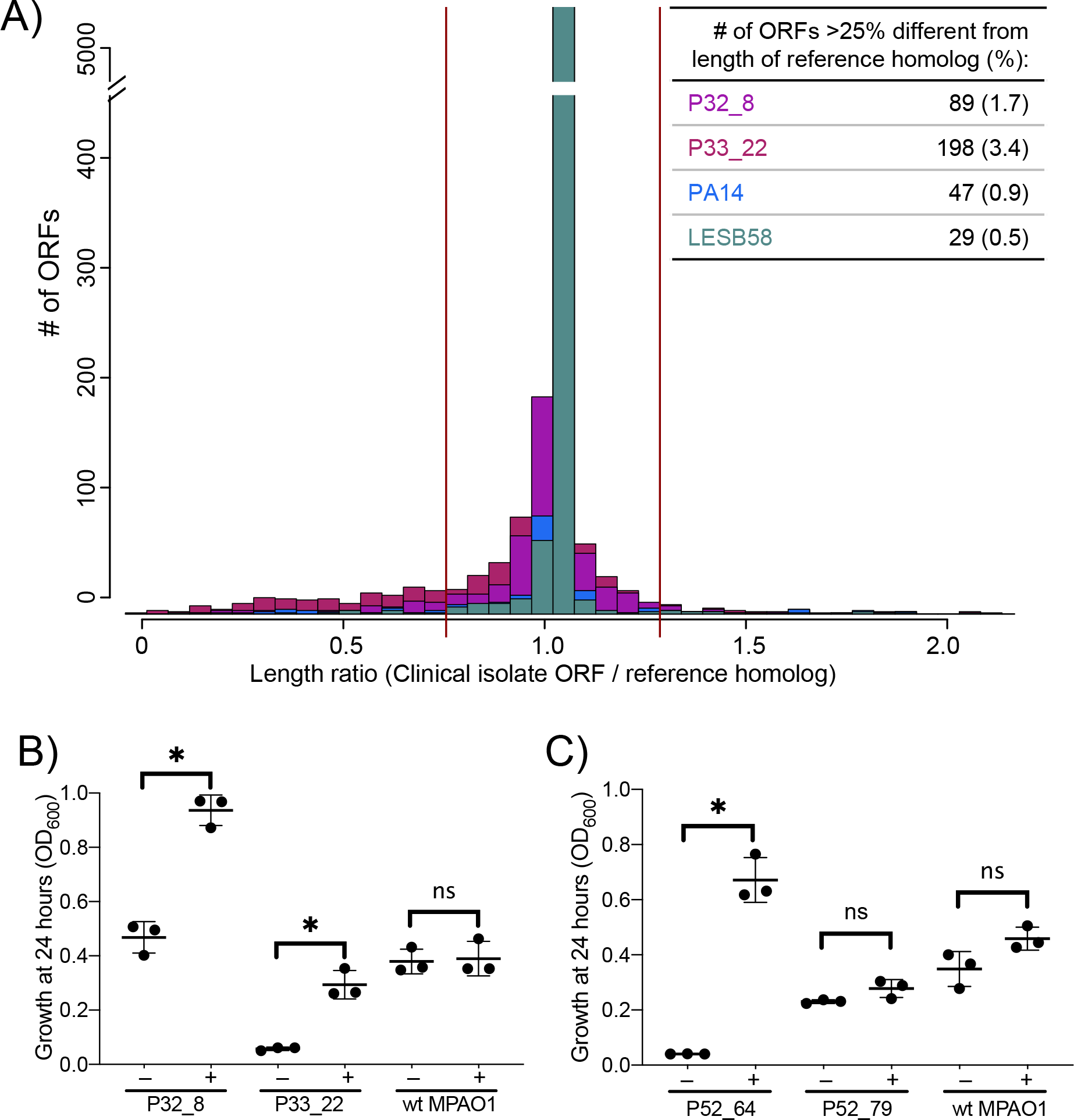
Signatures of host restriction and early genome degradation during adaptation of *P. aeruginosa* to the sinus environment. **A)** Gene fragmentation and run-on ORFs track with degree of pathoadaptation in sinus isolates. Gene fragmentation refers to ORFs that are less than 75% the length of their homolog in a non-CF, non-host adapted lab strain (PAO1 or PA14) and run-on ORFs are greater than 1.25% the length of their homolog. Representative reference isolates from the two sinus lineages with the most (Patient 33/P33_22) and least (Patient 32/P32_8) gene fragmentation compared to the closest related lab strain genome are plotted. ORFs from Patient 33’s isolate were compared to homologs in PA14, whereas ORFs in Patient 32, PA14, and LESB58 were compared to PAO1. Both the lab strain PA14 and the epidemic CF strain LESB58 had very few fragmented genes, suggesting that they are not following an evolutionary path toward host restriction. **B)** Patient 33’s lineage is auxotrophic for biosynthesis of at least one amino acid, whereas Patient 32’s lineage and the lab strain MPAO1 are not. Isolates were grown in M9 media (supplemented with nucleotide precursors, iron, and succinate), with or without 0.5% casamino acids (the + sign indicates the condition where casamino acids were added). *p < 0.05 by one-way ANOVA with post hoc Sidak’s test for multiple comparisons. **C)** Purine auxotrophy in Patient 52’s *P. aeruginosa* lineage. Reference isolate #64 has a SNP in PA4854/*purH,* relative to P52_79 and relative to PAO1, resulting in an amino acid change (Y481S). This mutation results in purine auxotrophy. P52_64 is unable to grow in M9 minimal media plus iron, succinate, and casamino acids, but growth is rescued upon the addition of 2mM adenosine. Plus or minus signs indicate whether adenosine was added to the media. * = p < 0.001; ns = p > 0.05 (one-way ANOVA with post hoc Tukey’s multiple comparisons test).

Finally, during the transition from a free-living to host-restricted lifestyle, bacterial genomes can become enriched with mobile genetic elements, especially transposases that mediate genomic rearrangements and deletions (*73, 81*). The reference isolate from Patient 24’s lineage harbored 92 transposases (Supplemental table 1), including 45 copies of the proliferative IS4 family transposase ISPa45 that inactivated numerous virulence-associated genes and brokered multi-gene deletions in later isolates collected from this person (Supplemental table 3). The IS element density of P24_18 (number of IS elements normalized to genome size in kilobases) was on par with reports of recent obligate host-associated bacteria (IS density: 0.014) (*74*). The IS density of all other CF CRS lineages ranged from 0.002 to 0.005 and select laboratory strains (PAO1, PA14, and LESB58) ranged from 0.001 to 0.002. These genomic adaptations suggest that early stages of degenerative genome evolution likely occurs as *P. aeruginosa* persists in the respiratory tract of people with CF.

## Discussion

When opportunistic pathogens like *P. aeruginosa* colonize susceptible hosts, the population experiences strong selection by the prevailing factors at the colonization site. For people with CF, this site is thought to be the upper respiratory tract, which shares some features of the lungs that ultimately become infected but is also distinct. An overarching question in the field of CF microbiology is whether the process of adaptive evolution by *P. aeruginosa,* which poses the greatest threat to people with CF, begins in the sinuses and sets the stage for chronic lung infection. Many people with CF suffer from chronic rhinosinusitis (CRS) and these infections could predispose spread to and persistence in the lower airway. We studied a longitudinal collection of *P. aeruginosa* from the sinuses of 14 adults with CF CRS and discovered that sinus populations display pathoadaptive phenotypes and mutations long described to occur in CF lung populations (*6*). The genomes of these isolates enabled inference of the population genetic processes shaping these infections and delineated three distinct routes by which *P. aeruginosa* evolves in the sinuses during CF CRS. Yet these processes converged on selection to amplify a common set of traits and functional specialization caused by the beginnings of genome degradation. This genetic decay was accelerated by high mutation rates, which were caused by defects in mismatch repair or rampant increases in IS transposition in some lineages, and enabled by genetic drift accelerated by population bottlenecks and effects of selective sweeps on linked, favored mutations (*82*). Overall, our study strongly suggests that some portion of the mutations present in lung *P. aeruginosa* populations may have emerged first in the upper respiratory tract, underscoring the importance of considering the sinuses as part of a unified airway system that plays a role in respiratory disease progression in CF.

The unified airway hypothesis suggests that the URT and LRT are subject to the same infectious and inflammatory events, and that managing upper airway health can benefit the LRT and vice versa (*70, 71*). We previously demonstrated a link between sinus disease in the form of a sinus exacerbation and increased risk of subsequent pulmonary exacerbation in this same cohort of adults with CF and CRS (*36*). In the present study, we found that the sinuses resemble reports of CF lungs from both a microbiome and *P. aeruginosa* population genetics perspective. Our cohort exhibited low microbial diversity in their sinuses, with most individuals harboring communities dominated by *Staphylococcus* spp. and/or Pseudomonadaceae. Furthermore, the population structure of *P. aeruginosa* in the sinuses resembled adult CF lung populations in that most study participants harbored a single clonal lineage of *P. aeruginosa* for the duration of our two-year study. Sinus isolates displayed a diversity of pathoadaptive phenotypes associated with long term persistence in the CF respiratory tract. When possible, we identified the causative mutations driving several of these phenotypes (e.g. increased biofilm formation caused by a mutation in hybrid diguanylate cyclase/phosphodiesterase, PA5295); the mutations occurred in genes previously associated with these phenotypes in studies of CF lung isolates. Together, these findings support a unified CF airway model in which the sinuses act both as a training ground for *P. aeruginosa* to begin adapting to the host environment and as a reservoir for seeding the lungs with these pre-adapted isolates. Continuing work is aimed at examining *P. aeruginosa* populations using paired sinus and sputum samples from CF adults and children to test the hypothesis that pathoadaptive mutations that first arise in the sinuses may subsequently become enriched in sputum populations. Our goal is to understand whether and how we may therapeutically relieve selective pressures in the sinuses (e.g. iron chelation therapy) in order to prevent or delay the evolution of pathoadaptive traits by *P. aeruginosa* in CF.

Mutators are commonly isolated from sputum of people with CF (*51*). Although the accumulation of beneficial traits along with many neutral or deleterious ones may be the consequence of a higher mutation rate, the mutator phenotype itself is unseen by selection. Rather, mutators evolve by the indirect effects of selection on linked traits, such as loss of acute virulence in CF, increased biofilm formation, metabolic adaptations, and antibiotic resistance (*51, 53, 58, 83–85*). The probability of this linkage increases in small populations under strong selection that are limited for beneficial genetic variation or under conditions when effects of drift relative to selection are amplified (*86*). Our study reveals two distinct pathways to a mutator phenotype that are consistent with strong selection and small population sizes: first, by more commonly recognized genetic defects in DNA repair like *mutS* and *mutL* (*87*), and second, by increased rates of transposition of IS resulting from weakened selection to control their activity. One lineage in this study isolated from Patient 24 combined both of these processes, where ISPa45 disrupted *mutS* and further accelerated evolutionary rate. The evolution of a mutator phenotype catalyzed by an IS element inactivating *mutS* is unusual and has been reported once previously for *P. aeruginosa* in CF (*67*). However, a recent study by Chu *et al*. identified an IS element inserted into *mutS* in a marine bacterium, *Vibrio splendicus*, and went on to predict additional putative inactivations of *mutS* by IS elements in hundreds of diverse Betaproteobacteria and Gammaproteobacteria from human and environmental sources (*88*). Therefore, IS elements likely play a common role in the evolution of mutator phenotypes, in addition to their more well-known roles imparting antimicrobial resistance and reduced virulence phenotypes. In Patient 9’s lineage, non-synonymous mutations in *dinB* and *dnaQ* coincided with the emergence of mutators, although we did not identify whether one or both genes was causative of the mutator phenotype. Overall, the prevalence of mutators in 3 of our 6 whole genome sequenced lineages suggests that diverse mutator genotypes persist as a result of linked selection on traits that confer an adaptive advantage to *P. aeruginosa* in the sinuses, as has previously be described in the lungs.

Although the recurrent evolution of mutator lineages in people with CF and CRS suggests that *P. aeruginosa* lives in small populations where drift is relatively strong, the spectra of observed mutations demonstrates that selection for adaptive traits in the infection remained effective. As evidence, most mutations during our two-year study were non-synonymous and many of the affected genes are common targets of mutations among *P. aeruginosa* isolated from CF sputum. Indeed, we were able to link many of these mutations to their expected phenotypes based on prior literature. However, there were two notable surprises that we encountered when linking genotypes to phenotypes. First, we observed instances where non-synonymous mutations became fixed in genes belonging to pathways that already appeared to be inactivated and we were unable to detect changes in expected phenotypes (Supplemental figures 6CDF and Supplemental table 8, Supplemental figure 7C-G and Supplemental table 9, and Supplemental tables 4 and 10). This was particularly pronounced for mutations in pyoverdine biosynthesis in lineages from Patients 9, 24, and 33 and mutations in biofilm formation genes in Patient 33’s lineage. In these cases, we hypothesize that the fixation of these non-synonymous mutations represents relaxed selection on these previously inactivated loci rather than adaptive evolution. Alternatively, these mutations may still provide a selective advantage in the host by fine tuning the phenotypes that only appeared inactivated in our *in vitro* assays if our experimental conditions were insufficient to detect these subtle changes. The second surprise when linking genotypes to phenotypes occurred in the phenotypically least pathoadapted lineage, Patient 32. The last isolate collected from this lineage acquired a frameshift mutation in the flagellar regulator FleS. However, rather than inactivating flagellar motility as would be expected if this lineage were following a typical route toward pathoadaptation to the respiratory tract, this mutation resulted in increased swimming motility. One explanation for this observation comes from the model of the sinuses acting as a training ground for environmental *P. aeruginosa* to pre-adapt to the host prior to reaching the LRT. In this case, Patient 32’s lineage could still be exploring mutational space as it evolves in the sinuses, but this weakly deleterious will eventually be removed from the population (*50*). An alternative explanation is that this lineage is actually well adapted for the environment within Patient 32’s sinuses and represents an alternative path, perhaps comparatively rare among evolved CF *P. aeruginosa* populations, taken by this lineage as it navigates the fitness landscape of the sinuses.

We also observed some genomic signatures consistent with *P. aeruginosa* recently transitioning to a host-restricted lifestyle during long term colonization of the sinuses, including proliferation of IS elements in Patient 24’s lineage and a trend toward the highest number of fragmented genes and pseudogenes in the most pathoadapted lineage (Patient 33’s lineage). Diverse host-restricted microbes, from human pathogens to insect endosymbionts, show evidence of genome degradation when compared to their closest free-living relatives (*89, 90*). While we do not know how long each of our study participants was infected by their respective *P. aeruginosa* lineage, it is clear that the evolutionary timescale of our study is very short compared to the timescales described in previous studies of early genome degradation (*91, 92*). Still, our observation of IS expansion and putative pseudogene formation among *P. aeruginosa* lineages provides new insight into evolutionary processes occurring during long term colonization of the CF respiratory tract by an opportunistic pathogen. All sinus isolate genomes had more IS elements and putative pseudogenes compared to the 3 reference genomes we compared them to: two isolates from acute infections (PAO1 and PA14) and one epidemic strain that retains the ability to transmit between people with CF (LESB58). That these *P. aeruginosa* lineages can accumulate inactivated genes and harbor proliferative IS and not be outcompeted by another *P. aeruginosa* strain or other bacteria in the sinuses further suggests that the population size is small and/or that these lineages are occupying distinct niches in the sinus environment. A small population size and a protected niche would result in decreased efficiency of purifying selection, allowing for the inactivation of genes through genetic drift and the proliferation of IS elements, as has been described in host-restricted bacteria (*74*). While CF sputum can contain up to 10^8^ CFU of *P. aeruginosa* per milliliter (*93*), in a study of explanted CF lungs, Jorth *et al*. found that *P. aeruginosa* populations become isolated within distinct regions of the lung and independently evolve (*94*). Furthermore, Bjarnsholt *et al*. showed that *P. aeruginosa* tends to reside in small, spatially organized aggregates consisting of between 10^1^ to 10^4^ cells, based on the approximate diameter of aggregates (*95*). Using quorum sensing as a model of microbial interactions, Darch *et al*. examined the working distance of small aggregates of *P. aeruginosa* and found that aggregates had to be within 120 to 180 µm apart to sense diffusible signals from each other (*96*). It is possible that selection in the form of competition with other microbes may share a similar working distance in the CF sinuses, allowing for small, isolated *P. aeruginosa* populations to evolve under the same rules governing small populations of host-restricted bacteria. Finally, the Liverpool epidemic strain LESB58 displayed the fewest signatures of host restriction, likely supporting its versatility in infecting multiple individuals as an epidemic strain. Future work is needed examining a larger collection of high quality genomes from *P. aeruginosa* isolated from different stages of CF respiratory infections to determine whether the signatures of early host restriction tend to track with time spent evolving in the host, and whether a lack of these genomic signatures is a feature of strains with epidemic potential.

Overall, we observed a combination of adaptive evolution and genome evolutionary patterns consistent with recent host-restriction as the forces shaping *P. aeruginosa* genome evolution in the sinuses of adults with CF. Future work will determine whether these two processes occur simultaneously or represent two successive steps in pathoadaptation whereby *P. aeruginosa* first adaptively evolves to achieve a high level of fitness before proceeding down a path that would ultimately result in genome degradation during chronic CF respiratory infections. Our findings underscore the contribution of CF CRS to overall CF respiratory health and raise the question of whether clinical management of URT disease in CF could include interventions aimed at therapeutically relieving the selective pressures in the sinuses that drive pathoadaptation.

## Supplemental text

### Pyoverdine production is lost as P. aeruginosa evolves in the sinuses

The most commonly mutated gene across all sequenced *P. aeruginosa* lineages in our study was PA2424/*pvdL*, a gene required for pyoverdine biosynthesis, which was mutated in five of six lineages (Supplemental table 6). *P. aeruginosa* has a variety of iron acquisition strategies, including two siderophores (pyoverdine and pyochelin), two systems for heme uptake (Has and Phu), and one system for ferrous iron uptake (Feo). During chronic CF lung infections, isolates from CF sputum are known to reduce their production of pyoverdine, a trait that is thought to be pathoadaptive (*97–99*). We sought to determine if *P. aeruginosa* sinus isolates also change their iron acquisition strategies by acquiring loss-of-function mutations in pyoverdine production, which would suggest that the selective pressures driving pathoadaptation of iron acquisition in the sinuses are similar to the lungs. We identified 33 unique mutations in genes involved in iron acquisition, occurring in five of the six study participants’ lineages (Supplemental table 8). Each of the five lineages with iron acquisition mutations had at least one mutation in a gene predicted to be involved in pyoverdine usage. We examined pyoverdine production by all six sequenced *P. aeruginosa* lineages and found that these mutations resulted in reduced pyoverdine production in isolates from two lineages. *P. aeruginosa* lineages in both Patients 41 and 52 evolved mutations that resulted in loss of pyoverdine production (Supplemental figure 6), either through mutation of transcriptional regulators *pvdS* or *ampR* (Patient 41) or through a single nucleotide deletion in a non-ribosomal peptide synthetase required for pyoverdine biosynthesis (*pvdL*; Patient 52). Interestingly, the pyoverdine mutations in lineages from Patients 9, 24, and 33 occurred in genetic backgrounds that did not produce appreciable levels of pyoverdine to begin with and thus, we were not able to observe any change in pyoverdine production under our experimental conditions. Overall, we observed at least one mutation in all *P. aeruginosa* iron acquisition pathways except for Has (heme uptake) and Feo (ferrous iron uptake; Supplemental table 5). This suggests that heme and ferrous iron may be abundant in the sinuses, providing the selective pressure for *P. aeruginosa* to inactivate its more energetically costly siderophore-based iron uptake systems, as has been suggested in the LRT (*100, 101*).

### Variability in motility and biofilm formation within and between sinus lineages

Expression of genes involved in motility and biofilm formation are deeply intertwined in *P. aeruginosa* through regulatory systems that mediate lifestyle changes between acute virulence and chronic infection phenotypes (*102–104*). If *P. aeruginosa* pathoadapts in the sinuses, then we expect to see mutations that inactivate acute virulence genes, promote biofilm formation, or alter other phenotypes associated with long term persistence in the CF respiratory tract. We used the virulence factor categorization from http://pseudomonas.com/virulenceFactorEvidence/list to identify mutations occurring in genes with homology to known virulence-associated genes in PAO1. We found that predicted virulence factors, or regulators thereof, were mutated in all 6 *P. aeruginosa* lineages. The fewest mutations in virulence-associated genes occurred in Patient 32’s lineage (the lineage that we predicted to be the least pathoadapted based on cytotoxicity with airway epithelial cells and inflammatory cytokine induction phenotypes), whereas Patient 9’s lineage (containing mutator isolates) had the most (Supplemental table 5). Of the 64 virulence-associated genes that were mutated, 54 of these had previously been reported to be mutated studies of CF sputum isolates. These mutated genes included the type 3 secretion system, pyoverdine production, and type IV pili. We confirmed that a number of these mutations were loss-of-function in the following sections.

While three of the six whole genome sequenced *P. aeruginosa* lineages had already lost swimming motility before the start of this study, a phenotype indicative of pathoadaptation in CF (Figure 1B), we observed loss of swimming motility evolve during our study in 1 lineage that we whole genome sequenced. The Patient 9 reference isolate (P9_14) displays swimming motility, whereas all mutator isolates display decreased swimming motility relative to the reference, and the latest two isolates completely lost the ability to swim (Figure 4C). The complete loss of swimming motility in the latest two isolates could be attributable to at least two non-synonymous mutations in these isolates that would be predicted to abrogate swimming motility: a single nucleotide deletion in *cheA* (PA1458) leading to a frameshift and/or a nonsynonymous mutation in the transcriptional regulator FleQ (PA1097) (Supplemental table 5) (*105, 106*). The intermediate level of swimming motility in isolates 55 and 56 could be attributable to non-synonymous mutations in genes that influence cellular levels of cyclic diguanylate monophosphate (c-di-GMP), including the hybrid diguanylate cyclase (DGC)/phosphodiesterase (PDE) *mucR* (PA1727) and the PDE *nbdA* (PA3311) in both isolates (as well as P9_129 and 136), and the Pil-Chp surface mechanosensor *pilY1* (PA4554) and hybrid DGC/PDE PA5295 in P9_56(*107–109*). Thus, in Patient 9’s *P. aeruginosa* lineage, mutations in mutator isolates first lead to a reduction in swimming motility through mutations that cause increased in c-di-GMP (a physiologic condition known to suppress swimming motility), followed by a complete loss of swimming motility in the latest isolates due to additional mutations in genes required for flagellum biosynthesis. Looking more broadly, five of the 14 study participants for whom we collected longitudinal *P. aeruginosa* isolates had isolates that both retained and lost swimming motility (Table 2), representing additional cases where alteration of flagellar motility may be adaptive in the sinuses. However, two of these changes in swimming motility were associated with the introduction of a genetically distinct strain of *P. aeruginosa*, as determined by RAPD pattern, suggesting strain replacement may be an additional source of phenotypic heterogeneity of acute virulence factors in the sinuses.

In addition to loss of swimming motility, we observed loss of two additional acute virulence traits: pyocyanin production and twitching motility. *P. aeruginosa* can secrete a redox active phenazine called pyocyanin, which is known to cause damage to epithelium and is required during infection in a murine acute pneumonia model(*110*). Whereas Patient 9’s earliest isolates constitutively produced pyocyanin during growth on LB or under pyocyanin-inducing conditions, the latest isolates in this lineage did not (Figure 4D). The loss of pyocyanin production could be explained by a single nucleotide deletion in *pqsE* (PA100)(*111*). We also observed loss of twitching motility, a phenotype that requires functional type IV pili, in a different *P. aeruginosa* lineage that retained swimming motility over the course of the study. Two isolates in Patient 52’s lineage (#78 and 79) had a full length allele of a gene required for twitching motility and Pil-Chp-mediated surface sensing (PA0413/*chpA*) and were the only isolates that displayed twitching motility in this lineage. Overall, twitching motility varied across longitudinal isolates from 5 study participants (Table 2).

Biofilm formation is thought to play a key role in the persistence of *P. aeruginosa* during chronic CF lung infection. Many genes that promote biofilm formation are also inversely co-regulated with genes required for acute virulence, through a number of two-component systems and transcriptional regulators(*103, 104*). Through this network architecture, *P. aeruginosa* can switch from acute virulence to chronic and biofilm-forming phenotypes through one or very few mutations in these regulators(*6*). For these reasons, we expected to see mutations that promote biofilm formation occur among the sinus isolates. All six *P. aeruginosa* lineages had mutations in genes that are known to play a role in biofilm formation, ranging from mutations in 3 genes in Patient 32’s lineage to 26 mutated genes in Patient 9’s lineage (Supplemental table 9). However, most of these mutations would be predicted to decrease biofilm formation in a commonly used *in vitro* biofilm assay (crystal violet assay). We sought to determine the extent to which sinus isolates could form biofilms after 24 hours in a crystal violet biofilm assay, using either a rich media (LB) or synthetic CF sputum media (SCFM). With a few exceptions, longitudinal isolates from within each study participant’s *P. aeruginosa* lineage tended to form the same amount of biofilm (Supplemental figure 7), in contrast to our hypothesis that biofilm formation would tend to be higher in the later isolates from each lineage compared to the earlier isolates. While most of Patient 9’s *P. aeruginosa* isolates formed biofilm at approximately the same level as the lab strain PAO1 in synthetic CF sputum media (SCFM) or in LB, the mutator isolate #56 formed significantly higher levels of biofilm in SCFM, similar to the amount of biofilm formed by PAO1 Δ*wspF,* a strain that constitutively produces high levels of c-di-GMP and biofilm (Figure 4E; Supplemental figure 7AB). Isolate #56 has a unique early stop codon mutation in PA5295, a gene that encodes a hybrid diguanylate cyclase/phosphodiesterase. Deletion of PA5295 in PAO1 is known to lead to the RSCV phenotype in PAO1 and leads to overproduction of the biofilm matrix polysaccharides Pel and Psl (*112*). Furthermore, all 3 study participants for whom we collected mucoid isolates in this study had both mucoid and non-mucoid *P. aeruginosa* isolates present in their sinuses (Table 2). Non-mucoid revertants of the mucoid phenotype are frequently co-isolated with mucoid isolates in sputum from people with CF and chronic *P. aeruginosa* infection (*113–115*). In Patient 33, the most phenotypically pathoadapted lineage in our study, we co-isolated a non-mucoid isolate with a unique non-synonymous mutation in *alg44* (PA3542; P33_24) relative to the other mucoid isolates collected on the same study visit in this person (Supplemental table 9).

### Evolution of antibiotic resistance in the sinuses during CF CRS

In addition to oral and systemic antibiotics, all six people for whom we sequenced *P. aeruginosa* genomes received topical antibiotic treatment in their sinuses during the course of the study, with topical gentamicin or mupirocin being the most common (Supplemental table 11). With the exception of variation in beta-lactamase and fluoroquinolone resistance genes, all 6 lineages contained the same acquired antibiotic resistance genes (Supplemental table 12). Antimicrobial resistance evolved in sinus isolates through mutations in genetic targets that lead to intrinsic resistance, with the most mutations in known antimicrobial resistance-associated genes occurring in Patient 9’s lineage (30 unique sites mutated) and the least occurring in Patient 32’s lineage (1 mutation) (Supplemental table 10). The most commonly mutated gene involved in antimicrobial resistance was PA4020/*mpl* (four of six lineages), the inactivation of which leads to overproduction of the AmpC β-lactamase (*116*). Efflux pump mutations were also common, with three lineages acquiring mutations in MexXY-OprM and individual lineages acquiring mutations in additional RND efflux pumps or their regulators.

To determine whether *P. aeruginosa* evolves toward increased antimicrobial resistance in the sinuses, we determined the minimal inhibitory concentration (MIC) of five classes of antibiotics in all isolates from the six whole genome sequenced lineages (Supplemental table 9). This subset was chosen such that an isolate from the first and last visit dates were included, as well as additional isolates if they had mutations in genes predicted to influence antimicrobial resistance. We observed a range of antimicrobial resistance phenotypes in the different lineages. Depending on the antibiotic tested, this included (1) single- or multi-step changes of MIC between sensitivity and resistance that tended to track with time (usually in Patients 9, 24, and 41), and (2) heterogeneity in resistance among isolates within a study participant’s lineage that did not tend to track with time (usually in Patients 33 and 52), and (3) very little evidence of evolution toward antimicrobial resistance (most consistently in Patient 32) (Supplemental table 4). These phenotypes likely reflect the population structure of *P. aeruginosa* in each study participants’ sinuses, with heterogeneity in resistance patterns sampled on a single date reflecting a more diverse *P. aeruginosa* population.

First, we observed a tendency for MICs to increase in isolates collected over time for some antibiotics, especially in lineages from Patients 9, 24, and 41 (Supplemental table 4). The minimal inhibitory concentration (MIC) of aminoglycosides gentamicin and tobramycin increased in a stepwise manner in two Patient lineages. For example, Patient 9’s lineage contains mutator isolates that displayed an incremental increase in MIC relative to the first isolate from this patient (P9_14; Supplemental table 10). The first increase in aminoglycoside MIC in isolates P9_55 and P9_56 relative to P9_14 could be attributable to nonsynonymous mutations in *ndvB* (PA1163), *fusA2* (PA2071), or *opmD* (PA4208), whereas the additional increase in aminoglycoside MIC last isolate (P9_136) could be attributable to a mutation in *fleQ* (PA1097; Supplemental table 12) (*117–120*). Incremental increases in aminoglycoside resistance over time were also observed in Patient 33’s lineage and were associated with mutations in *ptsP* (PA0337)*, nuoH* (PA2643), and *wbpZ* (PA5447) (*121, 122*). Aside from the aminoglycosides, stepwise MIC increases over time were also observed for ceftazidime and aztreonam in Patient 24, both of which may be attributable in part to mutations in *mexE* (PA2493) and *mpl* (PA4020; Supplemental table 10) (*94, 116, 123, 124*).

Secondly, we observed heterogeneity in resistance patterns for a given antibiotic within lineages (typically Patients 33 and 52), where no temporal trend of a change in MIC was observed. In some cases, this heterogeneity was due to the co-existence of sensitive and resistant subpopulations that we sampled from the start of the study, whereas in other cases this was due to sampled isolates switching between sensitive or resistant over time. For example, Patient 33’s lineage contained a mix of isolates that were sensitive or resistant to piperacillin and meropenem on both the first and last visit date sampled (Supplemental table 4). In contrast, Patient 9’s lineage appears to have become temporarily more sensitive to ciprofloxacin (isolates #55 and 56) before ultimately becoming resistant (isolate #136). The decreased MIC of the intermediate isolates relative to the first isolate in this lineage could be attributable to multiple mutations that occurred when isolates in this lineage became mutators, including an early stop codon in *mexB*, whereas ciprofloxacin resistance in the last isolate could be attributable to a mutation in *nalD* (Supplemental table 10) (*124, 125*).

Finally, there were some cases in which the MIC of an antibiotic did not change in a particular lineage during the study, either because all isolates from a lineage remained sensitive (largely Patient 32’s isolates, consistent with a recent acquisition of this clone into the sinuses) or because resistance had been fixed in the population prior to the start of our sample collection. Notably, Patient 32’s lineage (predicted to be the least pathoadapted based on cytotoxicity for airway epithelial cells and inflammatory cytokine induction phenotypes) contained only one mutation predicted to influence antimicrobial resistance (Supplemental table 10). The last isolate from this lineage displayed a modest increase in the MIC of one antibiotic (meropenem-vaborbactam), but overall remained sensitive to all antibiotics tested according to their respective CLSI breakpoints (Supplemental table 4). However, despite this *in vitro* antimicrobial susceptibility and the periodic use of topical anti-Pseudomonal antibiotics (in addition to oral and systemic antibiotics; Supplemental table 11), this *P. aeruginosa* lineage persisted in Patient 32’s sinuses.

## Methods

### Study design, cohort selection, and sinus sampling

We performed a prospective, longitudinal study of 33 CF adults with symptomatic CRS and prior functional endoscopic sinus surgery (FESS) (*36*). Participants were treated in a CF-focused otolaryngology clinic at the University of Pittsburgh. During quarterly clinic visits and unscheduled clinic visits, at least two sinus swabs were collected endoscopically for 16S rRNA gene amplicon sequencing and bacterial culturing, and sinus wash was collected for iron and cytokine analysis. Of the 33 participants enrolled in the study and for whom sinus samples had been collected, we collected microbiome samples on at least two different study visits from 18 people and *P. aeruginosa* isolates were collected on at least two study visits from 14 people.

### 16S rRNA gene amplicon sequencing

Sequences from the Illumina MiSeq were deconvolved, and then processed through the Center for Medicine and the Microbiome (CMM) in house quality control pipeline, which includes dust low complexity filtering, quality value (QV) trimming, and trimming of primers used for16S rRNA gene amplification. Each read was first trimmed until the QV was no longer below 30. If the trimmed read was shorter than 75bp, the read was discarded. The read was also discarded if less than 95% of the bases were above a QV of 30. The scripts, fastq_quality_trimmer and fastq_quality_filter, from Hannon’s Cold Spring Harbor Laboratory’s FASTAX-Toolkit were used by the pipeline to perform the trimming and filtering, respectively. Mated pair forward and reversed reads were merged using the following criterion: minimum required overlap= 25 bp, proportion overlap mismatch > 0.2 bp, maximum N’s allowed = 4, and a read length minimum of 125 bp. The 16S reads were then taxonomically classified through the CMM’s Mothur^16^-based 16S clustering and sequence annotation pipeline, that performed sequence classifications with the Ribosomal Database Project (RDP) reference sequences. To monitor for contamination, three environmental controls and two extraction kit controls were sequenced, along with two *E. coli* positive controls. The composition of control samples differed significantly from all sequenced study samples (PERMANOVA; p < 0.0001), as did the read counts for the environmental and kit controls compared to the study samples (T-test; p-value <0.0001).

### P. aeruginosa isolate collection

For every participant’s study visits, a sinus swab was streaked onto Pseudomonas Isolation agar and incubated at 37°C for 48 hours. Single colonies were selected based on colony morphology and frozen as a 30% glycerol stock at −80°C. Genomic DNA from all *P. aeruginosa* isolates was extracted using a QIAgen DNeasy kit (Qiagen, Hilden, Germany) and the 16S rRNA gene was amplified using primers 63f and 1387r (*126*). Amplicons were purified enzymatically with ExoSAP-It (Applied Biosystems) prior to Sanger sequencing (Eurofins Genomics, Louisville, KY) to confirm their identity as *P. aeruginosa*.

### P. aeruginosa isolate genotyping

Purified genomic DNA from *P. aeruginosa* isolates was genotyped by Random Amplification of Polymorphic DNA (RAPD) typing using primer 272 (*127*). RAPD amplicons were visualized on a 1% agarose gel that included a GeneRuler 1kb plus DNA ladder (Thermo Scientific, Waltham, MA) and a RAPD performed with PAO1 genomic DNA to aid in comparison of clinical isolates’ patterns. For each participant’s longitudinal *P. aeruginosa* collection, isolates were considered clonally-related if they had an identical RAPD pattern.

### P. aeruginosa isolate phenotyping

We screened for the following phenotypes by spotting 3µL of an LB overnight culture of each clinical isolate, as well as PAO1 or other controls, onto the appropriate agar plate. Verification of growth, colony size, and lysis and sheen phenotypes were determined after 48 hours of grown on LB agar (24 hours at 37°C and 24 hours at room temperature). Protease production was determined on Brain Heart Infusion (BHI) agar plates supplemented with 10% skim milk, after 24 hours of growth at 37°C. Rhamnolipid production was determined following the protocol by Köhler and colleagues (*128*). Rhamnolipid agar plates contained 42mM Na_2_HPO_4_, 22mM KH_2_PO_4_, 8.5mM NaCl, and 15g/L Noble agar supplemented with the following filter-sterilized ingredients after autoclaving: 0.1mM CaCl_2_, 2mM MgSO_4_, 0.2% glucose, 0.05% glutamate, 0.025% methylene blue, 0.2% Hexadecyltrimethylammonium bromide (CTAB), and 0.05% casamino acids. Rhamnolipid plates were incubated for 24 hours at 37°C, followed by 72 hours at room temperature in the dark. Mucoidy was determined on both Pseudomonas Isolation Agar and LB agar after 24 hours of incubation at 37°C, followed by 48 hours at room temperature. Controls used to determine mucoidy included PAO1 (non-mucoid), FRD1 (mucoid; contains the *mucA22* allele), and FRD875 (nonmucoid; FRD1 with a disrupted *algD*)(*129*). Hyper-pigment binding and rugosity were determined on Vogel-Bonner Minimal Media(*130*) (VBMM; 0.2 g/ L MgSO_4_•7H_2_O, 2.0 g /L citric acid, 3.5 g/L NaNH_4_HPO_4_•4H_2_O, 10 g/L K_2_HPO_4_, pH 7.0) with 10 mM citrate, 40μg/ml congo red, and 15μg/ml brilliant blue as previously described(*112*) The controls for hyper-pigment binding and rugosity phenotypes were PAO1 (not an RSCV) and PAO1 with a clean deletion of *wspF* (an RSCV(*131*)). Killing of *S. aureus* was determined using a modified version of a bacteriophage plaque assay(*132*) in which *S. aureus* SA113 was grown to OD_600_ 0.5 in TSA broth, then suspended at a 1:100 dilution in 0.7% LB agar that was autoclaved and cooled to 50°C in a water bath. *P. aeruginosa* isolates were spotted atop the *S. aureus* overlay plates, incubated at 37°C for 24 hours, and clearing of *S. aureus* from the soft agar overlay was indicative of *S. aureus* killing or growth inhibition by the *P. aeruginosa* isolate. A control for the *S. aureus* killing assay was PAO1, which inhibits the growth or kills *S. aureus* SA113 under these conditions. Swimming motility was determined by stabbing a pipette tip dipped in stationary phase *P. aeruginosa* into the middle of LB Lenox supplemented with 0.3% agar and incubating at 30°C for 24 hours (*94*). Twitching motility was determined similarly by stabbing the pipette tip to the bottom of an LB with 1% agar plate. After 48 hours at 37°C, the agar was carefully removed with a spatula, the plastic Petri dish was stained for 10 minutes with 1% crystal violet, and then washed with deionized water(*133*). Controls for motility assays were PAO1 (swims and twitches), PAO1 Δ*fliC* (does not swim), and PAO1 Δ*pilA* (does not twitch).

### Cytotoxicity and IL-8 secretion in a polarized CF airway epithelial cell model

CF airway epithelial cells (CFBE41o-) were seeded at 2.5×10^5^ cells per well onto 12-mm Transwell permeable membrane supports (Corning) and grown for 48 hours with minimal essential medium (MEM) supplemented with fetal bovine serum, penicillin, streptomycin, and plasmocin both apically and basolaterally. After 48 hours, cells were allowed to polarize by growing at the air-liquid interface for 5 additional days prior to experimentation. Cytotoxicity was assessed by measuring a change in transepithelial electrical resistance (TEER) over time, following inoculating filters apically with sstationary phase bacterial cultures at an MOI of 25-30. Thirty minutes prior to each TEER measurement, MEM clear media was added apically to polarized CFBE41o-and returned to incubation at 37oC with 5% CO2. After each TEER measurement time point, the apical media was removed and cells were allowed to incubate at 37oC with 5% CO2 until 30 minutes prior to the next TEER reading. Interleukin 8 (IL-8) secretion was measured by enzyme-linked immunosorbent assay (ELISA).

### Whole genome sequencing of longitudinal P. aeruginosa isolates

Whole genome sequencing was performed on all isolates from six study participants. The six participants were chosen based on their having more than six *P. aeruginosa* isolates in our collection, from more than three study visits. Genomic DNA from all *P. aeruginosa* isolates from patients 9, 24, 32, 33, and 41 was extracted using a QIAgen DNeasy kit (Qiagen, Hilden, Germany). All samples were sequenced to at least 80-fold mean read coverage on an Illumina NextSeq 500. Reads were trimmed using Trimmomatic version 0.36 in paired end mode, reads shorter than 70 nucleotides were removed, and trimmed reads were quality checked with FastQC version 0.11.5.

### Hybrid genome assembly of reference isolates, assembly of subsequent isolates, genome annotation, and phylogenetic analyses

We *de novo* assembled the genome of the first *P. aeruginosa* sinus isolate from each participant, using both Illumina short read and Oxford Nanopore long reads to create a hybrid assembly as a reference genome for each of the six study participants’ lineages (Supplemental table 1). The earliest isolate from each of the six participants was additionally sequenced on a MinION (Oxford Nanopore Technologies) and Guppy was used to call bases and trim barcodes. To create a separate *P. aeruginosa* reference isolate for each person, reads were *de novo* assembled using Unicycler version 0.4.4 in conservative mode. Contigs shorter than 1 kB were removed and the assemblies were quality checked with QUAST version 4.3. All Illumina-only sequenced subsequent isolates were assembled using SPAdes version 3.11.0 and contigs smaller than 1kB were removed. Any isolates with fewer than 95% mapped reads in the breseq analysis described below were examined for evidence of contamination by the following 3 methods: identifying contigs with comparatively low coverage, BLASTp searching each gene on each contig in its SPAdes assembly against RefSeq to determine if the most common top hit is from a species other than *P. aeruginosa,* and by examining the average GC content of the contig. Contaminating contigs were removed from 3 isolates prior to running prokka and roary: P32_36, P32, 108, and P41_119. All assemblies were annotated with prokka version 1.14.6. Core genome SNPs were identified by aligning genes in prokka-generated .gff files using roary (*134, 135*) version 3.12.0 and the following command with the optional flag --mafft. A phylogenetic tree was constructed from the core_gene_alignment.aln file with RAxML version 8.2.4 using the following command: raxmlHPC-PTHREADS -x 2421 -f a -m GTRGAMMA -n name_of_output_tree.txt -p 748 -s /path/to/roary/core_gene_alignment.aln -T 16 -w /path/to/output/directory -N 100.

### Estimating the ratio of substitution rates (dN/dS) for virulence-associated genes

We used codeML from the PAML software package (*136*) to calculate the ratio of non-synonymous mutations per non-synonymous site (dN) to synonymous mutations per synonymous site (dS). We included all 67 genomes in the analysis and used *P. aeruginosa* PA7 as a reference strain. Each of the 67 genomes was queried against the PA7 genome using DIAMOND BLASTp (*137, 138*). The results were then parsed with a custom python script, and pairwise proteins alignments between isolate and PA7 were generated using MUSCLE (*139*). The protein alignments and corresponding gene sequences were then used to generate codon alignments using the pal2nal.pl script, which is part of the PAML package. We used the M0 model (NSsites = 0), which calculates one dN/dS value (gene-wide average) for each codon alignment. To visualize differences in virulence capacity in Figure 2B, we took a subset of dN/dS values corresponding to virulence-associated genes, based on annotations of the PAO1 genome available at: http://pseudomonas.com/virulenceFactorEvidence/list. The PCA plot was generated in RStudio, using R version 3.5.1 (*140*), and the R package factoextra (https://cloud.r-project.org/web/packages/factoextra/index.html). To avoid missing values, which are incompatible with PCA analysis, homologs not present in all isolates and in PA7 were omitted from this analysis. The final plot represents dN/dS values for 236 genes out of the 369 genes annotated as virulence-associated in PAO1 from pseudomonas.com.

### Identification of mutations in longitudinal collections of P. aeruginosa isolates from six study participants

We used breseq version 0.31.0 in consensus mode to identify mutations in each person’s subsequent isolates relative to their *de novo,* hybrid*-*assembled reference genome (first isolate)’s prokka-generated .gff file. The optional flag --consensus-minimum-coverage-each-strand 4 was used to exclude mutations supported by very few reads. All mutations were manually inspected and curated to remove false positives due to misaligned reads.

### Comparison of core and accessory genomic contests between 6 whole genome sequenced lineages of P. aeruginosa

We used the command anvi-script-FASTA-to-contigs-db in anvi’o version 5.5.0 to create a contigs database containing the 6 hybrid-assembled reference isolates and the lab strain PAO1. The genomes were annotated with functional categories (Cluster of Orthologous Genes; COG) using anvi-run-ncbi-cogs with the optional flag --search-with blastp. Following generation of a genome storage profile with anvi-gen-genomes-storage, a pangenome analysis was performed with anvi-pan-genome using the following flags: --mcl-inflation 8 --use-ncbi-blast. Each strain’s unique accessory genes (gene clusters that were not present in any other clinical isolate or PAO1) were defined by a bin and bins were exported using anvi-summarize. The dot plot of counts of unique accessory genes per COG category was produced using DotPlot.R in the software package FeGenie (*141*).

### Permutation test for gene-level parallel evolution

We performed a permutation test in R to determine whether genes mutated 2 or more times could be due to random chance for each lineage, based on the number of unique non-synonymous mutations within coding regions of genes. For each lineage, we performed 10,000 trials in which we simulated the appropriate number of mutations within coding regions of genes on the PAO1 chromosome. The PAO1 genome in FASTA nucleotide format was read into R using the command readDNAStringSet from the R package Biostrings (*142*). We estimated the p value empirically, based on the proportion of permutations that yielded a value more extreme than the observed value we were testing (e.g. if testing the likelihood that one gene could be mutated two or more times due to random chance, then the p value reflects proportion of trials in which one or more genes were mutated two or more times). We omitted false positives in very long genes with multiple mutations due to their length based on a SNP density of 2 mutations per 1000 bp (*143*).

### Identification of fragmented genes and pseudogenes

Fragmented genes and pseudogenes were identified using the Pseudofinder software (https:// https://github.com/filip-husnik/pseudofinder/). To identify pseudogenes, Pseudofinder performs a DIAMOND BLASTp search (e-value cutoff of 1E-5) of query genes (ORFs in amino acid format) against a reference protein database. We used the first isolate from each of the 6 whole-genome sequenced lineages as the query, as well as PA14 and LESB58. The reference protein database in most cases consisted of PAO1 proteins; one of the isolates (P33_22), was queried against PA14 because that is its closest lab-strain relative. After BLASTp is complete, Pseudofinder analyzes the results and compares the query genes against their homologs from the reference database with respect to gene length and premature stop codons. Query genes that deviate more than 25% from the length of their lab strain homolog are flagged as pseudogene candidates. Additionally, fragmented pseudogenes are identified based on their homology to the same ORF in the reference database and the close proximity of gene fragments to each other. These gene fragments are likely derived from premature stop codons, frameshifts, or IS disruptions that result in two or more ORFs that belong to the same ancestral gene. However, gene duplications (i.e. two homologous genes encoded adjacent to each other) can also be identified using this approach. To address these cases, we identified and removed gene duplications from the pseudogene count by performing a BLASTp search (e-value cutoff of 1E-5) of the candidate fragments against each other. Candidate fragments with homology to each other were then manually inspected and removed from the list of pseudogene candidates.

### Pyoverdine production by P. aeruginosa

*P. aeruginosa* isolates were grown to stationary phase in LB broth at 37°C, then inoculated in M9 media (33.9g/L Na2HPO4, 15g/L KH2PO4, 5g/L KH4Cl, 2.5g/L NaCl0.6mM MgCl2, 1.75mM CaCl2)(*144*) supplemented with 20mM sodium succinate and 0.5% iron-limited casamino acids. The iron-limited casamino acids solution was generated by pre-treating the solution with 5g Chelex-100 per 100mL of casamino acids for 1 hour while stirring, then filter-sterilizing.

### Pyocyanin production by P. aeruginosa

Isolates were grown for 24 hours shaking at 37°C in 5mL of King’s A media. To extract pyocyanin, the 5mL cultures were centrifuged and supernatants were removed. To the supernatants, 3mL of chloroform was added, the solution was vortexed for 20 seconds, and then centrifuged for 10 minutes at 4,000 rpm. The chloroform layer was extracted, 1mL of 0.2M HCl added, then the solution was vortexed and centrifuged for 5 minutes at 4,000 rpm. The pink layer was removed and the absorbance at 520nm was measured. The pyocyanin concentration in micrograms per milliliter of culture supernatant was calculated by multiplying the absorbance at 520nm by 17.072 (the molar extinction coefficient of pyocyanin at 520nm)(*145*).

### Iron measurement in sinus wash

Total iron in sinus wash was measured using a QuantiChrom Iron Assay Kit (BioAssay Systems, Hayward, CA, USA).

### Cytokine panel

Cytokine levels in sinus wash were quantified using a Luminex 35-Plex Human Panel (Invitrogen, Frederick, MD, USA).

### Data availability and Reproducibility

All sequencing reads were deposited in NCBI under BioProject accession number TBD.

## Acknowledgements

We thank the people with CF who participated in our study and contributed to this research. This work was supported by a Cystic Fibrosis Foundation (CFF) Carol Basbaum Memorial Research Fellowship (ARMBRU19F0) and National Institutes of Health (NIH) T32HL129949 to CRA, NIH T32HL129949 and CFF MELVIN15F0 to JAM, CFF ZEMKE16A0 and NIH 5K23HL131930 to ACZ, NIH T32AI060525 and CFF ZAMORA20F0 to PZV, NIH T35DK065521 to ILF, NIH R01HL138630 to AM, NIH R21HL143091 to BAM, University of Pittsburgh CTSI Pilot Program and NIH NCATS UL1 TR0000005 to SEL and JMB, NIH R61HL137077 and GILEAD Investigator Sponsored Research Award to SEL, VSC, and JMB, NIH U01AI124302 to VSC, and NIH R01HL123771, P30DK072506, CFF BOMBER14G0, and CFF RDP BOMBER19R0 to JMB. The graphical abstract was made using BioRender.com.

---

**Supplemental figure 1.**
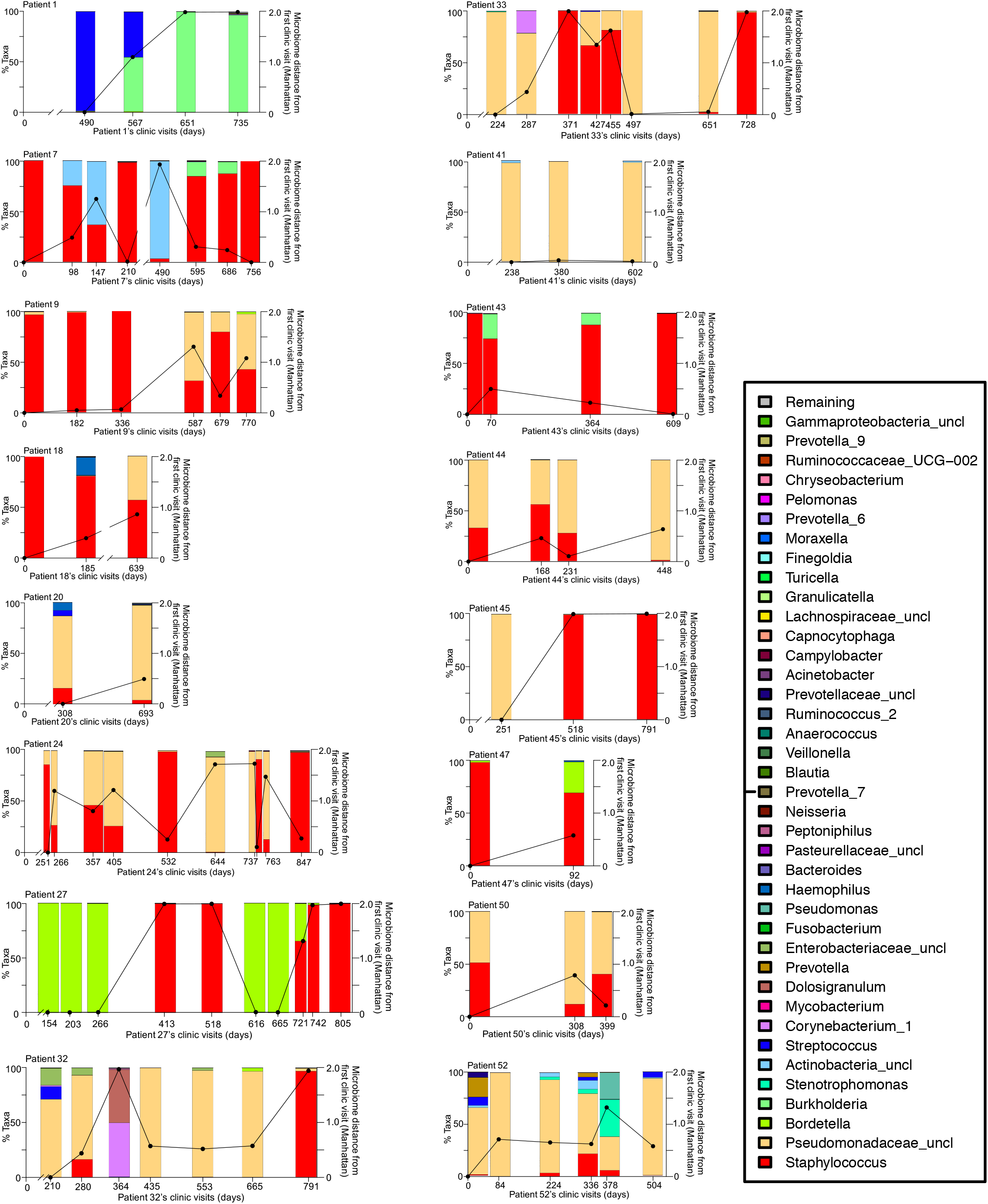
The sinus microbiome of adults with CF CRS contains traditional CF pathogens. Taxa bar plots depicting the percent of each OTU present at clinic visit dates for the 18 study participants for whom we sequenced microbiome samples at two or more clinic visits. Colors represent the taxa in the key at the right. *Staphylococcus* spp. is red, Pseudomonads are represented by Pseudomonadaceae in peach and *Pseudomonas* spp. in dark teal. Numbers beneath each plot are the days since study enrollment for each person. Plotted on the right Y axis is the microbiota dissimilarity of each timepoint compared to the first sample (Manhattan distance).

**Supplemental figure 2.**
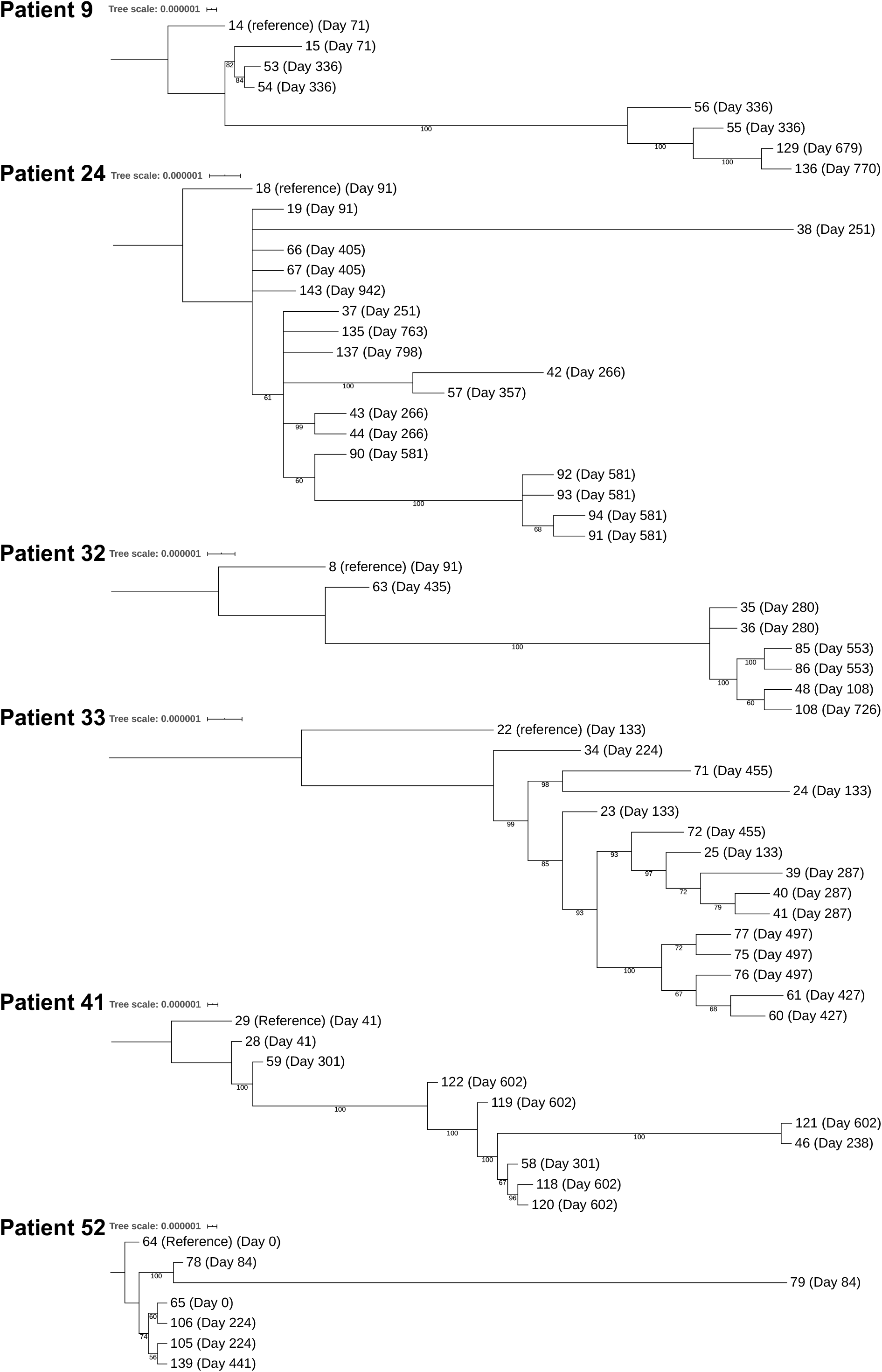
Six individuals harbor genetically distinct, clonal lineages of *P. aeruginosa* that evolved during the study. Date of isolation tends to track with phylogenetic relationships of isolates, suggesting isolates are representative of a population of *P. aeruginosa* that is evolving over time. Phylogenetic trees based on SNPs in core genomes from roary. Trees were built with RAxML using the GTRGAMMA model with 100 bootstraps. Each tree was rooted to the isolate used as the reference for breseq. Dates of isolation are listed next to each isolate. Branches with bootstrap values <50 are collapsed, and bootstrap values > 50 are listed for all branches.

**Supplemental figure 3.**
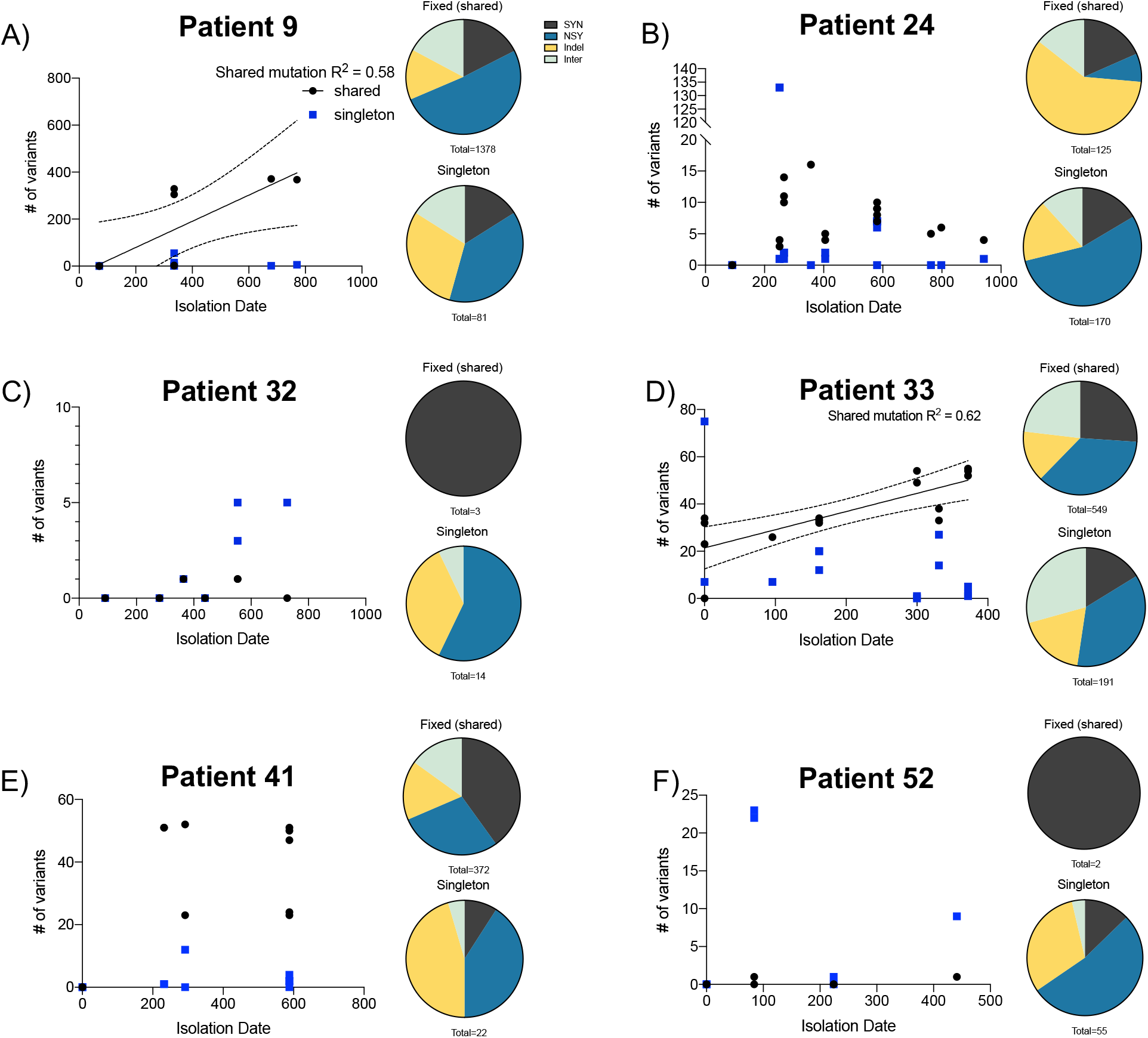
Variability in mutation fixation among *P. aeruginosa* lineages from the sinuses of six people with CF. Number of shared (black) and singleton (blue) mutations are plotted per isolate collected over time. Note that the x and y axes varies for the different participants’ lineages because of variability in the amount of time in the study and the number of mutations occurring in each lineage. Each *P. aeruginosa* isolate is represented by one black and one blue dot at the time point when it was collected. Mutations were “shared” if they occurred in at least one isolate collected on a different study date. Singleton isolates were not present in an isolate outside of that study date. Multiple pairs of black and blue dots indicate that more than one isolate was collected at that time point. Linear regression was performed for shared counts from each person, and regression lines and R squared values are displayed when p < 0.05 for the regression slope. Dashed line is 95% confidence interval. The pie charts next to each lineages’ plot depict the percentage of either fixed or singleton mutations that were either synonymous (SYN; grey), non-synonymous (NSY; blue), Insertion or deletion within a coding region (Indel; yellow), or a mutation occurring in an intergenic region (Inter; green). Below each pie chart is the sum of all fixed or singleton mutations that occurred in all isolates from that lineage over the course of the study.

**Supplemental figure 4.**
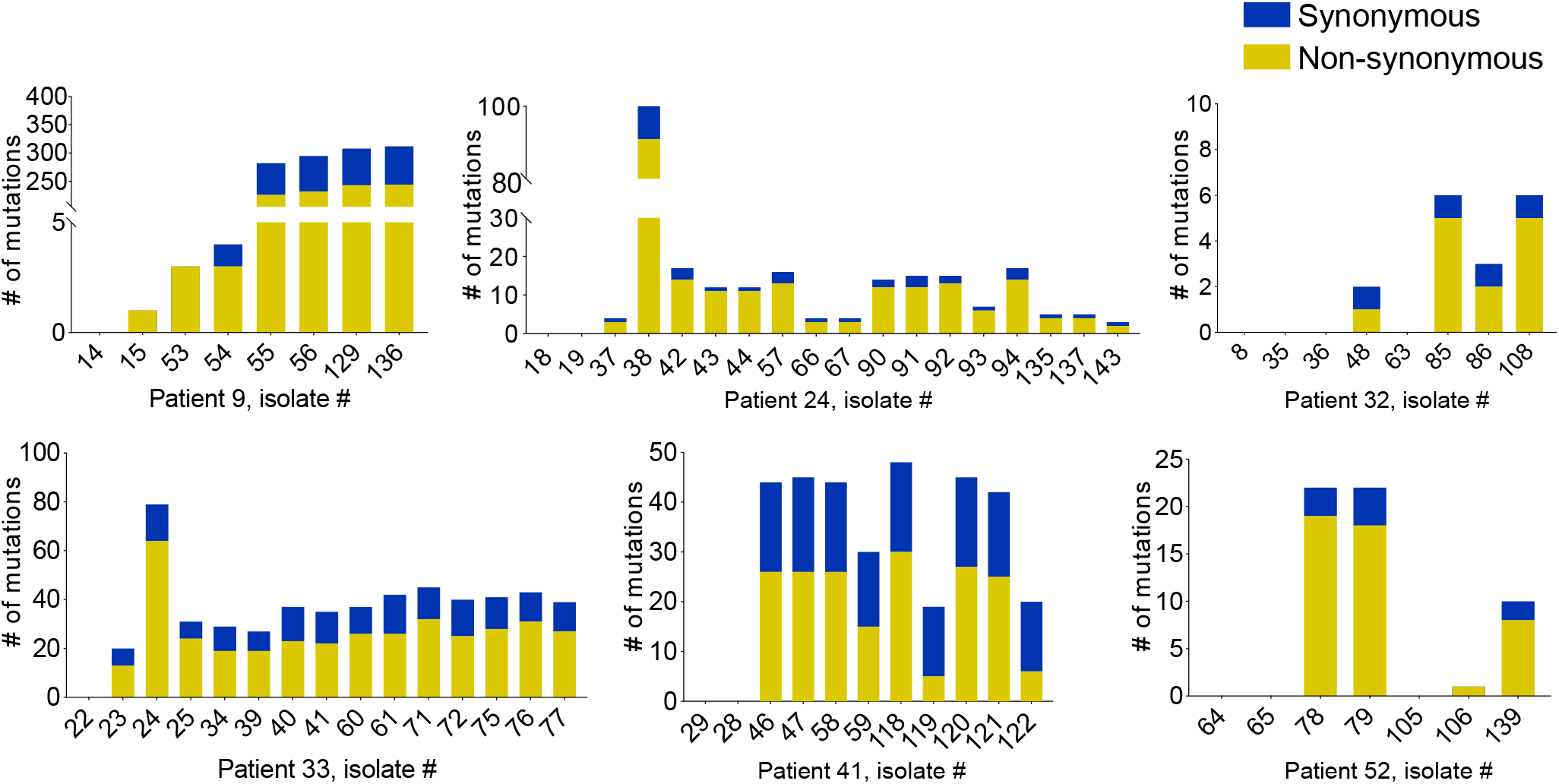
Isolates tend to have more non-synonymous mutations than synonymous mutations across all six sequenced lineages. Only mutations within coding regions of genes are displayed. Blue = synonymous mutations. Yellow = nonsynonymous mutations. Nonsynonymous mutations include deletions of entire genes. Putative mutations or genomic rearrangements listed as “Unassigned missing coverage evidence” or “Unassigned new junction evidence” in breseq’s output were not included.

**Supplemental figure 5.**
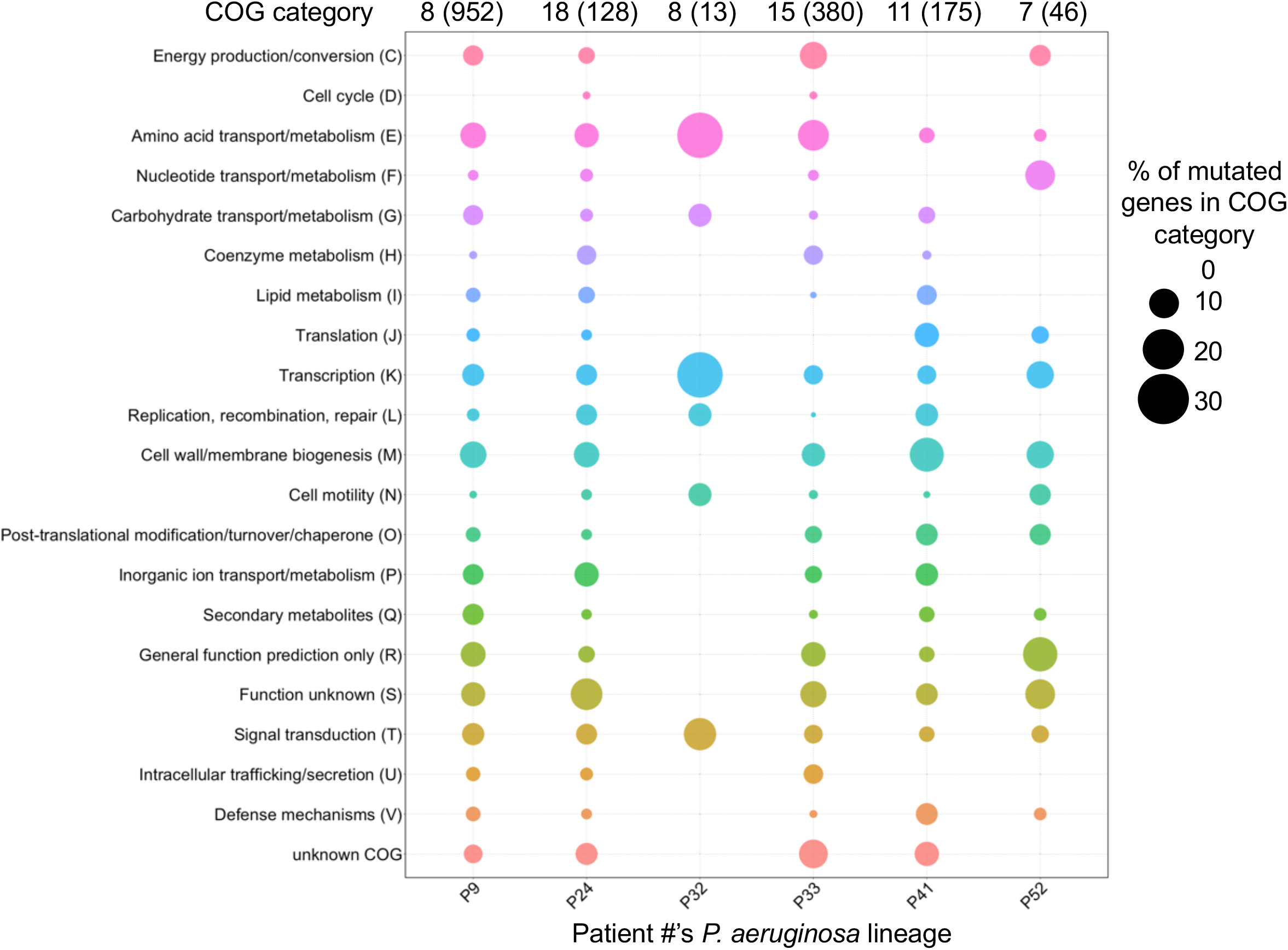
*P. aeruginosa* lineages undergo pathway-level mutational parallelism. All lineages acquired mutations in genes involved in amino acid transport and metabolism, motility, transcription, and signal transduction. Each column of dots represents the distribution of all non-synonymous mutations into the functional (COG) category of the gene they occurred in. Numbers starting with a “P” below each column denote the patient lineage. Numbers above the bars indicate the number of isolates in each lineage and, in parentheses, the total number of non-synonymous mutations that occurred in all isolates of that lineage, relative to the reference isolate. Mutations were called using breseq with a *de novo* assembled first isolate (hybrid nanopore and Illumina assembly) used as the reference.

**Supplemental figure 6.**
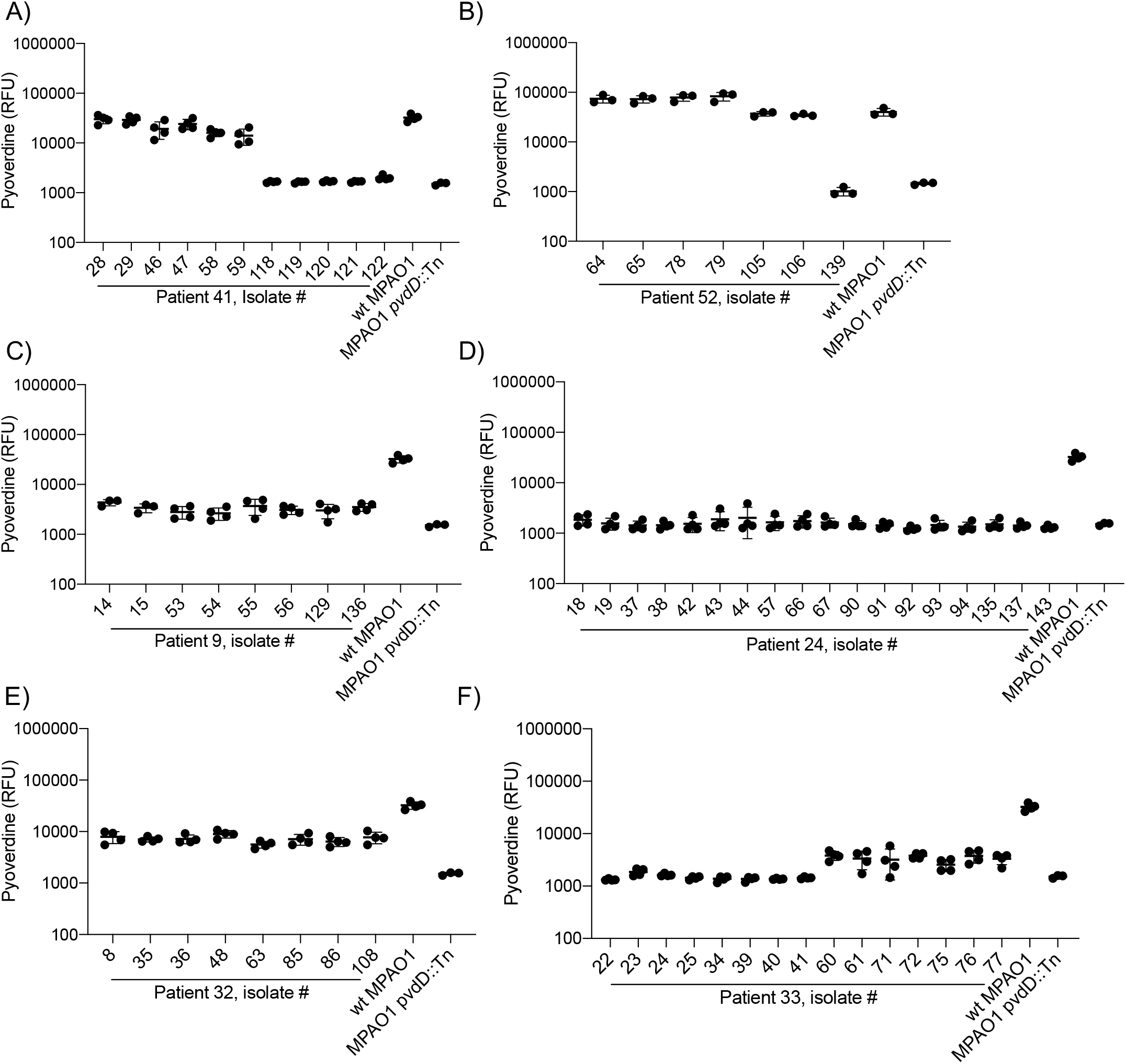
Isolates from Patients 41 and 52 evolve to produce less pyoverdine over time. Isolates were grown overnight in M9 minimal media with iron-limited casamino acids. To promote growth of auxotrophic isolates, adenosine was added to the media for all isolates and controls in B. **A)** Relative to the reference isolate #28, isolates #118 through 122 from Patient 41 all have mutations in genes required for pyoverdine regulation (a non-synonymous substitution in *pvdS* in isolates 118-121 or *ampR* in #122) and produce less pyoverdine during growth. **B)** Relative to the reference isolate #64, isolate #139 has a single base pair deletion in *pvdL* and does not produce pyoverdine. **C)** No change in pyoverdine in Patient 9’s lineage. **D)** No change in pyoverdine in Patient 24’s lineage. **E)** No change in pyoverdine in Patient 32’s lineage. **F)** No change in pyoverdine in Patient 33’s lineage.

**Supplemental figure 7.**
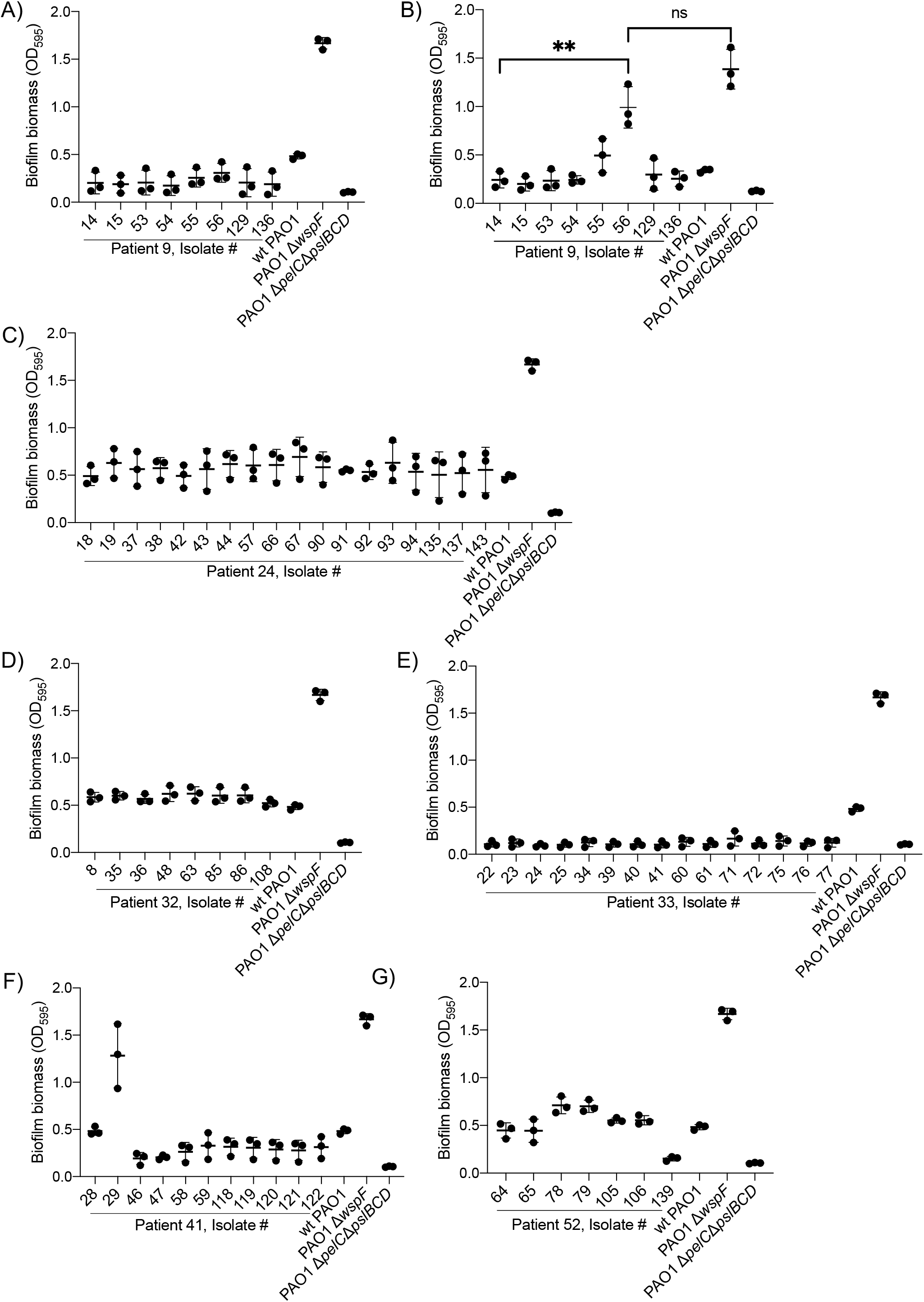
Biofilm formation of six whole genome-sequenced *P. aeruginosa* lineages. Biofilm formation in Patient 9 in LB **(A)** and SCFM **(B)**, **(C)** Patient 24, **(D)** Patient 32, **(E)** Patient 33, **(F)** Patient 41, and **(G)** Patient 52 was measured by crystal violet assay after 24 hours of growth in LB media at 37°C. Data are from 3 biological replicate experiments. Asterisk = p < 0.05 relative to wild type PAO1; ns = not significant (p > 0.05 by one-way ANOVA with Tukey’s multiple comparisons test).

**Supplemental Table 1.**
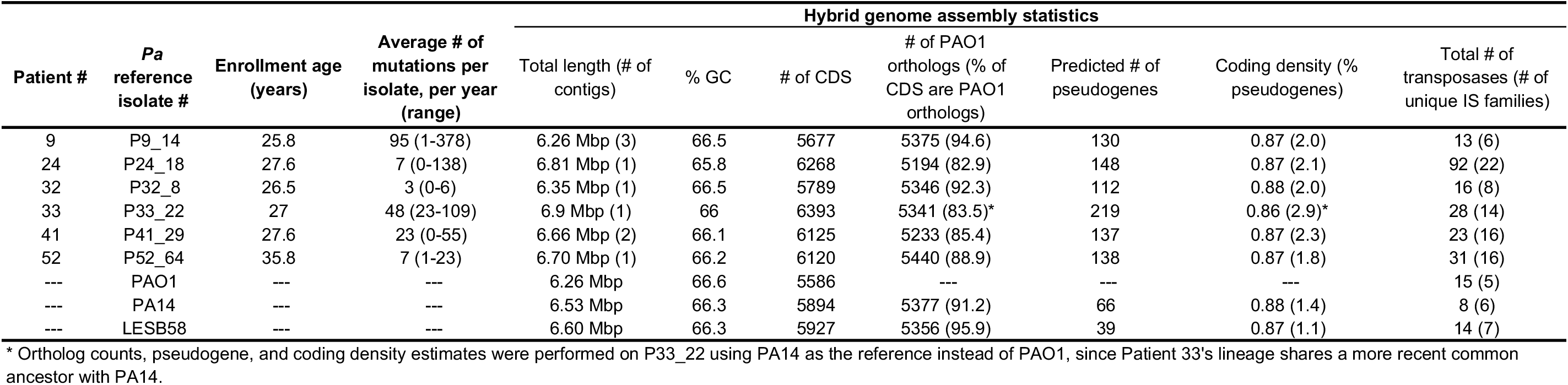
Characteristics of the hybrid-assembled genomes sequenced for the first isolate from the 6 study participants chosen for *P. aeruginosa* whole genome sequencing. The average number of mutations per isolate per year was calculated based all isolates from a participant’s *P. aeruginosa* lineage. The number of mutations includes SNPs, insertions and deletions, but not genomic rearrangements (“Unassigned new junction evidence” in breseq) or putative deletions that were called as “Unassigned missing coverage evidence” in breseq. The range of mutations represents the smallest and largest number of mutations that a single isolate had, relative to the reference isolate from that same patient. The lab strains PAO1, PA14, and LESB58 are included as a references for certain statistics. PAO1 orthologs in each reference genome were identified by nucleotide BLAST with default parameters and with PAO1 as the subject sequence. The number of pseudogenes was predicted for clinical isolates using gene length and fragmentation relative to orthologs in PAO1, using Pseudofinder (https://github.com/filip-husnik/pseudofinder). The coding density is the length in nucleotides of predicted coding sequences (CDS; after subtracting the length of all putative pseudogenes), normalized to the total length of the genome in nucleotides. The percent pseudogenes is the percentage of the total predicted CDS length that is pseudogenized. Number of unique IS element families is the number of transposases on that genome, excluding copies of the same transposase. The number of transposases in lab strains was determined based on the functional name from InterProScan on pseudomonas.com and the number of unique transposases was based on InterPro accession number/annotation. For ORFs that were assigned multiple transposase InterPro accession numbers, the annotation with the lowest Expect value (E) from a BLAST alignment was chosen.

**Supplemental table 2.** Predicted prophages present in the hybrid assembled first isolate from each of the 6 *P. aeruginosa* lineages for which WGS was performed. Prophages and their level of completeness were predicted with Phaster2.

| Isolate name | Predicted phage ID | Length (Kb) | # of proteins | Completeness |
| --- | --- | --- | --- | --- |
| **Patient 9_14** |  |  |  |  |
| Region 1 | phiCTX | 18.4 | 24 | questionable |
| Region 2 | Pf1 | 6.6 | 8 | questionable |
| **Patient 24_18** |  |  |  |  |
| Region 1 | JBD93 | 40 | 57 | intact |
| Region 2 | PMG1 | 31.9 | 35 | questionable |
| Region 3 | PMG1 | 43.6 | 53 | questionable |
| Region 4 | Pf1 | 18.8 | 10 | questionable |
| Region 5 | phiCTX | 18.4 | 25 | questionable |
| Region 6 | phiCTX | 40.1 | 48 | intact |
| **Patient 32_8** |  |  |  |  |
| Region 1 | phiCTX | 18.4 | 23 | questionable |
| Region 2 | JDB93 | 36.3 | 51 | questionable |
| Region 3 | phi92 | 26.2 | 8 | incomplete |
| **Patient 33_22** |  |  |  |  |
| Region 1 | JBD69 | 35.6 | 51 | incomplete |
| Region 2 | SPN1S | 10.9 | 8 | incomplete |
| Region 3 | phi2 | 44.7 | 63 | intact |
| Region 4 | Pf1 | 19.3 | 12 | questionable |
| Region 5 | F10 | 53.6 | 65 | intact |
| Region 6 | Pf1 | 27.1 | 29 | questionable |
| Region 7 | YMC11/R656 | 30.6 | 36 | intact |
| Region 8 | JBD69 | 34.2 | 47 | incomplete |
| Region 9 | phiCTX | 48.8 | 50 | intact |
| **Patient 41_29** |  |  |  |  |
| Region 1 | YMC11/02/R656 | 20.2 | 24 | incomplete |
| Region 2 | Pf1 | 13.8 | 13 | intact |
| Region 3 | Pf1 | 18.9 | 10 | incomplete |
| **Patient 52_64** |  |  |  |  |
| Region 1 | YMC11/02/R656 | 29.5 | 32 | intact |
| Region 2 | phiCTX | 51.1 | 47 | intact |
| Region 3 | YMC11/02/R656 | 62.9 | 57 | intact |
| Region 4 | Pf1 | 15.1 | 16 | questionable |

**Supplemental table 3.**
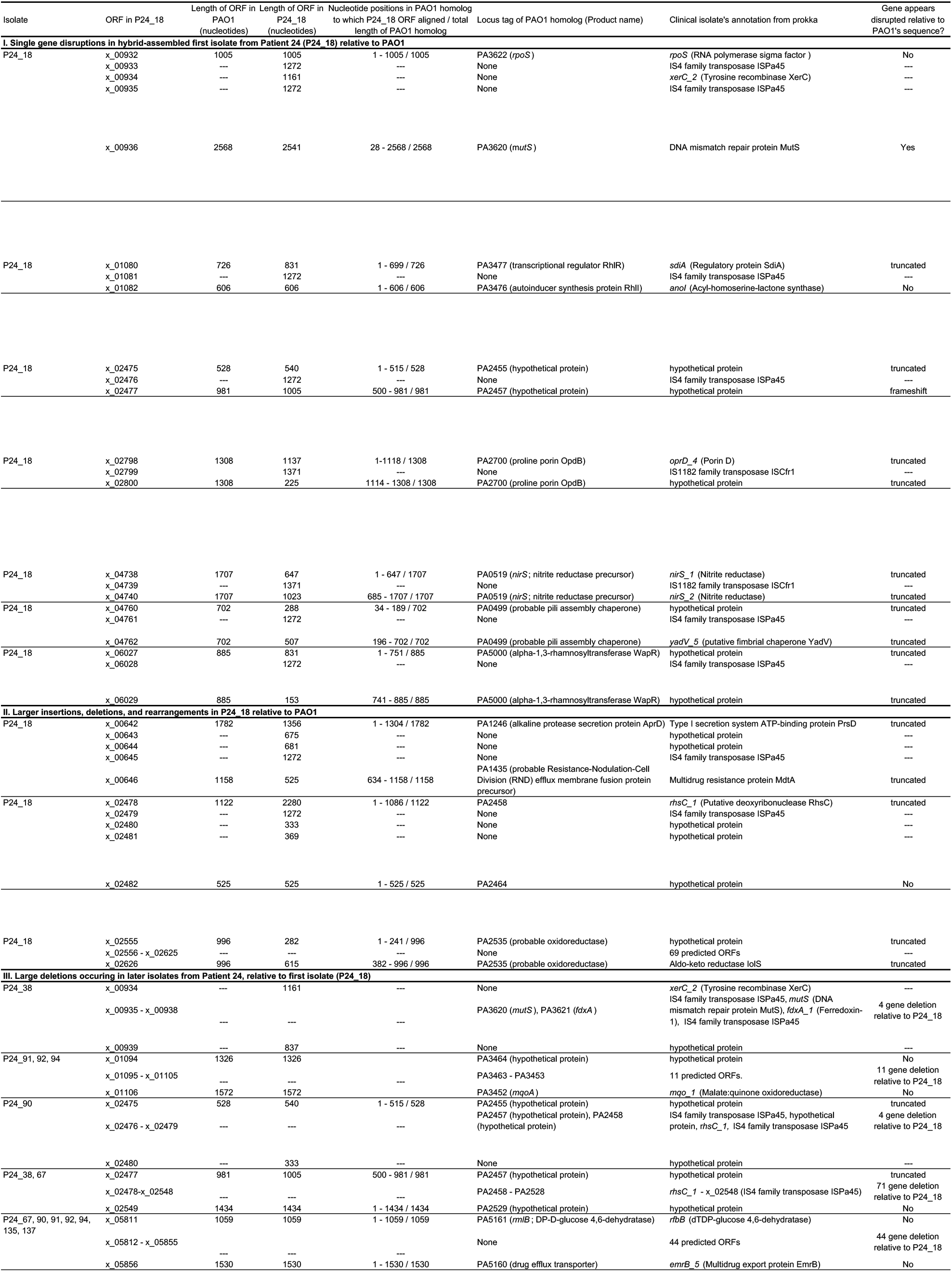
Summary of transposase insertions that appear to disrupt coding sequences of genes in Patient 24’s *P. aeruginosa* lineage. Sections I and II describe transposase insertions or deletions that were present in the hybrid-assembled first isolate from this lineage (P24_18), when compared to predicted homologs in PAO1. Predicted ORFs in P24_18 are only annotated with a PAO1 homolog if the aligned region of the query sequence (P24_18 ORF) shared >95% nucleotide identity with the subject sequence (PAO1 gene) to which it aligned by BLASTn. Section III describes apparent deletions associated with transposases in later Patient 24 isolates, relative to the hybrid-assembled reference genome P24_18. These deletions were identified by breseq and appeared in either the “Predicted mutations” or “Missing coverage” section. Information from “Unassigned new junction evidence” was not included in this table. The clinical isolate’s annotation was provided by prokka and is included because not every ORF has a PAO1 homolog. While the P24_18 genome harbors 92 predicted transposases from 22 transposase families, IS4 family transposase ISPa45 is the most common transposase (45 copies) and seems to be the one that most frequently disrupts predicted genes.

**Supplemental table 4.**
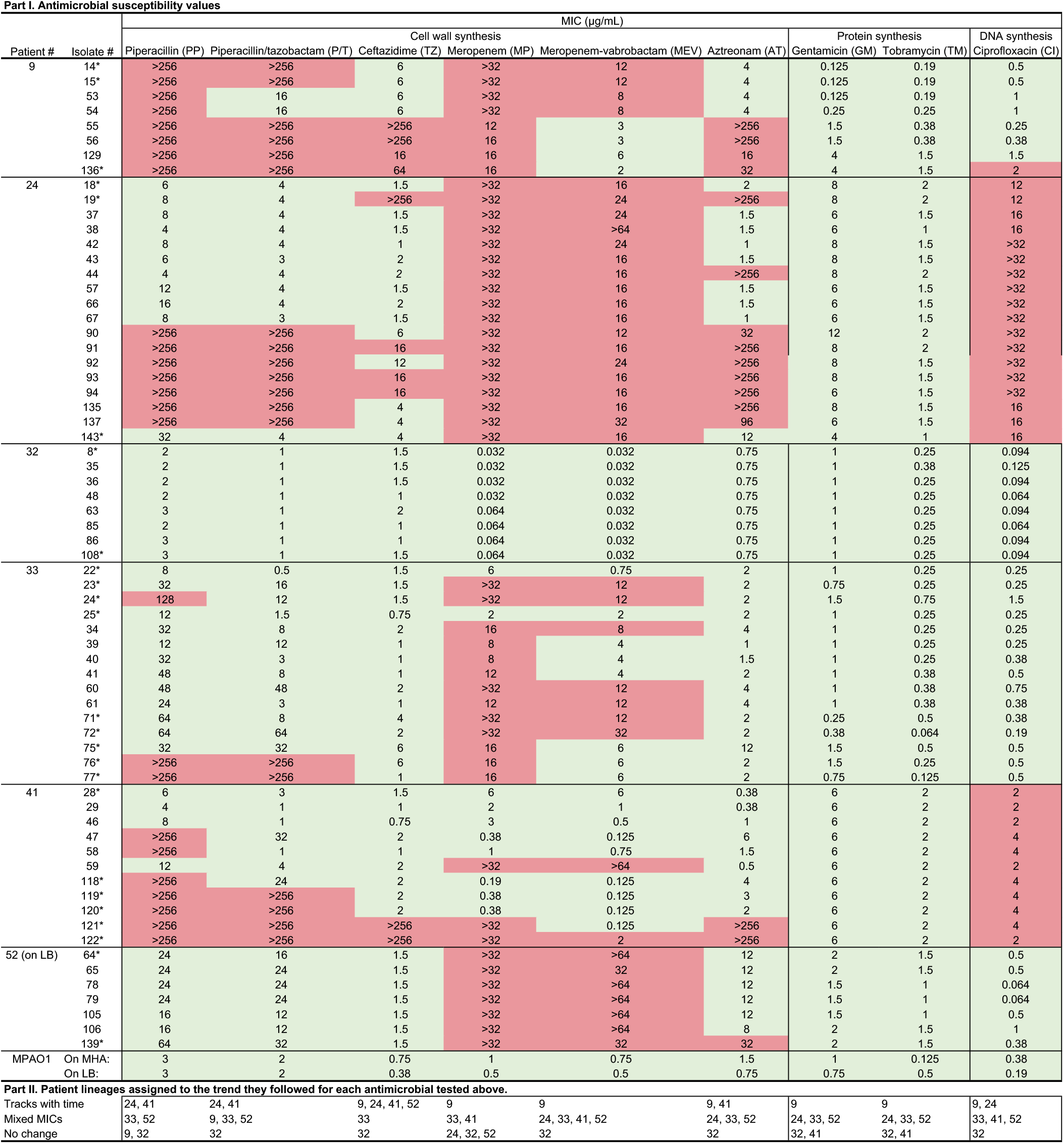
Part I of this table displays minimal inhibitory concentrations of select isolates from 6 patients’ whole genome sequenced *P. aeruginosa* lineages. Isolates that were collected on the first or last visit date are marked with an asterisk (*) for each patient. Additional isolates from each lineage were included based on their mutations predicted to influence antimicrobial resistance in Supplememtal table 8. For Patient 24, isolates 90 and 91 were included instead of isolate #143 because #143 did not have any mutations predicted to influence antimicrobial resistance. E-tests were performed as biological triplicates on Muller-Hinton agar (MHA) and a single representative value is listed in the table. For Patient 52, most isolates did not grow on MHA, so these E-tests were performed on LB agar. CLSI breakpoints of resistance for each antibiotic in micrograms per mililiter are as follows: piperacillin (≥ 128), ceftazidime (≥ 16), meropenem (≥ 8), aztreonam (≥ 16), gentamicin (≥ 16), tobramycin (≥ 16), ciprofloxacin (≥ 2). Resistance based on CLSI breakpoints is colored red, whereas intermediate or susceptible conditions are marked as green. Part II of this table shows which patient lineages fell into 1 of 3 trends for evolution of antimicrobial resistance, for each drug tested: (1) “Tracks with time” indicates single-or multi-step changes of MIC between sensitivity to resistance that tended to track with time (usually Patients 9, 24, and/or 41), (2) “Mixed MICs” indicates heterogeneity in resistance among isolates within a patient’s lineage that did not tend to track with time (usually Patients 33 and 52), and (3) “No change” indicates that there was very little to no evidence of evolution of antimicrobial susceptibility (most consistently Patient 32). A patient’s *P. aeruginosa* lineage can only be assigned to one category per drug tested.

**Supplemental table 5.**
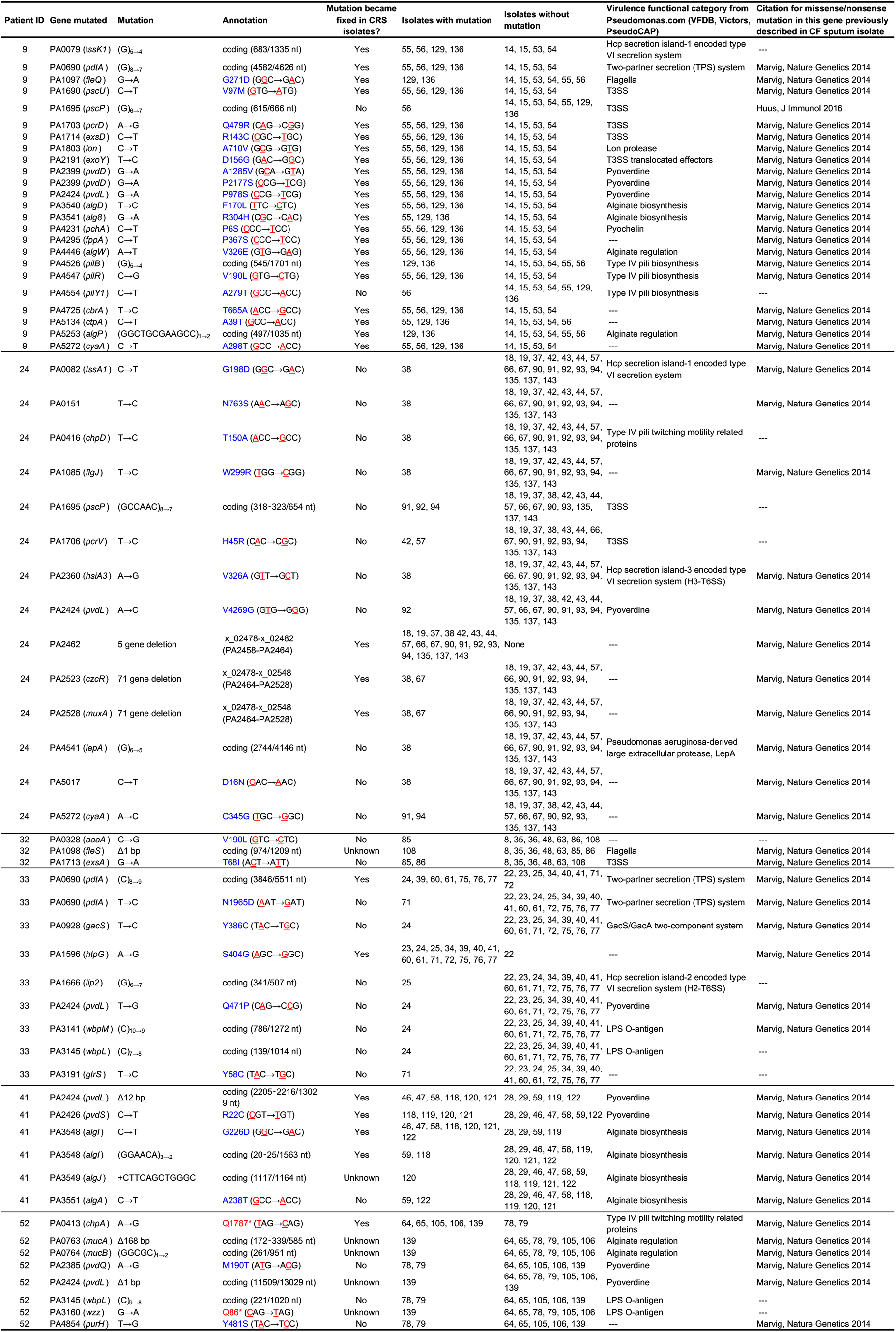
Virulence genes with non-synonymous mutations within the coding region. Genes were identified as virulence genes based on the classification from http://pseudomonas.com/virulenceFactorEvidence/list, which compiles data from VFD, Victors, and PseudoCAP. Dashed line indicates that no functional category was described or that no citation was found for a non-synonymous mutation in that gene among published reports of CF sputum isolates. Mutations were described as “fixed” in the population if they were present in isolates collected in at least two time points.

**Supplemental table 6.**
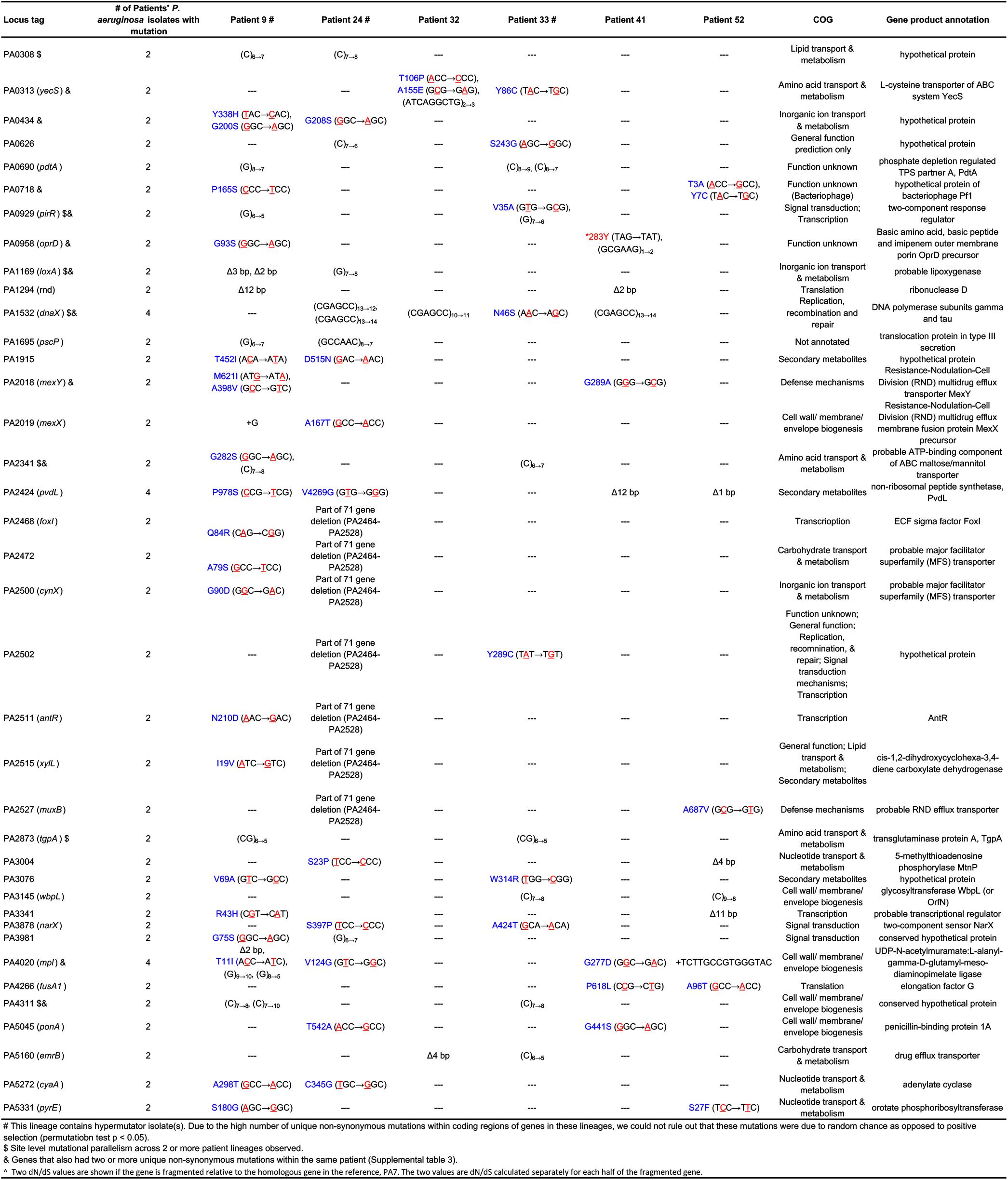
Genes with non-synonymous mutations occurring in two or more patients’ *P. aeruginosa* lineages. For *P. aeruginosa* lineages belonging to patients 32, 41, and 52, these mutations were found to be unlikely to be due to random chance (p <0.05; permutation test) and therefore likely represent examples of parallel adaptation occurring between patient lineages in the sinuses. Additionally, in all lineages, any gene mutated 4 times was unlikely due to random chance and is an example of parallel adaptation (p <0.05; permutation test).

**Supplemental table 7.**
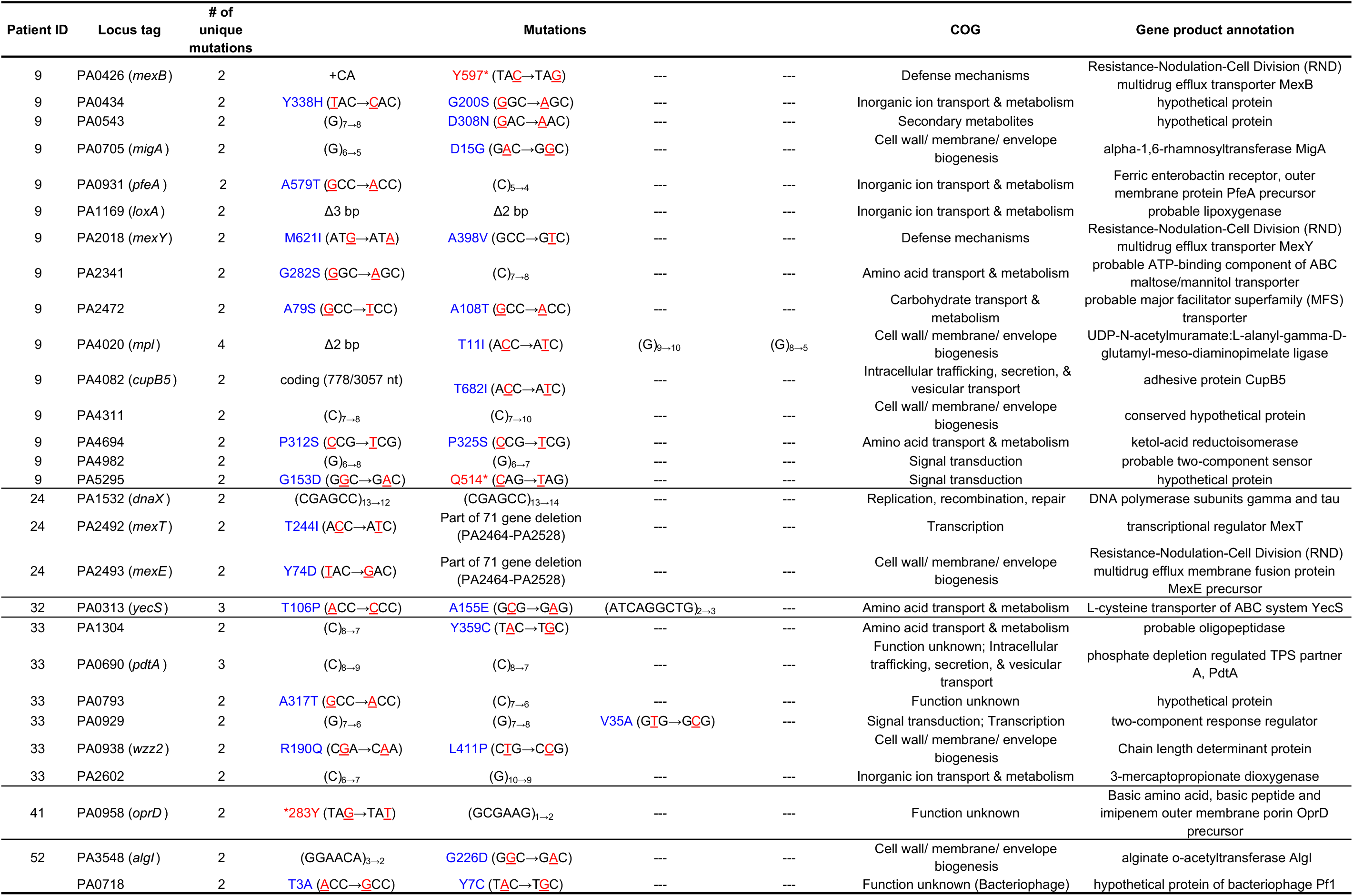
Genes with two or more unique, non-synonymous mutations occurring within a single patient’s *P. aeruginosa* lineage. Clusters of Orthologous Groups (COG) categories were assigned first based on annotation of the PAO1 genome by pseudomonas.com. Genes that did not have a COG annotation in the PAO1 genome were annotated with eggnog. Patients 9, 24, and 33’s lineages include multiple hypermutator isolates.

**Supplemental table 8.**
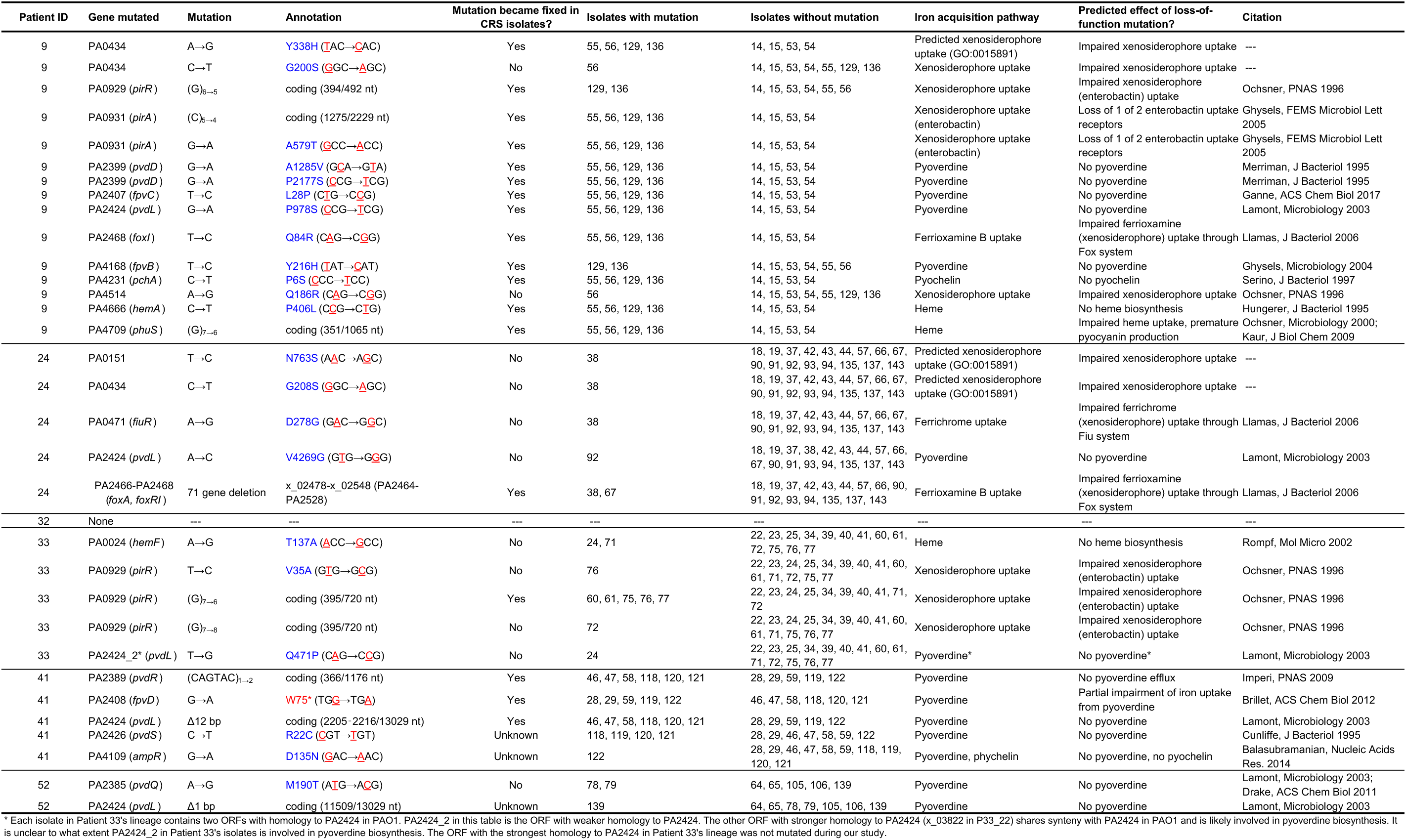
Mutations in iron acquisition genes. Mutation and annotation columns are imported from breseq. The predicted effect of a loss-of-function mutation in each gene is based on the citation(s) listed. Mutations were described as “fixed” in the population if they were present in isolates collected in at least two time points, including the last time point.

**Supplemental table 9.**
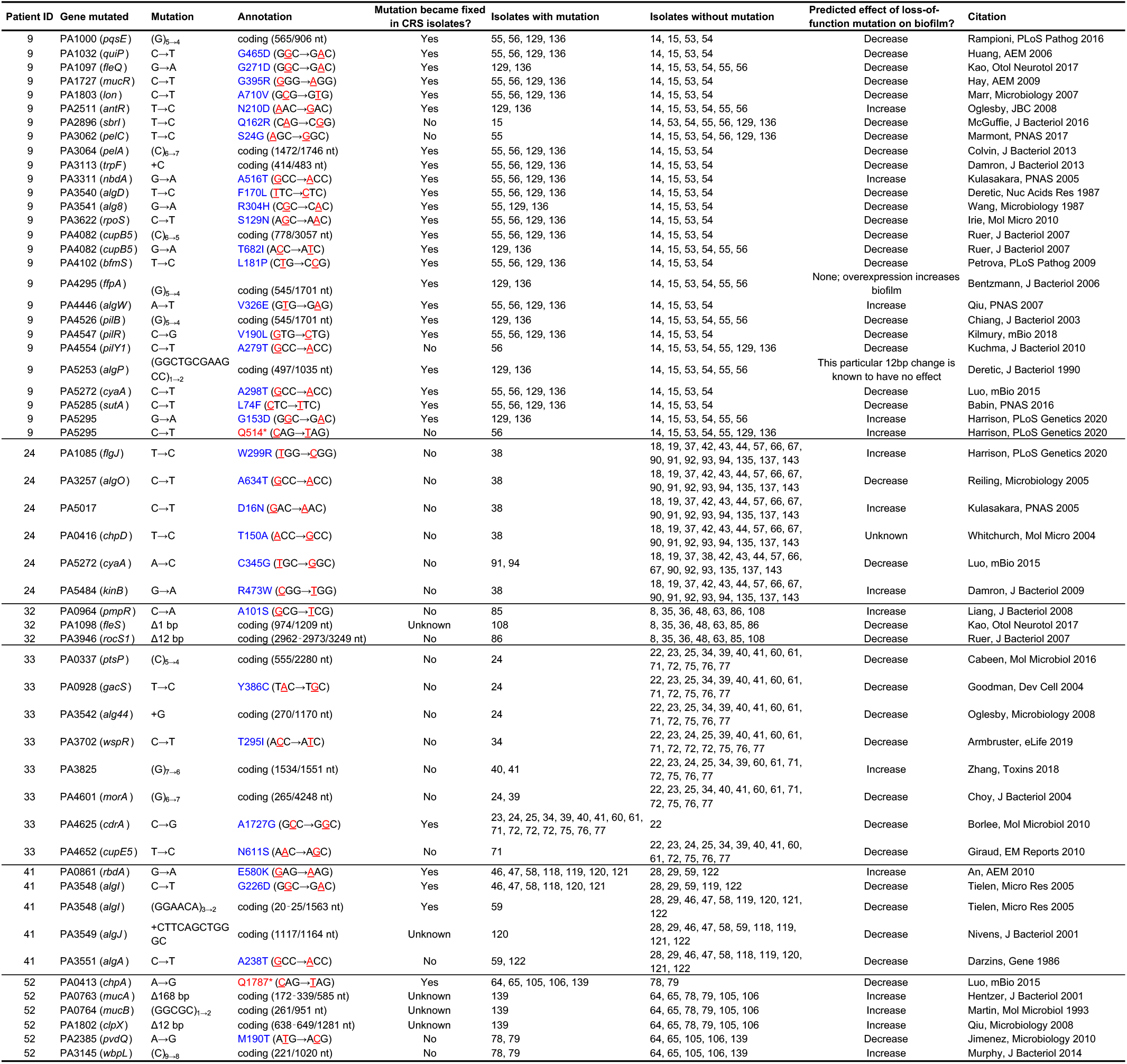
Mutations in biofilm formation genes. Mutation and annotation columns are imported from breseq. The predicted effect of a loss-of-function mutation in each gene is based on the citation(s) listed. Mutations were described as “fixed” in the population if they were present in isolates collected in at least two time points, including the last time point. “Unknown” indicates that it is unknown whether this mutation became fixed because it emerged in an isolate collected on

**Supplemental table 10.**
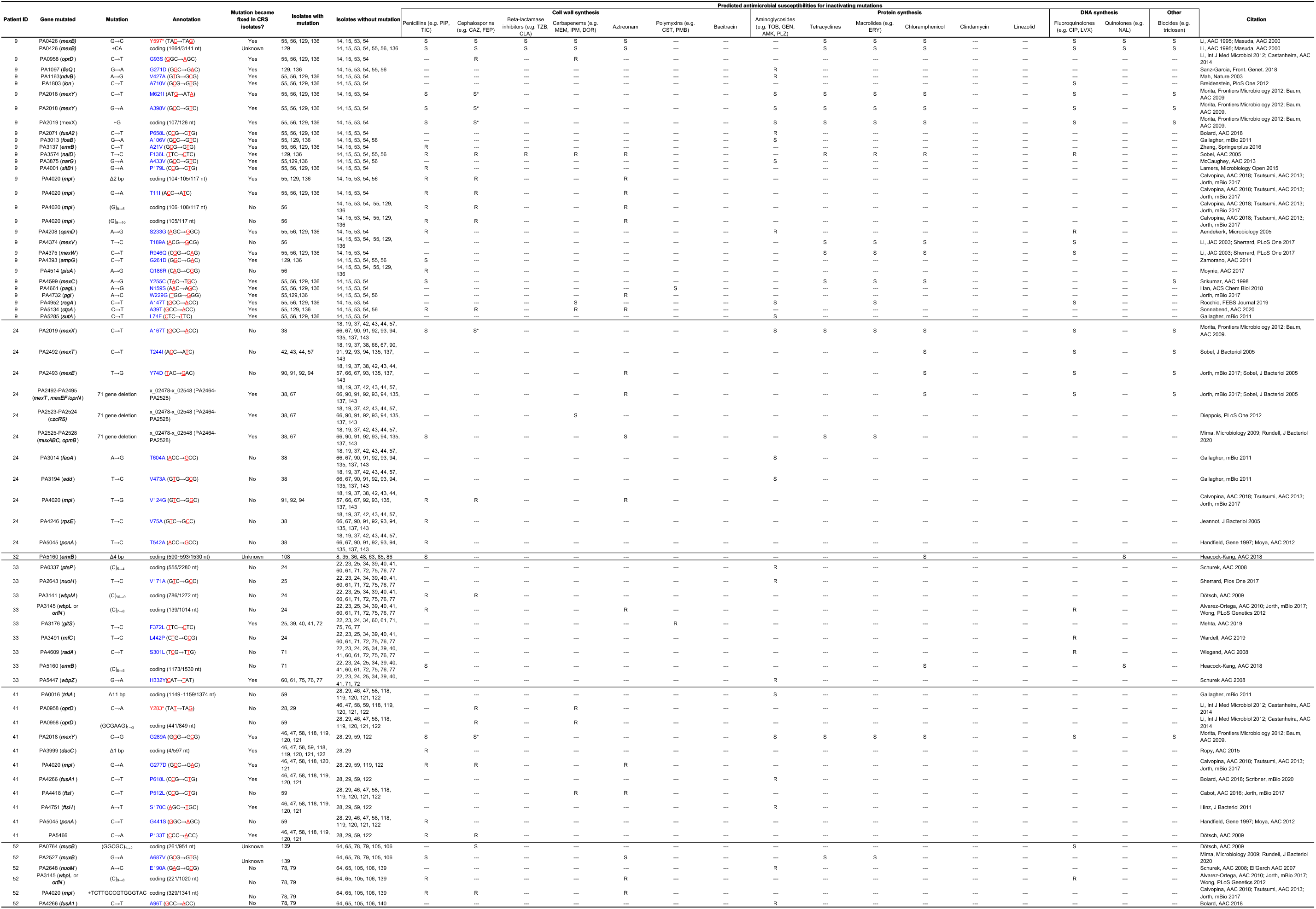
Mutations in antibiotic resistance genes. Mutation and annotation columns are imported from breseq. The predicted effect of a loss-of-function mutation in each gene is based on the citation(s) listed. “R” indicates that a loss-of-function mutation would confer increased resistance to at least one antibiotic in the class listed, whereas “S” indicates that it would confer increased susceptibility. Mutations were described as “fixed” in the population if they were present in isolates collected in at least two time points, including the last time point. “Unknown” indicates that it is unknown whether this mutation became fixed because it emerged in an isolate collected on the last time point for that study participant.

**Supplemental Table 11.** Topical antibiotic or steroid usage among all study participants (including those without longitudinal *P. aeruginosa* isolates).

| Patient # | Systemic or oral antibiotics | Nasal steroids | Topical antibiotics |  |  |  |  |
| --- | --- | --- | --- | --- | --- | --- | --- |
|  |  |  | Any? | Ciprofloxacin | Gentamicin | Mupirocin | Vancomycin |
| 9 | Yes | No | Yes | Yes | No | Yes | No |
| 24 | Yes | No | Yes | No | Yes | No | Yes |
| 32 | Yes | Yes | Yes | Yes | Yes | Yes | No |
| 33 | Yes | Yes | Yes | Yes | No | Yes | No |
| 41 | Yes | No | Yes | No | Yes | No | No |
| 52 | Yes | Yes | Yes | Yes | Yes | Yes | No |
| All other patients (n = 12) | 12/12 | 9/12 | 10/12 | 4/12 | 6/12 | 7/12 | 2/12 |

**Supplemental table 12.**
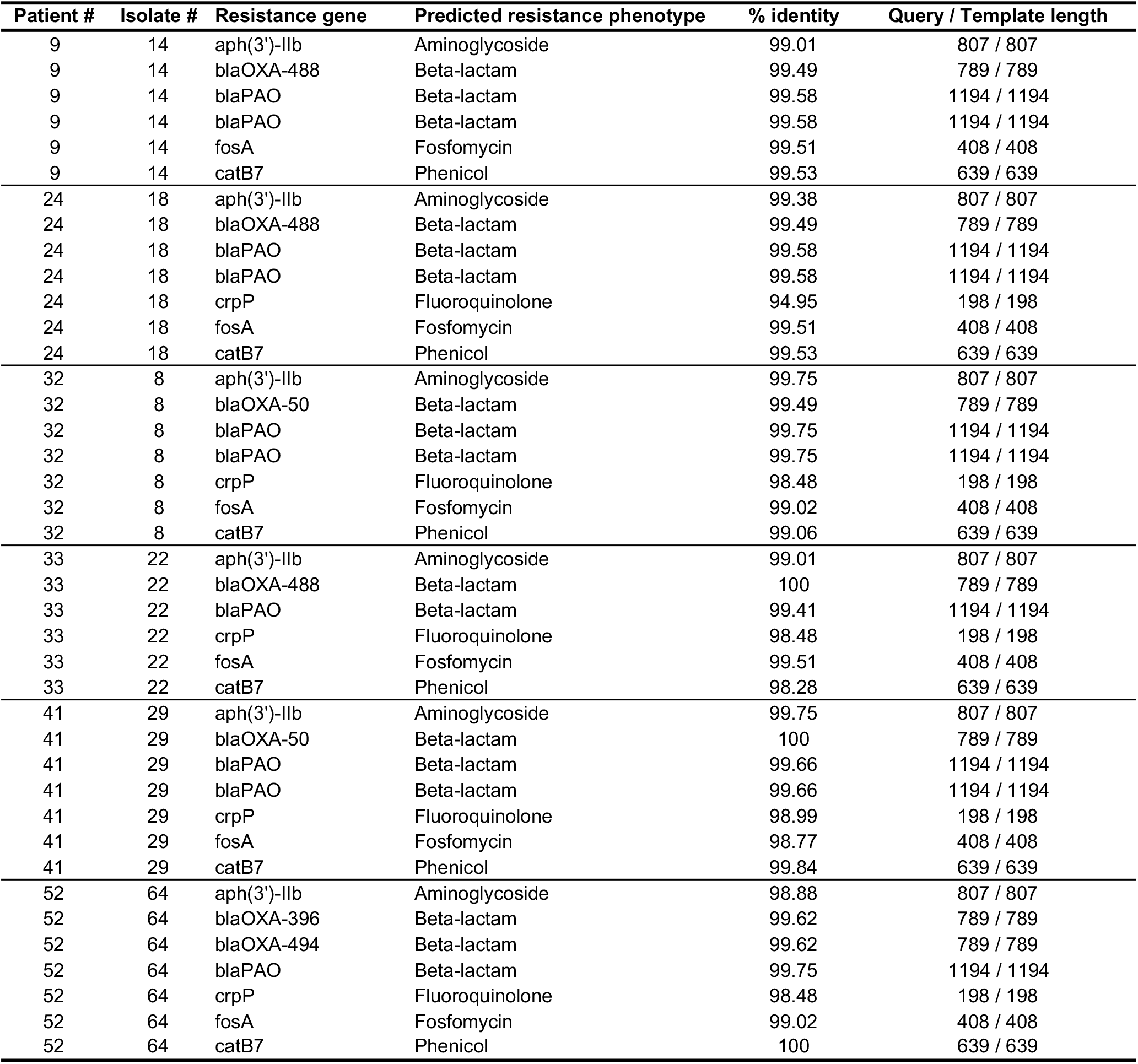
Antimicrobial resistance genes present on the earliest *P. aeruginosa* sinus isolate genome from each of the 6 patients for whom whole genome sequencing was performed. Resistance genes were predicted using ResFinder version 3.2.

## References

1. B. W. Ramsey, Management of pulmonary disease in patients with cystic fibrosis. N Engl J Med. 335, 179–188 (1996).

2. M. S. Muhlebach, B. T. Zorn, C. R. Esther, J. E. Hatch, C. P. Murray, L. Turkovic, S. C. Ranganathan, R. C. Boucher, S. M. Stick, M. C. Wolfgang, Initial acquisition and succession of the cystic fibrosis lung microbiome is associated with disease progression in infants and preschool children. PLoS pathogens. 14, e1006798–e1006798 (2018).

3. P. H. Nielsen, V. B. Rudkjøbing, T. R. Thomsen, K. N. Kragh, M. Givskov, M. Alhede, N. Høiby, T. Bjarnsholt, U. R. Johansen, The microorganisms in chronically infected end-stage and non-end-stage cystic fibrosis patients. FEMS Immunology & Medical Microbiology. 65, 236–244 (2012).

4. S. L. Heltshe, U. Khan, V. Beckett, A. Baines, J. Emerson, D. B. Sanders, R. L. Gibson, W. Morgan, M. Rosenfeld, Longitudinal development of initial, chronic and mucoid Pseudomonas aeruginosa infection in young children with cystic fibrosis. J Cyst Fibros. 17, 341–347 (2018).

5. T. W. R. Lee, K. G. Brownlee, S. P. Conway, M. Denton, J. M. Littlewood, Evaluation of a new definition for chronic Pseudomonas aeruginosa infection in cystic fibrosis patients. Journal of Cystic Fibrosis. 2, 29–34 (2003).

6. C. Winstanley, S. O’Brien, M. A. Brockhurst, Pseudomonas aeruginosa Evolutionary Adaptation and Diversification in Cystic Fibrosis Chronic Lung Infections. Trends in microbiology. 24, 327–337 (2016).

7. J. Diaz Caballero, S. T. Clark, B. Coburn, Y. Zhang, P. W. Wang, S. L. Donaldson, D. E. Tullis, Y. C. W. Yau, V. J. Waters, D. M. Hwang, D. S. Guttman, Selective Sweeps and Parallel Pathoadaptation Drive Pseudomonas aeruginosa Evolution in the Cystic Fibrosis Lung. mBio. 6, e00981 (2015).

8. H. K. Huse, T. Kwon, J. E. A. Zlosnik, D. P. Speert, E. M. Marcotte, M. Whiteley, Parallel Evolution in Pseudomonas aeruginosaover 39,000 Generations In Vivo. mBio. 1, e00199–10 (2010).

9. D. A. D’Argenio, M. Wu, L. R. Hoffman, H. D. Kulasekara, E. Déziel, E. E. Smith, H. Nguyen, R. K. Ernst, T. J. Larson Freeman, D. H. Spencer, M. Brittnacher, H. S. Hayden, S. Selgrade, M. Klausen, D. R. Goodlett, J. L. Burns, B. W. Ramsey, S. I. Miller, Growth phenotypes of Pseudomonas aeruginosa lasR mutants adapted to the airways of cystic fibrosis patients. Molecular Microbiology. 64, 512–533 (2007).

10. N. Mayer-Hamblett, B. W. Ramsey, H. D. Kulasekara, D. J. Wolter, L. S. Houston, C. E. Pope, B. R. Kulasekara, C. R. Armbruster, J. L. Burns, G. Retsch-Bogart, M. Rosenfeld, R. L. Gibson, S. I. Miller, U. Khan, L. R. Hoffman, Pseudomonas aeruginosa phenotypes associated with eradication failure in children with cystic fibrosis. Clin Infect Dis. 59, 624– 631 (2014).

11. M. M. Starkey, J. H. Hickman, L. Ma, N. Zhang, S. De Long, A. Hinz, S. Palacios, C. Manoil, M.J. Kirisits, T. D. Starner, D. J. Wozniak, C. S. Harwood, M. R. Parsek, Pseudomonas aeruginosa Rugose Small-Colony Variants Have Adaptations That Likely Promote Persistence in the Cystic Fibrosis Lung. Journal of Bacteriology. 191, 3492–3503 (2009).

12. N. Mayer-Hamblett, M. Rosenfeld, R. L. Gibson, B. W. Ramsey, H. D. Kulasekara, G. Z. Retsch-Bogart, W. Morgan, D. J. Wolter, C. E. Pope, L. S. Houston, B. R. Kulasekara, U. Khan, J. L. Burns, S. I. Miller, L. R. Hoffman, Pseudomonas aeruginosa In Vitro Phenotypes Distinguish Cystic Fibrosis Infection Stages and Outcomes. American Journal of Respiratory and Critical Care Medicine. 190, 289–297 (2014).

13. D. Derry, Development of mucoid *Pseudomonas aeruginosa* coincides with pulmonary deterioration in cystic fibrosis. Thorax. 60, 334 (2005).

14. M. E. Purdy-Gibson, M. France, T. C. Hundley, N. Eid, S. K. Remold, Pseudomonas aeruginosa in CF and non-CF homes is found predominantly in drains. Journal of Cystic Fibrosis. 14, 341–346 (2015).

15. J. G. Mainz, R. Michl, W. Pfister, J. F. Beck, Cystic Fibrosis Upper Airways Primary Colonization with Pseudomonas aeruginosa: Eradicated by Sinonasal Antibiotic Inhalation. Am J Respir Crit Care Med. 184, 1089–1090 (2011).

16. A. Folkesson, L. Jelsbak, L. Yang, H. K. Johansen, O. Ciofu, N. Hoiby, S. Molin, Adaptation of Pseudomonas aeruginosa to the cystic fibrosis airway: an evolutionary perspective. Nat Rev Micro. 10, 841–851 (2012).

17. S. Boutin, A. H. Dalpke, Acquisition and adaptation of the airway microbiota in the early life of cystic fibrosis patients. Molecular and Cellular Pediatrics. 4, 1 (2017).

18. H. J. C. Bonestroo, K. M. de Winter-de Groot, C. K. van der Ent, H. G. M. Arets, Upper and lower airway cultures in children with cystic fibrosis: do not neglect the upper airways. J Cyst Fibros. 9, 130–134 (2010).

19. S. K. Hansen, M. H. Rau, H. K. Johansen, O. Ciofu, L. Jelsbak, L. Yang, A. Folkesson, H. Ø. Jarmer, K. Aanæs, C. von Buchwald, N. Høiby, S. Molin, Evolution and diversification of Pseudomonas aeruginosa in the paranasal sinuses of cystic fibrosis children have implications for chronic lung infection. The Isme Journal. 6, 31 (2011).

20. K. Aanæs, Bacterial sinusitis can be a focus for initial lung colonisation and chronic lung infection in patients with cystic fibrosis. Journal of Cystic Fibrosis. 12, S1–S20 (2013).

21. M. Alanin, K. Aanaes, N. Høiby, T. Pressler, M. Skov, K. Nielsen, D. Taylor-Robinson, E. Waldmann, H. Krogh Johansen, C. von Buchwald, Sinus surgery postpones chronic Gram-negative lung infection: cohort study of 106 patients with cystic fibrosis. Rhinology. 54, 206–213 (2016).

22. J. G. Mainz, L. Naehrlich, M. Schien, M. Käding, I. Schiller, S. Mayr, G. Schneider, B. Wiedemann, L. Wiehlmann, N. Cramer, W. Pfister, B. C. Kahl, J. F. Beck, B. Tümmler, Concordant genotype of upper and lower airways *P aeruginosa* and *S aureus* isolates in cystic fibrosis. Thorax. 64, 535 (2009).

23. P. Wilson, C. Lambert, S. B. Carr, C. Pao, Paranasal sinus pathogens in children with cystic fibrosis: Do they relate to lower respiratory tract pathogens and is eradication successful? Journal of Cystic Fibrosis. 13, 449–454 (2014).

24. K. L. Whiteson, B. Bailey, M. Bergkessel, D. Conrad, L. Delhaes, B. Felts, J. K. Harris, R. Hunter, Y. W. Lim, H. Maughan, R. Quinn, P. Salamon, J. Sullivan, B. D. Wagner, P. B. Rainey, The Upper Respiratory Tract as a Microbial Source for Pulmonary Infections in Cystic Fibrosis. Parallels from Island Biogeography. Am J Respir Crit Care Med. 189, 1309–1315 (2014).

25. V. G. Gentile, G. Isaacson, Patterns of Sinusitis in Cystic Fibrosis. The Laryngoscope. 106, 1005–1009 (1996).

26. J. M. Robertson, E. M. Friedman, B. K. Rubin, Nasal and sinus disease in cystic fibrosis. Paediatric Respiratory Reviews. 9, 213–219 (2008).

27. S. K. Lucas, R. Yang, J. M. Dunitz, H. C. Boyer, R. C. Hunter, 16S rRNA gene sequencing reveals site-specific signatures of the upper and lower airways of cystic fibrosis patients. Journal of cystic fibrosis : official journal of the European Cystic Fibrosis Society. 17, 204– 212 (2018).

28. S. D. Pletcher, A. N. Goldberg, E. K. Cope, Loss of Microbial Niche Specificity Between the Upper and Lower Airways in Patients With Cystic Fibrosis. The Laryngoscope. 129, 544–550 (2019).

29. E. K. Cope, A. N. Goldberg, S. D. Pletcher, S. V. Lynch, Compositionally and functionally distinct sinus microbiota in chronic rhinosinusitis patients have immunological and clinically divergent consequences. Microbiome. 5, 53–53 (2017).

30. M. R. Chaaban, A. Kejner, S. M. Rowe, B. A. Woodworth, Cystic fibrosis chronic rhinosinusitis: a comprehensive review. American journal of rhinology & allergy. 27, 387– 395 (2013).

31. J. Nelson, P. Karempelis, J. Dunitz, R. Hunter, H. Boyer, Pulmonary aspiration of sinus secretions in patients with cystic fibrosis. International Forum of Allergy & Rhinology. 8, 385–388 (2018).

32. O. Ciofu, H. K. Johansen, K. Aanaes, T. Wassermann, M. Alhede, C. von Buchwald, N. Høiby, P. aeruginosa in the paranasal sinuses and transplanted lungs have similar adaptive mutations as isolates from chronically infected CF lungs. Journal of Cystic Fibrosis. 12, 729–736 (2013).

33. H. K. Johansen, K. Aanaes, T. Pressler, K. G. Nielsen, J. Fisker, M. Skov, N. Høiby, C. von Buchwald, Colonisation and infection of the paranasal sinuses in cystic fibrosis patients is accompanied by a reduced PMN response. Journal of Cystic Fibrosis. 11, 525–531 (2012).

34. S. Walter, P. Gudowius, J. Bosshammer, U. Römling, H. Weissbrodt, W. Schürmann, H. von der Hardt, B. Tümmler, Epidemiology of chronic Pseudomonas aeruginosa infections in the airways of lung transplant recipients with cystic fibrosis. Thorax. 52, 318–321 (1997).

35. J. A. Bartell, L. M. Sommer, J. A. J. Haagensen, A. Loch, R. Espinosa, S. Molin, H. K. Johansen, Evolutionary highways to persistent bacterial infection. Nature Communications. 10, 629 (2019).

36. A. C. Zemke, M. Nouraie, J. Moore, J. R. Gaston, N. Rowan, J. Pilewski, J. M. Bomberger, S. E. Lee, Clinical predictors of cystic fibrosis chronic rhinosinusitis severity (2019).

37. G. B. Rogers, M. P. Carroll, D. J. Serisier, P. M. Hockey, G. Jones, K. D. Bruce, Characterization of Bacterial Community Diversity in Cystic Fibrosis Lung Infections by Use of 16S Ribosomal DNA Terminal Restriction Fragment Length Polymorphism Profiling. J. Clin. Microbiol. 42, 5176 (2004).

38. F. J. Whelan, A. A. Heirali, L. Rossi, H. R. Rabin, M. D. Parkins, M. G. Surette, Longitudinal sampling of the lung microbiota in individuals with cystic fibrosis. PLOS ONE. 12, e0172811 (2017).

39. B. Coburn, P. W. Wang, J. Diaz Caballero, S. T. Clark, V. Brahma, S. Donaldson, Y. Zhang, A. Surendra, Y. Gong, D. Elizabeth Tullis, Y. C. W. Yau, V. J. Waters, D. M. Hwang, D. S. Guttman, Lung microbiota across age and disease stage in cystic fibrosis. Scientific Reports. 5, 10241 (2015).

40. A. A. Heirali, N. Acosta, D. G. Storey, M. L. Workentine, R. Somayaji, I. Laforest-Lapointe, W. Leung, B. S. Quon, Y. Berthiaume, H. R. Rabin, B. J. Waddell, L. Rossi, M. G. Surette, M. D. Parkins, The effects of cycled inhaled aztreonam on the cystic fibrosis (CF) lung microbiome. Journal of Cystic Fibrosis. 18, 829–837 (2019).

41. A. A. Fodor, E. R. Klem, D. F. Gilpin, J. S. Elborn, R. C. Boucher, M. M. Tunney, M. C. Wolfgang, The Adult Cystic Fibrosis Airway Microbiota Is Stable over Time and Infection Type, and Highly Resilient to Antibiotic Treatment of Exacerbations. PLOS ONE. 7, e45001 (2012).

42. M. Hoggard, K. Biswas, M. Zoing, B. Wagner Mackenzie, M. W. Taylor, R. G. Douglas, Evidence of microbiota dysbiosis in chronic rhinosinusitis. International Forum of Allergy & Rhinology. 7, 230–239 (2017).

43. A. W. G. Flight, A. Smith, C. Paisey, J. R. Marchesi, M. J. Bull, P. J. Norville, K. J. Mutton, A.K. Webb, R. J. Bright-Thomas, A. M. Jones, E. Mahenthiralingam, Rapid Detection of Emerging Pathogens and Loss of Microbial Diversity Associated with Severe Lung Disease in Cystic Fibrosis. J Clin Microbiol. 53, 2022–2029 (2015).

44. L. Cuthbertson, A. W. Walker, A. E. Oliver, G. B. Rogers, D. W. Rivett, T. H. Hampton, A. Ashare, J. S. Elborn, A. De Soyza, M. P. Carroll, L. R. Hoffman, C. Lanyon, S. M. Moskowitz, G. A. O’Toole, J. Parkhill, P. J. Planet, C. C. Teneback, M. M. Tunney, J. B. Zuckerman, K. D. Bruce, C. J. van der Gast, Lung function and microbiota diversity in cystic fibrosis. Microbiome. 8, 45 (2020).

45. J. L. Kennedy, M. A. Hubbard, P. Huyett, J. T. Patrie, L. Borish, S. C. Payne, Sino-nasal outcome test (SNOT-22): a predictor of postsurgical improvement in patients with chronic sinusitis. Ann Allergy Asthma Immunol. 111, 246–251.e2 (2013).

46. A. J. Psaltis, G. Li, R. Vaezeafshar, K.-S. Cho, P. H. Hwang, Modification of the lund-kennedy endoscopic scoring system improves its reliability and correlation with patient-reported outcome measures. The Laryngoscope. 124, 2216–2223 (2014).

47. L. Freschi, J. Jeukens, I. Kukavica-Ibrulj, B. Boyle, M.-J. Dupont, J. Laroche, S. Larose, H. Maaroufi, J. L. Fothergill, M. Moore, G. L. Winsor, S. D. Aaron, J. Barbeau, S. C. Bell, J. L. Burns, M. Camara, A. Cantin, S. J. Charette, K. Dewar, É. Déziel, K. Grimwood, R. E. W. Hancock, J. J. Harrison, S. Heeb, L. Jelsbak, B. Jia, D. T. Kenna, T. J. Kidd, J. Klockgether, J. S. Lam, I. L. Lamont, S. Lewenza, N. Loman, F. Malouin, J. Manos, A. G. McArthur, J. McKeown, J. Milot, H. Naghra, D. Nguyen, S. K. Pereira, G. G. Perron, J.-P. Pirnay, P. B. Rainey, S. Rousseau, P. M. Santos, A. Stephenson, V. Taylor, J. F. Turton, N. Waglechner, P. Williams, S. W. Thrane, G. D. Wright, F. S. L. Brinkman, N. P. Tucker, B. Tümmler, C. Winstanley, R. C. Levesque, Clinical utilization of genomics data produced by the international Pseudomonas aeruginosa consortium. Frontiers in Microbiology. 6, 1036 (2015).

48. M. R. Hendricks, L. P. Lashua, D. K. Fischer, B. A. Flitter, K. M. Eichinger, J. E. Durbin, S. N. Sarkar, C. B. Coyne, K. M. Empey, J. M. Bomberger, Respiratory syncytial virus infection enhances *Pseudomonas aeruginosa* biofilm growth through dysregulation of nutritional immunity. Proc Natl Acad Sci USA. 113, 1642 (2016).

49. B. W. Holloway, Genetic Recombination in Pseudomonas aeruginosa. Microbiology. 13 (1955), pp. 572–581.

50. E. J. Feil, J. E. Cooper, H. Grundmann, D. A. Robinson, M. C. Enright, T. Berendt, S. J. Peacock, J. M. Smith, M. Murphy, B. G. Spratt, C. E. Moore, N. P. J. Day, How Clonal Is *Staphylococcus aureus*? J. Bacteriol. 185, 3307 (2003).

51. A. Oliver, R. Cantón, P. Campo, F. Baquero, J. Blázquez, High Frequency of Hypermutable *Pseudomonas aeruginosa* in Cystic Fibrosis Lung Infection. Science. 288, 1251 (2000).

52. R. L. Marvig, H. K. Johansen, S. Molin, L. Jelsbak, Genome Analysis of a Transmissible Lineage of Pseudomonas aeruginosa Reveals Pathoadaptive Mutations and Distinct Evolutionary Paths of Hypermutators. PLOS Genetics. 9, e1003741 (2013).

53. C. A. Colque, A. G. Albarracín Orio, S. Feliziani, R. L. Marvig, A. R. Tobares, H. K. Johansen, S. Molin, A. M. Smania, Hypermutator *Pseudomonas* aeruginosa Exploits Multiple Genetic Pathways To Develop Multidrug Resistance during Long-Term Infections in the Airways of Cystic Fibrosis Patients. Antimicrob. Agents Chemother. 64, e02142–19 (2020).

54. O. Ciofu, B. Riis, T. Pressler, H. E. Poulsen, N. Høiby, Occurrence of Hypermutable *Pseudomonas aeruginosa* in Cystic Fibrosis Patients Is Associated with the Oxidative Stress Caused by Chronic Lung Inflammation. Antimicrob. Agents Chemother. 49, 2276 (2005).

55. M. Hogardt, C. Hoboth, S. Schmoldt, C. Henke, L. Bader, J. Heesemann, Stage-Specific Adaptation of Hypermutable Pseudomonas aeruginosa Isolates during Chronic Pulmonary Infection in Patients with Cystic Fibrosis. The Journal of Infectious Diseases. 195, 70–80 (2007).

56. S. Montanari, A. Oliver, P. Salerno, A. Mena, G. Bertoni, B. Tümmler, L. Cariani, M. Conese, G. Döring, A. Bragonzi, Biological cost of hypermutation in Pseudomonas aeruginosa strains from patients with cystic fibrosis. Microbiology. 153 (2007), pp. 1445– 1454.

57. A. Oliver, Mutators in cystic fibrosis chronic lung infection: Prevalence, mechanisms, and consequences for antimicrobial therapy. International Journal of Medical Microbiology. 300, 563–572 (2010).

58. L. H. Sanders, A. Rockel, H. Lu, D. J. Wozniak, M. D. Sutton, Role of *Pseudomonas aeruginosa dinB*-Encoded DNA Polymerase IV in Mutagenesis. J. Bacteriol. 188, 8573 (2006).

59. J. Vandecraen, M. Chandler, A. Aertsen, R. Van Houdt, The impact of insertion sequences on bacterial genome plasticity and adaptability. Critical Reviews in Microbiology. 43, 709– 730 (2017).

60. E. Sentausa, P. Basso, A. Berry, A. Adrait, G. Bellement, Y. Couté, S. Lory, S. Elsen, I. Attrée, Insertion sequences drive the emergence of a highly adapted human pathogen. Microbial Genomics (2019), doi:https://doi.org/10.1099/mgen.0.000265.

61. J. C. Evans, H. Segal, A Novel Insertion Sequence, IS *Pa26*, in *oprD* of *Pseudomonas aeruginosa* Is Associated with Carbapenem Resistance. Antimicrob. Agents Chemother. 51, 3776 (2007).

62. D. J. Wolter, N. D. Hanson, P. D. Lister, Insertional inactivation of oprD in clinical isolates of Pseudomonas aeruginosa leading to carbapenem resistance. FEMS Microbiology Letters. 236, 137–143 (2004).

63. A. U. Kresse, H. Blöcker, U. Römling, ISPa20 advances the individual evolution of Pseudomonas aeruginosa clone C subclone C13 strains isolated from cystic fibrosis patients by insertional mutagenesis and genomic rearrangements. Archives of Microbiology. 185, 245–254 (2006).

64. D. A. D’Argenio, M. Wu, L. R. Hoffman, H. D. Kulasekara, E. Déziel, E. E. Smith, H. Nguyen, R. K. Ernst, T. J. Larson Freeman, D. H. Spencer, M. Brittnacher, H. S. Hayden, S. Selgrade, M. Klausen, D. R. Goodlett, J. L. Burns, B. W. Ramsey, S. I. Miller, Growth phenotypes of Pseudomonas aeruginosa lasR mutants adapted to the airways of cystic fibrosis patients. Molecular Microbiology. 64, 512–533 (2007).

65. D. Papadopoulos, D. Schneider, J. Meier-Eiss, W. Arber, R. E. Lenski, M. Blot, Genomic evolution during a 10,000-generation experiment with bacteria. Proc Natl Acad Sci USA. 96, 3807 (1999).

66. O. Tenaillon, J. E. Barrick, N. Ribeck, D. E. Deatherage, J. L. Blanchard, A. Dasgupta, G. C. Wu, S. Wielgoss, S. Cruveiller, C. Médigue, D. Schneider, R. E. Lenski, Tempo and mode of genome evolution in a 50,000-generation experiment. Nature. 536, 165–170 (2016).

67. A. U. Kresse, S. D. Dinesh, K. Larbig, U. Römling, Impact of large chromosomal inversions on the adaptation and evolution of Pseudomonas aeruginosa chronically colonizing cystic fibrosis lungs. Molecular Microbiology. 47, 145–158 (2003).

68. M. Vareille, E. Kieninger, M. R. Edwards, N. Regamey, The airway epithelium: soldier in the fight against respiratory viruses. Clinical microbiology reviews. 24, 210–229 (2011).

69. A. Licari, R. Castagnoli, C. F. Denicolò, L. Rossini, A. Marseglia, G. L. Marseglia, The Nose and the Lung: United Airway Disease? Frontiers in Pediatrics. 5, 44 (2017).

70. A. Kicic, E. de Jong, K.-M. Ling, K. Nichol, D. Anderson, P. A. B. Wark, D. A. Knight, A. Bosco, S. M. Stick, A. Kicic, S. M. Stick, D. A. Knight, E. Kicic-Starcevich, L. W. Garratt, M. Padros-Goosen, E.-L. Tan, E. N. Sutanto, K. Looi, J. Hillas, T. Iosifidis, N. C. Shaw, S. T. Montgomery, K.-M. Ling, K. M. Martinovich, F. J. Lannigan, R. Bergesio, B. Lee, S. Vijaya-Sekeran, P. Swan, M. Heaney, I. Forsyth, T. Schoep, A. Larcombe, M. Hunter, K. McGee, N. Millington, A. Kicic, S. M. Stick, E. Kicic-Starcevich, L. W. Garratt, E. N. Sutanto, K. Looi, J. Hillas, T. Iosifidis, N. C. Shaw, S. T. Montgomery, K.-M. Ling, K. M. Martinovich, M. W.-P. Poh, D. R. Laucirica, C. Schofield, S. McLean, K. Landwehr, E. de Jong, N. Farrow, E. Roscioli, D. Parsons, D. A. Knight, C. Grainge, A. T. Reid, S.-L. Loo, P. C. Veerati, Assessing the unified airway hypothesis in children via transcriptional profiling of the airway epithelium. Journal of Allergy and Clinical Immunology. 145, 1562– 1573 (2020).

71. J. H. Krouse, R. W. Brown, S. M. Fineman, J. K. Han, A. J. Heller, S. Joe, H. J. Krouse, H. C. Pillsbury, M. W. Ryan, M. C. Veling, Asthma and the unified airway. Otolaryngol Head Neck Surg. 136, S75–S106 (2007).

72. M. R. Scribner, A. Santos-Lopez, C. W. Marshall, C. Deitrick, V. S. Cooper, Parallel Evolution of Tobramycin Resistance across Species and Environments. mBio. 11, e00932–20 (2020).

73. J. P. McCutcheon, N. A. Moran, Extreme genome reduction in symbiotic bacteria. Nature Reviews Microbiology. 10, 13–26 (2012).

74. N. A. Moran, G. R. Plague, Genomic changes following host restriction in bacteria. Current Opinion in Genetics & Development. 14, 627–633 (2004).

75. X.-J. Yu, D. H. Walker, Y. Liu, L. Zhang, Amino acid biosynthesis deficiency in bacteria associated with human and animal hosts. Infect Genet Evol. 9, 514–517 (2009).

76. L. Rahme, E. Stevens, S. Wolfort, J. Shao, R. Tompkins, F. Ausubel, Common virulence factors for bacterial pathogenicity in plants and animals. Science. 268, 1899 (1995).

77. K. Cheng, R. L. Smyth, J. R. Govan, C. Doherty, C. Winstanley, N. Denning, D. P. Heaf, H. van Saene, C. A. Hart, Spread of β-lactam-resistant Pseudomonas aeruginosa in a cystic fibrosis clinic. The Lancet. 348, 639–642 (1996).

78. E. Lerat, H. Ochman, Recognizing the pseudogenes in bacterial genomes. Nucleic Acids Res. 33, 3125–3132 (2005).

79. C.-H. Kuo, H. Ochman, The extinction dynamics of bacterial pseudogenes. PLoS Genet. 6, e1001050 (2010).

80. M Syberg-Olsen, A. Garber, P. Keeling, J. McCutcheon, Pseudofinder (GitHub repository, 2020; https://github.com/filip-husnik/pseudofinder/).

81. G. R. Burke, N. A. Moran, Massive genomic decay in Serratia symbiotica, a recently evolved symbiont of aphids. Genome Biol Evol. 3, 195–208 (2011).

82. V. S. Cooper, D. Schneider, M. Blot, R. E. Lenski, Mechanisms causing rapid and parallel losses of ribose catabolism in evolving populations of Escherichia coli B. J Bacteriol. 183, 2834–2841 (2001).

83. B. R. Boles, P. K. Singh, Endogenous oxidative stress produces diversity and adaptability in biofilm communities. Proc Natl Acad Sci USA. 105, 12503 (2008).

84. C. Hoboth, R. Hoffmann, A. Eichner, C. Henke, S. Schmoldt, A. Imhof, J. Heesemann, M. Hogardt, Dynamics of Adaptive Microevolution of Hypermutable Pseudomonas aeruginosa during Chronic Pulmonary Infection in Patients with Cystic Fibrosis. The Journal of Infectious Diseases. 200, 118–130 (2009).

85. D. J. Waine, D. Honeybourne, E. G. Smith, J. L. Whitehouse, C. G. Dowson, Association between Hypermutator Phenotype, Clinical Variables, Mucoid Phenotype, and Antimicrobial Resistance in *Pseudomonas aeruginosa*. J. Clin. Microbiol. 46, 3491 (2008).

86. Y. Raynes, P. D. Sniegowski, Experimental evolution and the dynamics of genomic mutation rate modifiers. Heredity. 113, 375–380 (2014).

87. A. Oliver, A. Mena, Bacterial hypermutation in cystic fibrosis, not only for antibiotic resistance. Clinical Microbiology and Infection. 16, 798–808 (2010).

88. N. D. Chu, S. A. Clarke, S. Timberlake, M. F. Polz, A. D. Grossman, E. J. Alm, A Mobile Element in mutS Drives Hypermutation in a Marine Vibrio. mBio. 8, e02045–16 (2017).

89. M. A. Campbell, P. Łukasik, C. Simon, J. P. McCutcheon, Idiosyncratic Genome Degradation in a Bacterial Endosymbiont of Periodical Cicadas. Current Biology. 27, 3568–3575.e3 (2017).

90. V. Merhej, M. Royer-Carenzi, P. Pontarotti, D. Raoult, Massive comparative genomic analysis reveals convergent evolution of specialized bacteria. Biol Direct. 4, 13–13 (2009).

91. J. Wei, M. B. Goldberg, V. Burland, M. M. Venkatesan, W. Deng, G. Fournier, G. F. Mayhew, G. Plunkett 3rd, D. J. Rose, A. Darling, B. Mau, N. T. Perna, S. M. Payne, L. J. Runyen-Janecky, S. Zhou, D. C. Schwartz, F. R. Blattner, Complete genome sequence and comparative genomics of Shigella flexneri serotype 2a strain 2457T. Infect Immun. 71, 2775–2786 (2003).

92. J. Parkhill, M. Sebaihia, A. Preston, L. D. Murphy, N. Thomson, D. E. Harris, M. T. G. Holden, C. M. Churcher, S. D. Bentley, K. L. Mungall, A. M. Cerdeño-Tárraga, L. Temple, K. James, B. Harris, M. A. Quail, M. Achtman, R. Atkin, S. Baker, D. Basham, N. Bason, I. Cherevach, T. Chillingworth, M. Collins, A. Cronin, P. Davis, J. Doggett, T. Feltwell, A. Goble, N. Hamlin, H. Hauser, S. Holroyd, K. Jagels, S. Leather, S. Moule, H. Norberczak, S. O’Neil, D. Ormond, C. Price, E. Rabbinowitsch, S. Rutter, M. Sanders, D. Saunders, K. Seeger, S. Sharp, M. Simmonds, J. Skelton, R. Squares, S. Squares, K. Stevens, L. Unwin, S. Whitehead, B. G. Barrell, D. J. Maskell, Comparative analysis of the genome sequences of Bordetella pertussis, Bordetella parapertussis and Bordetella bronchiseptica. Nature Genetics. 35, 32–40 (2003).

93. F. A. Stressmann, G. B. Rogers, P. Marsh, A. K. Lilley, T. W. V. Daniels, M. P. Carroll, L. R. Hoffman, G. Jones, C. E. Allen, N. Patel, B. Forbes, A. Tuck, K. D. Bruce, Does bacterial density in cystic fibrosis sputum increase prior to pulmonary exacerbation? Journal of Cystic Fibrosis. 10, 357–365 (2011).

94. P. Jorth, B. J. Staudinger, X. Wu, K. B. Hisert, H. Hayden, J. Garudathri, C. L. Harding, M. C. Radey, A. Rezayat, G. Bautista, W. R. Berrington, A. F. Goddard, C. Zheng, A. Angermeyer, M. J. Brittnacher, J. Kitzman, J. Shendure, C. L. Fligner, J. Mittler, M. L. Aitken, C. Manoil, J. E. Bruce, T. L. Yahr, P. K. Singh, Regional Isolation Drives Bacterial Diversification within Cystic Fibrosis Lungs. Cell Host Microbe. 18, 307–319 (2015).

95. T. Bjarnsholt, M. Alhede, M. Alhede, S. R. Eickhardt-Sørensen, C. Moser, M. Kühl, P. Ø. Jensen, N. Høiby, The in vivo biofilm. Trends in Microbiology. 21, 466–474 (2013).

96. S. E. Darch, O. Simoska, M. Fitzpatrick, J. P. Barraza, K. J. Stevenson, R. T. Bonnecaze, J. B. Shear, M. Whiteley, Spatial determinants of quorum signaling in a Pseudomonas aeruginosa infection model. Proc Natl Acad Sci U S A. 115, 4779–4784 (2018).

97. E. E. Smith, D. G. Buckley, Z. Wu, C. Saenphimmachak, L. R. Hoffman, D. A. D’Argenio, S. I. Miller, B. W. Ramsey, D. P. Speert, S. M. Moskowitz, J. L. Burns, R. Kaul, M. V. Olson, Genetic adaptation by *Pseudomonas aeruginosa* to the airways of cystic fibrosis patients. Proc Natl Acad Sci USA. 103, 8487 (2006).

98. A. T. Nguyen, M. J. O’Neill, A. M. Watts, C. L. Robson, I. L. Lamont, A. Wilks, A. G. Oglesby-Sherrouse, Adaptation of Iron Homeostasis Pathways by a *Pseudomonas aeruginosa* Pyoverdine Mutant in the Cystic Fibrosis Lung. J. Bacteriol. 196, 2265 (2014).

99. S. O’Brien, D. Williams, J. L. Fothergill, S. Paterson, C. Winstanley, M. A. Brockhurst, High virulence sub-populations in Pseudomonas aeruginosa long-term cystic fibrosis airway infections. BMC Microbiology. 17, 30 (2017).

100. R. C. Hunter, F. Asfour, J. Dingemans, B. L. Osuna, T. Samad, A. Malfroot, P. Cornelis, D. K. Newman, Ferrous iron is a significant component of bioavailable iron in cystic fibrosis airways. MBio. 4, e00557–13 (2013).

101. P. Cornelis, J. Dingemans, Pseudomonas aeruginosa adapts its iron uptake strategies in function of the type of infections. Frontiers in cellular and infection microbiology. 3, 75–75 (2013).

102. C.-Y. Chang, Surface Sensing for Biofilm Formation in Pseudomonas aeruginosa. Front Microbiol. 8, 2671–2671 (2018).

103. M. F. Moradali, S. Ghods, B. H. A. Rehm, Pseudomonas aeruginosa Lifestyle: A Paradigm for Adaptation, Survival, and Persistence. Frontiers in Cellular and Infection Microbiology. 7, 39 (2017).

104. S. Furukawa, S. L. Kuchma, G. A. O’Toole, Keeping Their Options Open: Acute versus Persistent Infections. J. Bacteriol. 188, 1211 (2006).

105. A. T. Y. Yeung, E. C. W. Torfs, F. Jamshidi, M. Bains, I. Wiegand, R. E. W. Hancock, J. Overhage, Swarming of Pseudomonas aeruginosa Is Controlled by a Broad Spectrum of Transcriptional Regulators, Including MetR. J. Bacteriol. 191, 5592 (2009).

106. S. K. Arora, B. W. Ritchings, E. C. Almira, S. Lory, R. Ramphal, A transcriptional activator, FleQ, regulates mucin adhesion and flagellar gene expression in Pseudomonas aeruginosa in a cascade manner. J. Bacteriol. 179, 5574 (1997).

107. I. D. Hay, U. Remminghorst, B. H. A. Rehm, MucR, a Novel Membrane-Associated Regulator of Alginate Biosynthesis in *Pseudomonas aeruginosa*. Appl. Environ. Microbiol. 75, 1110 (2009).

108. B. R. Kulasekara, C. Kamischke, H. D. Kulasekara, M. Christen, P. A. Wiggins, S. I. Miller, c-di-GMP heterogeneity is generated by the chemotaxis machinery to regulate flagellar motility. eLife. 2, e01402 (2013).

109. S. L. Kuchma, A. E. Ballok, J. H. Merritt, J. H. Hammond, W. Lu, J. D. Rabinowitz, G. A. O’Toole, Cyclic-di-GMP-Mediated Repression of Swarming Motility by Pseudomonas aeruginosa: the pilY1 Gene and Its Impact on Surface-Associated Behaviors. Journal of Bacteriology. 192, 2950–2964 (2010).

110. G. W. Lau, H. Ran, F. Kong, D. J. Hassett, D. Mavrodi, *Pseudomonas aeruginosa* Pyocyanin Is Critical for Lung Infection in Mice. Infect. Immun. 72, 4275 (2004).

111. J. M. Farrow 3rd, Z. M. Sund, M. L. Ellison, D. S. Wade, J. P. Coleman, E. C. Pesci, PqsE functions independently of PqsR-Pseudomonas quinolone signal and enhances the rhl quorum-sensing system. J Bacteriol. 190, 7043–7051 (2008).

112. J. J. Harrison, H. Almblad, Y. Irie, D. J. Wolter, H. C. Eggleston, T. E. Randall, J. O. Kitzman, B. Stackhouse, J. C. Emerson, S. Mcnamara, T. J. Larsen, J. Shendure, L. R. Hoffman, D. J. Wozniak, M. R. Parsek, Elevated exopolysaccharide levels in Pseudomonas aeruginosa flagellar mutants have implications for biofilm growth and chronic infections. PLOS Genetics. 16, e1008848 (2020).

113. R. G. Doggett, Incidence of mucoid Pseudomonas aeruginosa from clinical sources. Appl Microbiol. 18, 936–937 (1969).

114. N. Høiby, PREVALENCE OF MUCOID STRAINS OF PSEUDOMONAS AERUGINOSA IN BACTERIOLOGICAL SPECIMENS FROM PATIENTS WITH CYSTIC FIBROSIS AND PATIENTS WITH OTHER DISEASES. Acta Pathologica Microbiologica Scandinavica Section B Microbiology. 83B, 549–552 (1975).

115. O. Ciofu, B. Lee, M. Johannesson, N. O. Hermansen, P. Meyer, N. Høiby, Investigation of the algT operon sequence in mucoid and non-mucoid Pseudomonas aeruginosa isolates from 115 Scandinavian patients with cystic fibrosis and in 88 in vitro non-mucoid revertants. Microbiology. 154, 103–113 (2008).

116. Y. Tsutsumi, H. Tomita, K. Tanimoto, Identification of novel genes responsible for overexpression of ampC in Pseudomonas aeruginosa PAO1. Antimicrob Agents Chemother. 57, 5987–5993 (2013).

117. T.-F. Mah, B. Pitts, B. Pellock, G. C. Walker, P. S. Stewart, G. A. O’Toole, A genetic basis for Pseudomonas aeruginosa biofilm antibiotic resistance. Nature. 426, 306–310 (2003).

118. A. Bolard, P. Plésiat, K. Jeannot, Mutations in Gene fusA1 as a Novel Mechanism of Aminoglycoside Resistance in Clinical Strains of Pseudomonas aeruginosa. Antimicrob Agents Chemother. 62, e01835–17 (2018).

119. S. Aendekerk, S. P. Diggle, Z. Song, N. Høiby, P. Cornelis, P. Williams, M. Cámara, The MexGHI-OpmD multidrug efflux pump controls growth, antibiotic susceptibility and virulence in Pseudomonas aeruginosa via 4-quinolone-dependent cell-to-cell communication. Microbiology. 151 (2005), pp. 1113–1125.

120. F. Sanz-García, S. Hernando-Amado, J. L. Martínez, Mutational Evolution of Pseudomonas aeruginosa Resistance to Ribosome-Targeting Antibiotics. Frontiers in Genetics. 9, 451 (2018).

121. K. N. Schurek, A. K. Marr, P. K. Taylor, I. Wiegand, L. Semenec, B. K. Khaira, R. E. W. Hancock, Novel Genetic Determinants of Low-Level Aminoglycoside Resistance in *Pseudomonas aeruginosa*. Antimicrob. Agents Chemother. 52, 4213 (2008).

122. L. J. Sherrard, A. S. Tai, B. A. Wee, K. A. Ramsay, T. J. Kidd, N. L. Ben Zakour, D. M. Whiley, S. A. Beatson, S. C. Bell, Within-host whole genome analysis of an antibiotic resistant Pseudomonas aeruginosa strain sub-type in cystic fibrosis. PLOS ONE. 12, e0172179 (2017).

123. M. L. Sobel, S. Neshat, K. Poole, Mutations in PA2491 (*mexS*) Promote MexT-Dependent *mexEF-oprN* Expression and Multidrug Resistance in a Clinical Strain of *Pseudomonas aeruginosa*. J. Bacteriol. 187, 1246 (2005).

124. K. Calvopiña, M. B. Avison, Disruption of *mpl* Activates β-Lactamase Production in *Stenotrophomonas maltophilia* and *Pseudomonas aeruginosa* Clinical Isolates. Antimicrob. Agents Chemother. 62, e00638–18 (2018).

125. N. Masuda, E. Sakagawa, S. Ohya, N. Gotoh, H. Tsujimoto, T. Nishino, Substrate specificities of MexAB-OprM, MexCD-OprJ, and MexXY-oprM efflux pumps in Pseudomonas aeruginosa. Antimicrob Agents Chemother. 44, 3322–3327 (2000).

126. J. R. Marchesi, T. Sato, A. J. Weightman, T. A. Martin, J. C. Fry, S. J. Hiom, D. Dymock, W. G. Wade, Design and evaluation of useful bacterium-specific PCR primers that amplify genes coding for bacterial 16S rRNA. Appl Environ Microbiol. 64, 795–799 (1998).

127. E. Mahenthiralingam, M. E. Campbell, J. Foster, J. S. Lam, D. P. Speert, Random amplified polymorphic DNA typing of Pseudomonas aeruginosa isolates recovered from patients with cystic fibrosis. J Clin Microbiol. 34, 1129–1135 (1996).

128. T. Köhler, L. K. Curty, F. Barja, C. van Delden, J. C. Pechère, Swarming of Pseudomonas aeruginosa is dependent on cell-to-cell signaling and requires flagella and pili. J Bacteriol. 182, 5990–5996 (2000).

129. S. C. Woolwine, D. J. Wozniak, Identification of an *Escherichia coli pepA* Homolog and Its Involvement in Suppression of the *algB* Phenotype in Mucoid *Pseudomonas aeruginosa*. J. Bacteriol. 181, 107 (1999).

130. H. J. vogel, D. M. Bonner, Acetylornithinase of Escherichia coli: partial purification and some properties. J Biol Chem. 218, 97–106 (1956).

131. Z. T. Güvener, C. S. Harwood, Subcellular location characteristics of the Pseudomonas aeruginosa GGDEF protein, WspR, indicate that it produces cyclic-di-GMP in response to growth on surfaces. Molecular Microbiology. 66, 1459–1473 (2007).

132. H. G. Riggs Jr, E. D. Rosenblum, Transfection of lysostaphin-treated cells of Staphylococcus aureus. J Virol. 3, 33–37 (1969).

133. E. Déziel, Y. Comeau, R. Villemur, Initiation of biofilm formation by Pseudomonas aeruginosa 57RP correlates with emergence of hyperpiliated and highly adherent phenotypic variants deficient in swimming, swarming, and twitching motilities. J Bacteriol. 183, 1195–1204 (2001).

134. O. Tange, GNU Parallel - The Command-Line Power Tool.; login: The USENIX Magazine. 42–47 (2011).

135. A. J. Page, C. A. Cummins, M. Hunt, V. K. Wong, S. Reuter, M. T. G. Holden, M. Fookes, D. Falush, J. A. Keane, J. Parkhill, Roary: rapid large-scale prokaryote pan genome analysis. Bioinformatics. 31, 3691–3693 (2015).

136. Z. Yang, PAML 4: Phylogenetic Analysis by Maximum Likelihood. Molecular Biology and Evolution. 24, 1586–1591 (2007).

137. C. Camacho, G. Coulouris, V. Avagyan, N. Ma, J. Papadopoulos, K. Bealer, T. L. Madden, BLAST+: architecture and applications. BMC Bioinformatics. 10, 421 (2009).

138. B. Buchfink, C. Xie, D. H. Huson, Fast and sensitive protein alignment using DIAMOND. Nature Methods. 12, 59–60 (2015).

139. R. C. Edgar, MUSCLE: multiple sequence alignment with high accuracy and high throughput. Nucleic Acids Research. 32, 1792–1797 (2004).

140. R Core Team, A Language and Environment for Statistical Computing. *R Foundation for Statistical Computing*, Vienna. (2013) (available at http://www.R-project.org/).

141. A. I. Garber, K. H. Nealson, A. Okamoto, S. M. McAllister, C. S. Chan, R. A. Barco, N. Merino, FeGenie: A Comprehensive Tool for the Identification of Iron Genes and Iron Gene Neighborhoods in Genome and Metagenome Assemblies. Frontiers in Microbiology. 11, 37 (2020).

142. H. Pagès, R. Aboyoun, R. Gentleman, S. Debroy, Biostrings: Efficient manipulation of biological strings.

143. T. D. Lieberman, K. B. Flett, I. Yelin, T. R. Martin, A. J. McAdam, G. P. Priebe, R. Kishony, Genetic variation of a bacterial pathogen within individuals with cystic fibrosis provides a record of selective pressures. Nature Genetics. 46, 82–87 (2014).

144. P. Visca, G. Colotti, L. Serino, D. Verzili, N. Orsi, E. Chiancone, Metal regulation of siderophore synthesis in Pseudomonas aeruginosa and functional effects of siderophore-metal complexes. Appl Environ Microbiol. 58, 2886–2893 (1992).

145. D. W. Essar, L. Eberly, A. Hadero, I. P. Crawford, Identification and characterization of genes for a second anthranilate synthase in Pseudomonas aeruginosa: interchangeability of the two anthranilate synthases and evolutionary implications. J Bacteriol. 172, 884–900 (1990).

146. R. L. Marvig, L. M. Sommer, S. Molin, H. K. Johansen, Convergent evolution and adaptation of Pseudomonas aeruginosa within patients with cystic fibrosis. Nature Genetics. 47, 57–64 (2015).

147. K. E. Huus, J. Joseph, L. Zhang, A. Wong, S. D. Aaron, T.-F. Mah, S. Sad, Clinical Isolates of *Pseudomonas aeruginosa* from Chronically Infected Cystic Fibrosis Patients Fail To Activate the Inflammasome during Both Stable Infection and Pulmonary Exacerbation. J. Immunol., 1501642 (2016).

148. U. A. Ochsner, M. L. Vasil, Gene repression by the ferric uptake regulator in Pseudomonas aeruginosa: cycle selection of iron-regulated genes. Proc Natl Acad Sci USA. 93, 4409 (1996).

149. B. Ghysels, U. Ochsner, U. Möllman, L. Heinisch, M. Vasil, P. Cornelis, S. Matthijs, The Pseudomonas aeruginosa pirA gene encodes a second receptor for ferrienterobactin and synthetic catecholate analogues. FEMS Microbiology Letters. 246, 167–174 (2005).

150. T. R. Merriman, M. E. Merriman, I. L. Lamont, Nucleotide sequence of pvdD, a pyoverdine biosynthetic gene from Pseudomonas aeruginosa: PvdD has similarity to peptide synthetases. J Bacteriol. 177, 252–258 (1995).

151. G. Ganne, K. Brillet, B. Basta, B. Roche, F. Hoegy, V. Gasser, I. J. Schalk, Iron Release from the Siderophore Pyoverdine in Pseudomonas aeruginosa Involves Three New Actors: FpvC, FpvG, and FpvH. ACS Chem. Biol. 12, 1056–1065 (2017).

152. I. L. Lamont, L. W. Martin, Identification and characterization of novel pyoverdine synthesis genes in Pseudomonas aeruginosa. Microbiology. 149 (2003), pp. 833–842.

153. M. A. Llamas, M. Sparrius, R. Kloet, C. R. Jiménez, C. Vandenbroucke-Grauls, W. Bitter, The heterologous siderophores ferrioxamine B and ferrichrome activate signaling pathways in Pseudomonas aeruginosa. J Bacteriol. 188, 1882–1891 (2006).

154. L. Serino, C. Reimmann, P. Visca, M. Beyeler, V. D. Chiesa, D. Haas, Biosynthesis of pyochelin and dihydroaeruginoic acid requires the iron-regulated pchDCBA operon in Pseudomonas aeruginosa. J. Bacteriol. 179, 248 (1997).

155. C. Hungerer, B. Troup, U. Römling, D. Jahn, Regulation of the hemA gene during 5-aminolevulinic acid formation in Pseudomonas aeruginosa. Journal of bacteriology. 177, 1435–1443 (1995).

156. A. P. Kaur, I. B. Lansky, A. Wilks, The role of the cytoplasmic heme-binding protein (PhuS) of Pseudomonas aeruginosa in intracellular heme trafficking and iron homeostasis. J Biol Chem. 284, 56–66 (2009).

157. A. Rompf, C. Hungerer, T. Hoffmann, M. Lindenmeyer, U. Römling, U. Groß, M. O. Doss, H. Arai, Y. Igarashi, D. Jahn, Regulation of Pseudomonas aeruginosa hemF and hemN by the dual action of the redox response regulators Anr and Dnr. Molecular Microbiology. 29, 985–997 (1998).

158. F. Imperi, F. Tiburzi, P. Visca, Molecular basis of pyoverdine siderophore recycling in *Pseudomonas aeruginosa*. Proc Natl Acad Sci USA. 106, 20440 (2009).

159. K. Brillet, F. Ruffenach, H. Adams, L. Journet, V. Gasser, F. Hoegy, L. Guillon, M. Hannauer, A. Page, I. J. Schalk, An ABC Transporter with Two Periplasmic Binding Proteins Involved in Iron Acquisition in Pseudomonas aeruginosa. ACS Chem. Biol. 7, 2036–2045 (2012).

160. H. E. Cunliffe, T. R. Merriman, I. L. Lamont, Cloning and characterization of pvdS, a gene required for pyoverdine synthesis in Pseudomonas aeruginosa: PvdS is probably an alternative sigma factor. J Bacteriol. 177, 2744–2750 (1995).

161. D. Balasubramanian, H. Kumari, M. Jaric, M. Fernandez, K. H. Turner, S. L. Dove, G. Narasimhan, S. Lory, K. Mathee, Deep sequencing analyses expands the Pseudomonas aeruginosa AmpR regulon to include small RNA-mediated regulation of iron acquisition, heat shock and oxidative stress response. Nucleic Acids Res. 42, 979–998 (2014).

162. E. J. Drake, A. M. Gulick, Structural Characterization and High-Throughput Screening of Inhibitors of PvdQ, an NTN Hydrolase Involved in Pyoverdine Synthesis. ACS Chem. Biol. 6, 1277–1286 (2011).

163. G. Rampioni, M. Falcone, S. Heeb, E. Frangipani, M. P. Fletcher, J.-F. Dubern, P. Visca, L. Leoni, M. Cámara, P. Williams, Unravelling the Genome-Wide Contributions of Specific 2-Alkyl-4-Quinolones and PqsE to Quorum Sensing in Pseudomonas aeruginosa. PLOS Pathogens. 12, e1006029 (2016).

164. J. J. Huang, A. Petersen, M. Whiteley, J. R. Leadbetter, Identification of QuiP, the Product of Gene PA1032, as the Second Acyl-Homoserine Lactone Acylase of *Pseudomonas aeruginosa* PAO1. Appl. Environ. Microbiol. 72, 1190 (2006).

165. W. T. K. Kao, P. M. Gagnon, J. P. Vogel, R. A. Chole, FleQ, a Transcriptional Activator, Is Required for Biofilm Formation In Vitro But Does Not Alter Virulence in a Cholesteatomas Model. Otol Neurotol. 37, 977–983 (2016).

166. A. K. Marr, Joerg Overhage, Manjeet Bains, R. E. W. Hancock, The Lon protease of Pseudomonas aeruginosa is induced by aminoglycosides and is involved in biofilm formation and motility. Microbiology. 153 (2007), pp. 474–482.

167. A. G. Oglesby, J. M. Farrow, J.-H. Lee, A. P. Tomaras, E. P. Greenberg, E. C. Pesci, M. L. Vasil, The Influence of Iron on Pseudomonas aeruginosa Physiology: A REGULATORY LINK BETWEEN IRON AND QUORUM SENSING. Journal of Biological Chemistry. 283, 15558–15567 (2008).

168. B. A. McGuffie, I. Vallet-Gely, S. L. Dove, σ Factor and Anti-σ Factor That Control Swarming Motility and Biofilm Formation in *Pseudomonas aeruginosa*. J. Bacteriol. 198, 755 (2016).

169. L. S. Marmont, J. D. Rich, J. C. Whitney, G. B. Whitfield, H. Almblad, H. Robinson, M. R. Parsek, J. J. Harrison, P. L. Howell, Oligomeric lipoprotein PelC guides Pel polysaccharide export across the outer membrane of *Pseudomonas aeruginosa*. Proc Natl Acad Sci USA. 114, 2892 (2017).

170. K. M. Colvin, N. Alnabelseya, P. Baker, J. C. Whitney, P. L. Howell, M. R. Parsek, PelA deacetylase activity is required for Pel polysaccharide synthesis in Pseudomonas aeruginosa. J Bacteriol. 195, 2329–2339 (2013).

171. F. H. Damron, M. Barbier, E. S. McKenney, M. J. Schurr, J. B. Goldberg, Genes required for and effects of alginate overproduction induced by growth of Pseudomonas aeruginosa on Pseudomonas isolation agar supplemented with ammonium metavanadate. J Bacteriol. 195, 4020–4036 (2013).

172. H. Kulesekara, V. Lee, A. Brencic, N. Liberati, J. Urbach, S. Miyata, D. G. Lee, A. N. Neely, M. Hyodo, Y. Hayakawa, F. M. Ausubel, S. Lory, Analysis of Pseudomonas aeruginosa diguanylate cyclases and phosphodiesterases reveals a role for bis-(3′-5′)-cyclic-GMP in virulence. Proceedings of the National Academy of Sciences of the United States of America. 103, 2839–2844 (2006).

173. V. Deretic, J. F. Gill, A. M. Chakrabarty, Gene algD coding for GDPmannose dehydrogenase is transcriptionally activated in mucoid Pseudomonas aeruginosa. J. Bacteriol. 169, 351 (1987).

174. S.-K. Wang, I. SA’-CORREIA, A. Darzins, A. M. Chakrabarty, Characterization of the Pseudomonas aeruginosa Alginate (alg) Gene Region II. Microbiology. 133 (1987), pp. 2303–2314.

175. Y. Irie, M. Starkey, A. N. Edwards, D. J. Wozniak, T. Romeo, M. R. Parsek, Pseudomonas aeruginosa biofilm matrix polysaccharide Psl is regulated transcriptionally by RpoS and post-transcriptionally by RsmA. Mol Microbiol. 78, 158–172 (2010).

176. S. Ruer, S. Stender, A. Filloux, S. de Bentzmann, Assembly of fimbrial structures in Pseudomonas aeruginosa: functionality and specificity of chaperone-usher machineries. J Bacteriol. 189, 3547–3555 (2007).

177. O. E. Petrova, K. Sauer, A novel signaling network essential for regulating Pseudomonas aeruginosa biofilm development. PLoS Pathog. 5, e1000668–e1000668 (2009).

178. S. de Bentzmann, M. Aurouze, G. Ball, A. Filloux, FppA, a Novel *Pseudomonas aeruginosa* Prepilin Peptidase Involved in Assembly of Type IVb Pili. J. Bacteriol. 188, 4851 (2006).

179. P. Chiang, L. L. Burrows, Biofilm formation by hyperpiliated mutants of Pseudomonas aeruginosa. J Bacteriol. 185, 2374–2378 (2003).

180. S. L. N. Kilmury, L. L. Burrows, The *Pseudomonas aeruginosa* PilSR Two-Component System Regulates Both Twitching and Swimming Motilities. mBio. 9, e01310–18 (2018).

181. V. Deretic, W. M. Konyecsni, A procaryotic regulatory factor with a histone H1-like carboxy-terminal domain: clonal variation of repeats within algP, a gene involved in regulation of mucoidy in Pseudomonas aeruginosa. J Bacteriol. 172, 5544–5554 (1990).

182. Y. Luo, K. Zhao, A. E. Baker, S. L. Kuchma, K. A. Coggan, M. C. Wolfgang, G. C. L. Wong, G. A. O’Toole, A Hierarchical Cascade of Second Messengers Regulates Pseudomonas aeruginosa Surface Behaviors. mBio. 6 (2015), doi:10.1128/mBio.02456-14.

183. B. M. Babin, M. Bergkessel, M. J. Sweredoski, A. Moradian, S. Hess, D. K. Newman, D. Tirrell, SutA is a bacterial transcription factor expressed during slow growth in *Pseudomonas aeruginosa*. Proc Natl Acad Sci USA. 113, E597 (2016).

184. S. A. Reiling, J. A. Jansen, B. J. Henley, S. Singh, C. Chattin, M. Chandler, D. W. Rowen, Prc protease promotes mucoidy in mucA mutants of Pseudomonas aeruginosa. Microbiology. 151 (2005), pp. 2251–2261.

185. C. B. Whitchurch, A. J. Leech, M. D. Young, D. Kennedy, J. L. Sargent, J. J. Bertrand, A. B. T. Semmler, A. S. Mellick, P. R. Martin, R. A. Alm, M. Hobbs, S. A. Beatson, B. Huang, L. Nguyen, J. C. Commolli, J. N. Engel, A. Darzins, J. S. Mattick, Characterization of a complex chemosensory signal transduction system which controls twitching motility in Pseudomonas aeruginosa. Mol Microbiol. 52, 873–893 (2004).

186. M. T. Cabeen, S. A. Leiman, R. Losick, Colony-morphology screening uncovers a role for the Pseudomonas aeruginosa nitrogen-related phosphotransferase system in biofilm formation. Mol Microbiol. 99, 557–570 (2016).

187. A. L. Goodman, B. Kulasekara, A. Rietsch, D. Boyd, R. S. Smith, S. Lory, A Signaling Network Reciprocally Regulates Genes Associated with Acute Infection and Chronic Persistence in Pseudomonas aeruginosa. Developmental Cell. 7, 745–754 (2004).

188. C. R. Armbruster, C. K. Lee, J. Parker-Gilham, J. de Anda, A. Xia, K. Zhao, K. Murakami, B. S. Tseng, L. R. Hoffman, F. Jin, C. S. Harwood, G. C. Wong, M. R. Parsek, Heterogeneity in surface sensing suggests a division of labor in Pseudomonas aeruginosa populations. Elife. 8, e45084 (2019).

189. Y. Zhang, B. Xia, M. Li, J. Shi, Y. Long, Y. Jin, F. Bai, Z. Cheng, S. Jin, W. Wu, HigB Reciprocally Controls Biofilm Formation and the Expression of Type III Secretion System Genes through Influencing the Intracellular c-di-GMP Level in Pseudomonas aeruginosa. Toxins (Basel). 10, 424 (2018).

190. W.-K. Choy, L. Zhou, C. K.-C. Syn, L.-H. Zhang, S. Swarup, MorA defines a new class of regulators affecting flagellar development and biofilm formation in diverse Pseudomonas species. J Bacteriol. 186, 7221–7228 (2004).

191. B. R. Borlee, A. D. Goldman, K. Murakami, R. Samudrala, D. J. Wozniak, M. R. Parsek, Pseudomonas aeruginosa uses a cyclic-di-GMP-regulated adhesin to reinforce the biofilm extracellular matrix. Molecular Microbiology. 75, 827–842 (2010).

192. C. Giraud, C. Bernard, S. Ruer, S. De Bentzmann, Biological ‘glue’ and ‘Velcro’: molecular tools for adhesion and biofilm formation in the hairy and gluey bug Pseudomonas aeruginosa. Environmental Microbiology Reports. 2, 343–358 (2010).

193. P. Tielen, M. Strathmann, K.-E. Jaeger, H.-C. Flemming, J. Wingender, Alginate acetylation influences initial surface colonization by mucoid Pseudomonas aeruginosa. Microbiological Research. 160, 165–176 (2005).

194. D. E. Nivens, D. E. Ohman, J. Williams, M. J. Franklin, Role of alginate and its O acetylation in formation of Pseudomonas aeruginosa microcolonies and biofilms. J Bacteriol. 183, 1047–1057 (2001).

195. A. Darzins, B. Frantz, R. I. Vanags, A. M. Chakrabarty, Nucleotide sequence analysis of the phosphomannose isomerase gene (pmi) of Pseudomonas aeruginosa and comparison with the corresponding Escherichia coli gene manA. Gene. 42, 293–302 (1986).

196. M. Hentzer, G. M. Teitzel, G. J. Balzer, A. Heydorn, S. Molin, M. Givskov, M. R. Parsek, Alginate overproduction affects Pseudomonas aeruginosa biofilm structure and function. J Bacteriol. 183, 5395–5401 (2001).

197. P. N. Jimenez, G. Koch, E. Papaioannou, M. Wahjudi, J. Krzeslak, T. Coenye, R. H. Cool, W. J. Quax, Role of PvdQ in Pseudomonas aeruginosa virulence under iron-limiting conditions. Microbiology. 156 (2010), pp. 49–59.

198. K. Murphy, A. J. Park, Y. Hao, D. Brewer, J. S. Lam, C. M. Khursigara, Influence of O Polysaccharides on Biofilm Development and Outer Membrane Vesicle Biogenesis in *Pseudomonas aeruginosa* PAO1. J. Bacteriol. 196, 1306 (2014).

199. X. Z. Li, H. Nikaido, K. Poole, Role of mexA-mexB-oprM in antibiotic efflux in Pseudomonas aeruginosa. Antimicrob Agents Chemother. 39, 1948–1953 (1995).

200. H. Li, Y.-F. Luo, B. J. Williams, T. S. Blackwell, C.-M. Xie, Structure and function of OprD protein in Pseudomonas aeruginosa: from antibiotic resistance to novel therapies. Int J Med Microbiol. 302, 63–68 (2012).

201. M. Castanheira, S. E. Farrell, K. M. Krause, R. N. Jones, H. S. Sader, Contemporary diversity of β-lactamases among Enterobacteriaceae in the nine U.S. census regions and ceftazidime-avibactam activity tested against isolates producing the most prevalent β-lactamase groups. Antimicrob Agents Chemother. 58, 833–838 (2014).

202. E. B. M. Breidenstein, L. Janot, J. Strehmel, L. Fernandez, P. K. Taylor, I. Kukavica-Ibrulj, S. L. Gellatly, R. C. Levesque, J. Overhage, R. E. W. Hancock, The Lon Protease Is Essential for Full Virulence in Pseudomonas aeruginosa. PLOS ONE. 7, e49123 (2012).

203. Y. Morita, J. Tomida, Y. Kawamura, Responses of Pseudomonas aeruginosa to antimicrobials. Frontiers in Microbiology. 4, 422 (2014).

204. E. Z. Baum, S. M. Crespo-Carbone, B. J. Morrow, T. A. Davies, B. D. Foleno, W. He, A. M. Queenan, K. Bush, Effect of MexXY overexpression on ceftobiprole susceptibility in Pseudomonas aeruginosa. Antimicrobial agents and chemotherapy. 53, 2785–2790 (2009).

205. L. A. Gallagher, J. Shendure, C. Manoil, Genome-Scale Identification of Resistance Functions in *Pseudomonas aeruginosa* Using Tn-seq. mBio. 2, e00315–10 (2011).

206. G. McCaughey, D. F. Gilpin, T. Schneiders, L. R. Hoffman, M. McKevitt, J. S. Elborn, M. M. Tunney, Fosfomycin and Tobramycin in Combination Downregulate Nitrate Reductase Genes *narG* and *narH*, Resulting in Increased Activity against *Pseudomonas aeruginosa* under Anaerobic Conditions. Antimicrob. Agents Chemother. 57, 5406 (2013).

207. R. P. Lamers, U. T. Nguyen, Y. Nguyen, R. N. C. Buensuceso, L. L. Burrows, Loss of membrane-bound lytic transglycosylases increases outer membrane permeability and β-lactam sensitivity in Pseudomonas aeruginosa. MicrobiologyOpen. 4, 879–895 (2015).

208. Y. Li, T. Mima, Y. Komori, Y. Morita, T. Kuroda, T. Mizushima, T. Tsuchiya, A new member of the tripartite multidrug efflux pumps, MexVW–OprM, in Pseudomonas aeruginosa. Journal of Antimicrobial Chemotherapy. 52, 572–575 (2003).

209. L. Zamorano, T. M. Reeve, C. Juan, B. Moyá, G. Cabot, D. J. Vocadlo, B. L. Mark, A. Oliver, AmpG Inactivation Restores Susceptibility of Pan-β-lactam Resistant *Pseudomonas aeruginosa* Clinical Strains. Antimicrob. Agents Chemother. (2011), doi:10.1128/AAC.01688-10.

210. L. Moynié, A. Luscher, D. Rolo, D. Pletzer, A. Tortajada, H. Weingart, Y. Braun, M. G. P. Page, J. H. Naismith, T. Köhler, Structure and Function of the PiuA and PirA Siderophore-Drug Receptors from Pseudomonas aeruginosa and Acinetobacter baumannii. Antimicrob Agents Chemother. 61, e02531–16 (2017).

211. R. Srikumar, T. Kon, N. Gotoh, K. Poole, Expression of *Pseudomonas aeruginosa* Multidrug Efflux Pumps MexA-MexB-OprM and MexC-MexD-OprJ in a Multidrug-Sensitive *Escherichia coli* Strain. Antimicrob. Agents Chemother. 42, 65 (1998).

212. M.-L. Han, T. Velkov, Y. Zhu, K. D. Roberts, A. P. Le Brun, S. H. Chow, A. D. Gutu, S. M. Moskowitz, H.-H. Shen, J. Li, Polymyxin-Induced Lipid A Deacylation in Pseudomonas aeruginosa Perturbs Polymyxin Penetration and Confers High-Level Resistance. ACS Chem. Biol. 13, 121–130 (2018).

213. S. Rocchio, D. Santorelli, S. Rinaldo, M. Franceschini, F. Malatesta, F. Imperi, L. Federici, C. Travaglini-Allocatelli, A. Di Matteo, Structural and functional investigation of the Small Ribosomal Subunit Biogenesis GTPase A (RsgA) from Pseudomonas aeruginosa. The FEBS Journal. 286, 4245–4260 (2019).

214. M. S. Sonnabend, K. Klein, S. Beier, A. Angelov, R. Kluj, C. Mayer, C. Groß, K. Hofmeister, A. Beuttner, M. Willmann, S. Peter, P. Oberhettinger, A. Schmidt, I. B. Autenrieth, M. Schütz, E. Bohn, Identification of Drug Resistance Determinants in a Clinical Isolate of Pseudomonas aeruginosa by High-Density Transposon Mutagenesis. Antimicrob Agents Chemother. 64, e01771–19 (2020).

215. G. Dieppois, V. Ducret, O. Caille, K. Perron, The Transcriptional Regulator CzcR Modulates Antibiotic Resistance and Quorum Sensing in Pseudomonas aeruginosa. PLOS ONE. 7, e38148 (2012).

216. T. Mima, N. Kohira, Y. Li, H. Sekiya, W. Ogawa, T. Kuroda, T. Tsuchiya, Gene cloning and characteristics of the RND-type multidrug efflux pump MuxABC-OpmB possessing two RND components in Pseudomonas aeruginosa. Microbiology. 155 (2009), pp. 3509– 3517.

217. E. A. Rundell, N. Commodore, A. L. Goodman, B. I. Kazmierczak, A Screen for Antibiotic Resistance Determinants Reveals a Fitness Cost of the Flagellum in *Pseudomonas aeruginosa*. J. Bacteriol. 202, e00682–19 (2020).

218. K. Jeannot, M. L. Sobel, F. El Garch, K. Poole, P. Plésiat, Induction of the MexXY Efflux Pump in *Pseudomonas aeruginosa* Is Dependent on Drug-Ribosome Interaction. J. Bacteriol. 187, 5341 (2005).

219. J. Handfield, L. Gagnon, M. Dargis, A. Huletsky, Sequence of the ponA gene and characterization of the penicillin-binding protein 1A of Pseudomonas aeruginosa PAO1. Gene. 199, 49–56 (1997).

220. A. Ropy, G. Cabot, I. Sánchez-Diener, C. Aguilera, B. Moya, J. A. Ayala, A. Oliver, Role of *Pseudomonas aeruginosa* Low-Molecular-Mass Penicillin-Binding Proteins in AmpC Expression, β-Lactam Resistance, and Peptidoglycan Structure. Antimicrob. Agents Chemother. 59, 3925 (2015).

221. Y. Heacock-Kang, Z. Sun, J. Zarzycki-Siek, K. Poonsuk, I. A. McMillan, R. Chuanchuen, T. T. Hoang, Two Regulators, PA3898 and PA2100, Modulate the *Pseudomonas aeruginosa* Multidrug Resistance MexAB-OprM and EmrAB Efflux Pumps and Biofilm Formation. Antimicrob. Agents Chemother. 62, e01459–18 (2018).

222. A. Dötsch, T. Becker, C. Pommerenke, Z. Magnowska, L. Jänsch, S. Häussler, Genomewide Identification of Genetic Determinants of Antimicrobial Drug Resistance in *Pseudomonas aeruginosa*. Antimicrob. Agents Chemother. 53, 2522 (2009).

223. C. Alvarez-Ortega, I. Wiegand, J. Olivares, R. E. W. Hancock, J. L. Martínez, Genetic determinants involved in the susceptibility of Pseudomonas aeruginosa to beta-lactam antibiotics. Antimicrob Agents Chemother. 54, 4159–4167 (2010).

224. A. Wong, N. Rodrigue, R. Kassen, Genomics of Adaptation during Experimental Evolution of the Opportunistic Pathogen Pseudomonas aeruginosa. PLOS Genetics. 8, e1002928 (2012).

225. H. H. Mehta, A. G. Prater, K. Beabout, R. A. L. Elworth, M. Karavis, H. S. Gibbons, Y. Shamoo, The Essential Role of Hypermutation in Rapid Adaptation to Antibiotic Stress. Antimicrob Agents Chemother. 63, e00744–19 (2019).

226. S. J. T. Wardell, A. Rehman, L. W. Martin, C. Winstanley, W. M. Patrick, I. L. Lamont, Antimicrob. Agents Chemother., in press, doi:10.1128/AAC.01619-19.

227. I. Wiegand, A. K. Marr, E. B. M. Breidenstein, K. N. Schurek, P. Taylor, R. E. W. Hancock, Mutator Genes Giving Rise to Decreased Antibiotic Susceptibility in *Pseudomonas aeruginosa*. Antimicrob. Agents Chemother. 52, 3810 (2008).

228. G. Cabot, L. Zamorano, B. Moyà, C. Juan, A. Navas, J. Blázquez, A. Oliver, Evolution of *Pseudomonas aeruginosa* Antimicrobial Resistance and Fitness under Low and High Mutation Rates. Antimicrob. Agents Chemother. 60, 1767 (2016).

229. A. Hinz, S. Lee, K. Jacoby, C. Manoil, Membrane proteases and aminoglycoside antibiotic resistance. J Bacteriol. 193, 4790–4797 (2011).

230. F. El’Garch, K. Jeannot, D. Hocquet, C. Llanes-Barakat, P. Plésiat, Cumulative effects of several nonenzymatic mechanisms on the resistance of Pseudomonas aeruginosa to aminoglycosides. Antimicrob Agents Chemother. 51, 1016–1021 (2007).

